# Engineered protein circuits for cancer therapy

**DOI:** 10.1101/2025.04.16.647665

**Authors:** Andrew C. Lu, Lukas Moeller, Stephen Moore, Kevin Ho, Victoria Tobin, Albert Qiang, Evan Zhang, Zhijie Li, Mengmeng Wang, Shiyu Xia, Mark W. Budde, Haley Larson, Ali Ahmed Diaz, Bo Gu, James M. Linton, Leslie Klock, Michael J. Flynn, Queen Igomu, Xiaojing J. Gao, Daniel J. Siegwart, Hao Zhu, Michael B. Elowitz

## Abstract

A central challenge in cancer therapy is selectively killing cancer cells while minimizing resistance. Here, we engineer modular protease-based protein circuits that sense mutant RAS, the most frequently mutated oncogene in cancer, and conditionally activate cell death. Delivered transiently as mRNA in lipid nanoparticles (LNPs), circuits selectively eliminated RAS-mutant cancer cells in culture and suppressed aggressive, multifocal RAS-driven liver tumors. Compared to RAS inhibitors, circuits killed cancer cells independently of oncogene addiction, achieved potent cytotoxicity rather than cytostasis, and functioned at low RAS occupancy through catalysis rather than stoichiometric inhibition. Critically, circuits acquired minimal to no resistance under prolonged selection, while remaining potent against prevalent drug-resistance mechanisms including RAS amplification and bypass signaling. Together, these results establish design principles for engineering therapeutic protein circuits and highlight their potential to overcome longstanding limitations of existing therapeutic modalities.

---

Oncogenes play a dual role in cancer as drivers of proliferation and therapeutic targets. In many cases, cancer cell survival depends on continued oncogenic signaling, a phenomenon known as oncogenic addiction ^1,2^. This dependency led to the development of targeted inhibitors that suppress cancer by inhibiting specific oncogenes ^3,4^. Although they have transformed patient care ^4,5^, inhibitors also have fundamental limitations. Their mechanism for cancer cell elimination is indirect, depending on pathways downstream of the oncogene itself to translate inhibition into therapeutic outcomes ^6,7^. They are highly susceptible to resistance, which ultimately limits their durability ^8,9^. And they require oncogene addiction, which is incomplete or missing in many cancers ^10,11^. These observations provoke the question of whether and how it might be possible to target oncogenes in different ways that avoid or minimize these limitations.

One inspiration comes from the natural p53 tumor suppressor pathway, which senses oncogenic stress and directly responds by inducing cell cycle arrest, DNA repair, or cell death ^12,13^. Advances in protein engineering, synthetic biology, and mRNA delivery now make it possible to design analogous synthetic sense-and-response protein circuits. These circuits comprise engineered proteins, delivered to tumor and normal cells, that sense disease-specific signals and conditionally execute cell death or other responses in tumor cells (Fig. 1A). While aspects of this paradigm have been explored ^14–28^, key questions remain unclear: What design principles enable engineering of sensitive and specific therapeutic circuits? Can systemically delivered circuits suppress cancer in vivo? And most fundamentally, how do circuits differ from inhibitors with respect to resistance susceptibility and other critical features?

Here, we introduce modular, programmable therapeutic circuits that selectively kill cancer cells harboring mutant RAS, the most commonly mutated oncogene ^29–31^. We engineer protease-based sensors that detect a broad spectrum of clinically relevant RAS mutations and couple them to engineered effectors of apoptotic or pyroptotic cell death. Delivered transiently as mRNA in lipid nanoparticles (LNPs), these circuits selectively eliminated RAS-mutant cancer cells in culture and suppressed aggressive RAS-driven liver tumors in mice. Head-to-head comparisons with the RAS inhibitors Sotorasib and RMC-7977 (tool compound for Daraxonrasib) demonstrated key advantages of circuits: they kill independently of RAS addiction, are potently cytotoxic, act at low RAS occupancy, decouple killing from broad pathway suppression, and are less perturbative to cells with wild-type RAS signaling. Most strikingly, therapeutic circuits are less susceptible to acquired resistance, and remain active against prevalent drug resistance mechanisms such as RAS amplification, *cis* mutations, and bypass signaling pathways. These results establish therapeutic protein circuits as a new modality that complements and expands the existing arsenal of cancer therapies.

## Designing programmable mutant-RAS sensors

Previous work introduced a RAS-sensing architecture based on split protease reconstitution ^15,18,32,33^. The RAS binding domain from RAF1 was fused to complementary halves of a split Tobacco Etch Virus (TEV) protease, whose cleavage site is absent from the human proteome and thus minimizes off-target proteolysis (**fig. S1**). RAS activation then led to co-recruitment to active RAS clusters on the plasma membrane and reconstitution of the split protease (**Fig. 1A**, *“Sensor”*). This sensor (henceforth denoted sensor_v1_) responded well to overexpressed RAS, but was not sensitive enough to detect mutant RAS (**fig. S2**).

**Fig. 1.**
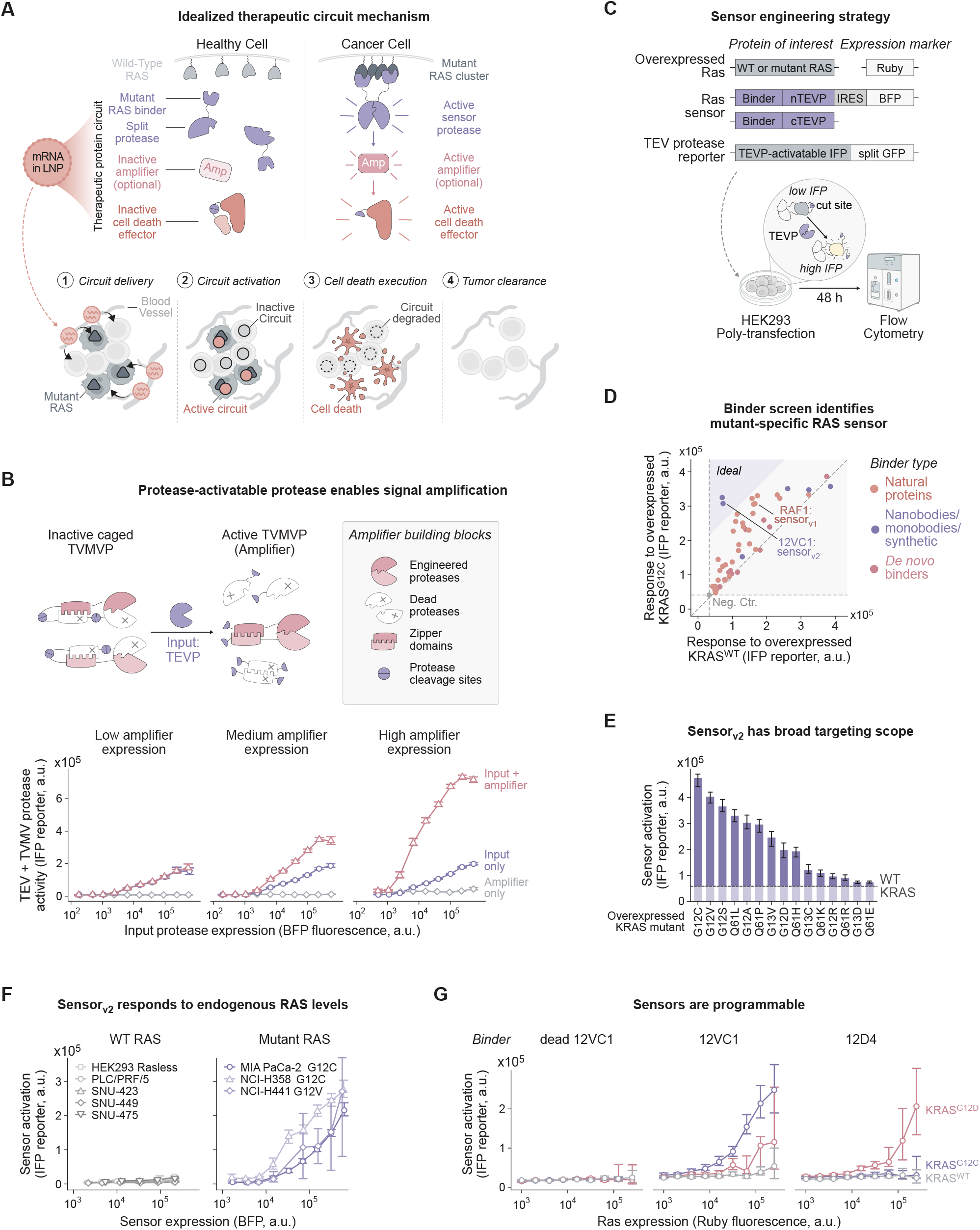
Engineering of mutant-specific RAS sensors. **(A)** Schematic of idealized therapeutic circuit mechanism. Top: Therapeutic protein circuits composed of a mutant-RAS binder fused to complementary split protease halves, an optional protease amplifier, and a cell-death effector are delivered as mRNA in lipid nanoparticles (LNPs). In cancer cells, mutant RAS on the plasma membrane co-recruits the split protease sensor halves, leading to protease reconstitution and activation of downstream effectors, triggering cell death. Bottom: (1) circuit delivery, (2) circuit activation, (3) cell-death execution, (4) tumor clearance. **(B)** Protease-activatable proteases enable signal amplification. Top: dual-caged TVMV protease (TVMVP) amplifier design. Bottom: median TEV+TVMV protease activity (IFP reporter) versus input protease expression (BFP). **(C)** Sensor engineering strategy. HEK293 cells were poly-transfected with wild-type or mutant RAS (expression marked by Ruby), candidate sensors (BFP), and a TEVP-activatable IFP reporter (GFP), then analyzed by flow cytometry. **(D)** Binder screen identifies mutant-specific RAS sensor. Data points represent median reporter activation (IFP reporter) in response to overexpressed KRAS^G12C^ (y-axis) versus KRAS^WT^ (x-axis) at high RAS expression. **(E)** The 12VC1 sensor (sensor_v2_) responds to diverse oncogenic KRAS mutations. Bars show median sensor activation (IFP reporter) for each overexpressed KRAS mutant; dashed line indicates response to KRAS^WT^. **(F)** The 12VC1 sensor detects endogenous mutant RAS levels. Median sensor activation (IFP reporter) as a function of sensor expression (BFP) is shown. **(G)** Replacing binders enables reprogramming of sensor targeting scope. Median sensor activation (IFP reporter) as a function of RAS expression (Ruby) for a sensor with a dead 12VC1 binder (negative control), the 12VC1 sensor, and the 12D4 sensor. Error bars and shaded error regions in **(B)** and **(E)-(G)** represent 95% confidence intervals of the median calculated by bootstrapping.

To increase sensitivity, inspired by amplifying proteolytic cascades in cell death pathways ^34,35^, we engineered a protease-activated protease amplification module, in which a tobacco vein mottling virus (TVMV) protease is proteolytically activated by the sensor TEV protease ^20,36^. Most designs provided little amplification, or even de-amplification (**fig. S3**). The exception was a dual-caged design, in which the TVMV protease is split, and each half is fused to a catalytically dead mutant of its complementary half, functioning as a caging domain (**Fig. 1B**, *see **Methods** for design details*). Engineered cleavage sites allow TEV protease to detach the functional halves from the caging domains. Further, each functional TVMV protease half was fused to a coiled-coil dimerization domain ^15,20,37^, whose strong affinity for its complementary domain drives competitive displacement of the caging domains. Thus, input TEV protease activity allows reconstitution of active TVMV protease. When expressed, this amplifier successfully increased sensitivity with minimal background, and proved useful in several experiments below (**Fig. 1B**).

While effective, the amplifier added considerable circuit complexity. As a simpler, complementary approach, we screened for alternative RAS binding domains that could provide greater sensitivity and mutant-specificity than the RAF1 domain. We compiled a list of natural, *de novo* designed, and engineered RAS binding domains (**Methods**) ^38–43^. Each candidate domain was fused to complementary split TEV protease domains to create a sensor. Finally, we polytransfected ^44^ each sensor together with mutant or wild-type RAS and fluorescent protease reporters to quantitatively explore sensor responses across a broad expression space in HEK293 cells (**Fig. 1C**, **fig. S4**, **Methods**). Among these, the 12VC1 monobody ^39^ achieved the greatest mutant-specificity (henceforth denoted sensor_v2_; **Fig. 1D**, **fig. S5 and S6**). This likely reflects its original identification through positive selection for KRAS^G12C^-binding and negative selection against binding to wild-type KRAS. We confirmed that KRAS inactivation by Sotorasib and the dominant-negative KRAS S17N mutation ^45^ abrogated sensor activity, confirming RAS dependence (**fig. S7**).

Sensor_v2_ provided remarkable sensitivity and specificity. It responded to diverse clinically relevant mutations in all three RAS isoforms (**Fig. 1E**, **fig. S8**; *see **Methods** section “Mutant-specificity of the 12VC1-based sensor”*), and accurately discriminated between mutant and wild-type RAS at the single-cell level (**fig. S10**). Most importantly, it sensitively responded to physiological expression levels of RAS mutants in the cancer cell lines MIA PaCa-2 (KRAS^G12C^), NCI-H358 (KRAS^G12C^), and NCI-H441 (KRAS^G12V^; **Fig. 1F**). By contrast, it exhibited minimal response in cancer cell lines with wild-type RAS, including HEK293, PLC/PRF/5, SNU-423, SNU-449, and SNU-475 (**Fig. 1F**).

To gain insight into biophysical requirements for sensor activation, we analyzed the response of the sensor_v2_ to a set of KRAS mutations affecting different protein functions (**fig. S11**). Disrupting the KRAS dimer or clustering interface had little effect on sensor sensitivity ^46–48^. In contrast, deleting or weakening the membrane-targeting domains (the CAAX motif and polybasic domain) abolished or reduced sensor activity ^49–51^. These results suggest that RAS membrane localization, but not clustering, is required for sensor activation.

We also added two additional modifications to enhance sensor performance. First, inspired by natural self-cleaving viral polyproteins ^52^, we concatenated the two sensor components into a single polyprotein (sensor_v3_), whose TEV protease products post-translationally cleave themselves into distinct proteins (**fig. S12**). This design increased maximal activation. Second, we incorporated mutations known to enhance TEV protease catalytic activity ^53^, which further improved sensitivity (**fig. S13**). This final design (sensor_v4_) is used in all experiments below, unless otherwise noted.

Finally, we asked whether the modularity of the sensor design enabled reprogramming of the sensor’s mutant selectivity. We replaced 12VC1 with the previously described KRAS^G12D^-specific monobody 12D4^41^. This sensor responded strongly to G12D but not wild-type KRAS (**Fig. 1G**), demonstrating that sensor specificity can be effectively redirected by swapping binding domains.

Taken together, these results revealed that split protein reconstitution, mutant-specific binding, and competitive displacement of caging domains are sufficient to create sensors and amplifiers capable of detecting and discriminating physiological levels of mutant RAS in human cells.

### Circuits couple RAS sensing to cell death

With a mutant-specific sensor in hand, we next sought to develop full therapeutic circuits that selectively kill cancer cells when delivered as mRNA in LNPs. First, as a cell death effector, we adapted a previously engineered caspase-3 whose natural activating proteolytic cleavage site ^18,54,55^ was replaced by the TEV protease cleavage site ^18^. We further incorporated a membrane localization tag to colocalize the effector (henceforth denoted as casp) with the reconstituted sensor, bound to RAS clusters on the plasma membrane ^56^. Next, we confirmed that a liver-tropic LNP formulation (based on the lipid 4A3-SC8, **Methods**) can efficiently deliver *in vitro* transcribed mRNA to cells in culture in a dose-dependent fashion ^57,58^ (**fig. S14**, *and see below*).

To evaluate circuit performance, we encapsulated engineered sensor_v4_ and casp mRNAs in separate LNPs, co-delivered them at varying concentrations to human cancer cell lines, and analyzed cell viability 3-5 days post-treatment (**Fig. 2A**). Both the sensor and the effector were well tolerated across a wide dosage regime when delivered individually (**Fig. 2B**). In contrast, co-delivery of the complete circuit, denoted sensor_v4_-casp, induced potent, near-complete, dose-dependent cell death in RAS-mutant cancer cell lines, but not in RAS wild-type cancer cells (**Fig. 2A**). These results demonstrate that LNP-delivered therapeutic circuits can selectively kill human RAS-mutant cancer cells.

**Fig. 2.**
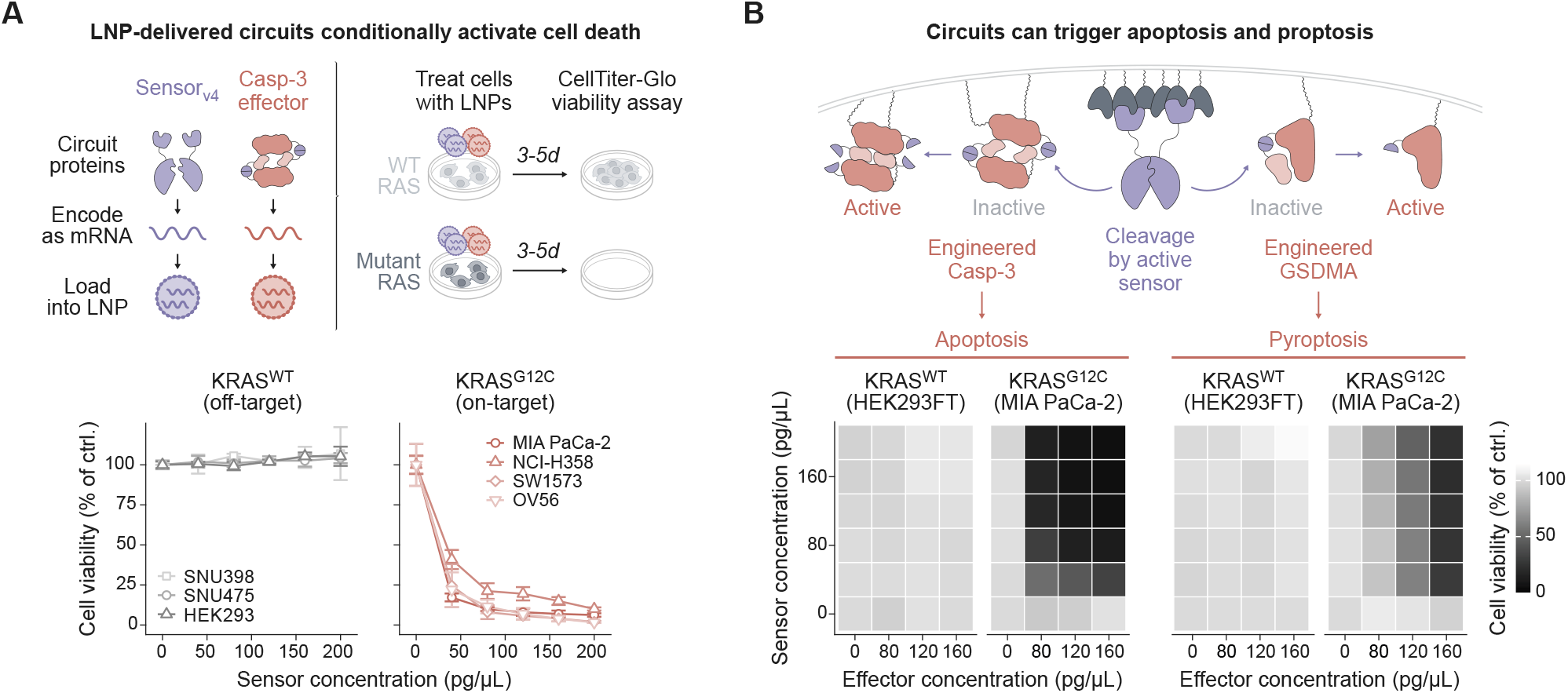
Circuits potently and selectively eliminate RAS-mutant cancer cells. **(A)** LNP-delivered sensor_v4_-casp circuit conditionally eliminates RAS-mutant cancer cells in vitro. RAS-wild-type and mutant cells were treated with mRNA-LNP-delivered circuits for 3-5 days. Normalized mean cell viability ± s.d. is shown (n = 3 replicates). **(B)** Circuits trigger apoptosis or pyroptosis. Heatmaps show normalized mean cell viability (n = 3 replicates).

Whereas apoptosis is confined to the dying cell, pyroptosis is an inflammatory form of cell death whose effects can spread beyond the affected cell ^18,59–61^. Pyroptosis has been shown to achieve tumor clearance even when triggered in a minority of cancer cells ^62,63^, making it an ideal circuit output that could leverage the immune system to compensate for incomplete delivery to tumor cells (**fig. S15**). Pyroptosis is naturally triggered by proteolytic activation of gasdermins (GSDMs) ^59,64,65^. To test conditional activation of pyroptosis, we co-delivered sensor_v4_ with a TEV cleavage-activatable gasdermin (GSDMA; **Fig. 2B**) ^18,62,66^. As above, these components induced negligible background cell death alone, but potently and selectively eliminated RAS-mutant human cancer cells when co-delivered as a complete therapeutic circuit (sensor_v4_-GSDM; **Fig. 2B**, **fig. S16**). Taken together, these results show that therapeutic circuits, delivered as mRNA in LNPs, can effectively target RAS-mutant cancer cells through two complementary cell death programs.

### Circuits prevent RAS-driven liver tumors

We next asked whether circuits could function *in vivo*. The liver represents a site of high unmet need for RAS-driven disease, and hepatocytes can be efficiently transfected with existing LNP formulations ^67–69^. RAS mutations are common in intrahepatic cholangiocarcinoma (in about 15-25%) and dominate cancers that metastasize to the liver ^70^. In particular, about half of colorectal cancers carry RAS mutations ^71^, and up to half of all these patients develop unresectable liver metastases, a leading cause of colorectal cancer morbidity and mortality ^72,73^.

Liver tumors develop in complex microenvironments, with vasculature, immune cells, and extracellular matrix influencing cancer growth ^74,75^. To recapitulate these features, we used an established autochthonous liver cancer model in immunocompetent mice ^76,77^. In this model, hydrodynamic tail vein injection (HDT) enables transfection of hepatocytes with DNA encoding the oncogenic NRAS^G12V^ RAS variant, an shRNA targeting TP53 (shTP53), and the Sleeping Beauty (SB100) transposase for stable genomic integration and sustained expression ^78–80^. The model generates aggressive, multifocal liver cancers that advance to late-stage disease within three to five weeks after induction ^78^.

To determine whether circuits could function *in vivo* independently of potential delivery limitations, we co-administered DNA-encoded circuits together with tumor-inducing agents (**Fig. 3A**). This approach allows tumor initiation and circuit expression to occur in the same liver cells. These early experiments were performed before the optimized sensor_v4_ was developed, and therefore used the initial RAF1-RBD-based sensor_v1_ paired with the Casp3 apoptosis effector (sensor_v1_-casp). To test whether the proteolytic amplifier could increase circuit sensitivity, we performed these experiments with and without amplification. In one mouse cohort, we let tumors develop for five weeks, terminated the experiment, and measured total surface tumor areas and liver-to-body weight ratios, standard measures of tumor burden in liver tumor models (**Fig. 3A**, **Methods**). In a separate cohort, we tracked survival to determine the durability of tumor prevention.

**Fig. 3.**
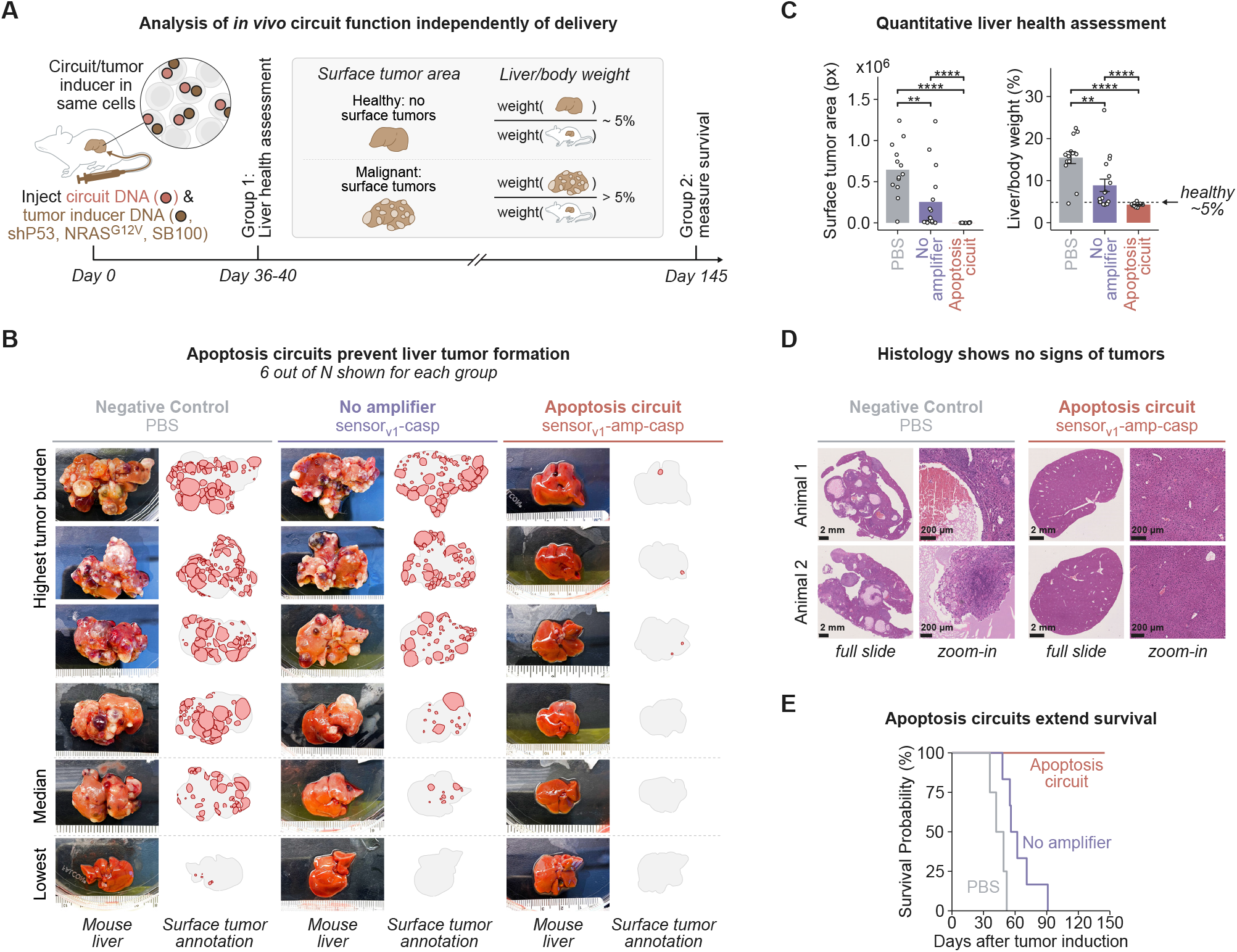
Circuits prevent RAS-driven liver tumors in vivo. **(A)** DNA-encoded circuit and tumor-inducer plasmids (shTP53, NRAS^G12V^, SB100 transposase) were co-administered to mouse hepatocytes via hydrodynamic tail-vein injection (HDT). Animals were sacrificed for liver-health assessment or followed for survival analysis. **(B)** Representative mouse livers and blinded surface-tumor annotations for negative control (PBS), no amplifier (sensor_v1_-casp), and apoptosis circuit (sensor_v1_-amp-casp) are shown. **(C)** Quantification of tumor burden. Data points represent individual animals, bars depict mean ± s.e.; dashed line indicates liver-to-body weight ratio of healthy animals (∼5%). **(D)** H&E staining of liver sections from negative control (PBS) and apoptosis circuit (sensor_v1_-amp-casp) animals. **(E)** Kaplan-Meier survival curves (n = 7-10 mice per condition).

Mice that received no treatment showed robust, multifocal tumor formation (**Fig. 3B**, **fig. S17**). The sensor_v1_-casp circuit significantly, but incompletely, reduced tumor burden, as assessed by total surface tumor area (p = 4.1 × 10^-3^) and liver-to-body weight ratios (p = 3.0 × 10^-3^; **Fig. 3, B and C**, **fig. S17 and S18**, **table S1**). The same treatment extended median survival from 42 days (in untreated animals) to 59 days (p = 7 × 10^-4^; **Fig. 3E**). Strikingly, however, inclusion of the amplifier module (sensor_v1_-amp-casp) eliminated detectable surface tumors (p = 3.2 × 10^-6^) and restored liver-to-body weight ratios to those of healthy animals (p = 3.5 × 10^-7^; **Fig. 3, B and C**, **fig. S17 and S18**, **table S1**). Absence of tumors was also confirmed by histology (**Fig. 3D**). Further, in the parallel cohort, all mice treated with this sensor_v1_-amp-casp circuit survived to the 145 day endpoint (p = 1 × 10^-4^ for comparison to untreated, see Methods; **Fig. 3E**). Overall, these results demonstrate that co-delivered therapeutic circuits can prevent tumor formation *in vivo*.

### Systemically-delivered circuits suppress liver tumors

While DNA-encoded circuits can prevent tumor formation by co-delivered oncogenic constructs, a more clinically relevant question is whether systemically delivered and mRNA-encoded circuits can suppress previously initiated tumors. It is unclear whether LNPs reaching tumor tissue from the bloodstream can deliver enough mRNA to a sufficient number of tumor cells to achieve tumor suppression. As a first step, to assess delivery, we initiated tumors (day 0) using the HDT procedure above, along with GFP to label cells receiving the oncogenic constructs (**Fig. 4A**). On day 5, we administered a single dose of nuclear-localized tdTomato mRNA-LNP, and sacrificed animals 12 hours later for analysis. We observed efficient transfection of healthy hepatocytes, as indicated by tdTomato fluorescence (**Fig. 4A**, **fig. S19**). We also observed GFP-positive tumor clones of ∼5 to ∼100 cells scattered throughout the liver. Within these transformed clones, many cells exhibited strong co-localized tdTomato and GFP fluorescence, indicating that the LNP delivery system can access transformed cells (**Fig. 4A**, **fig. S19**).

**Fig. 4.**
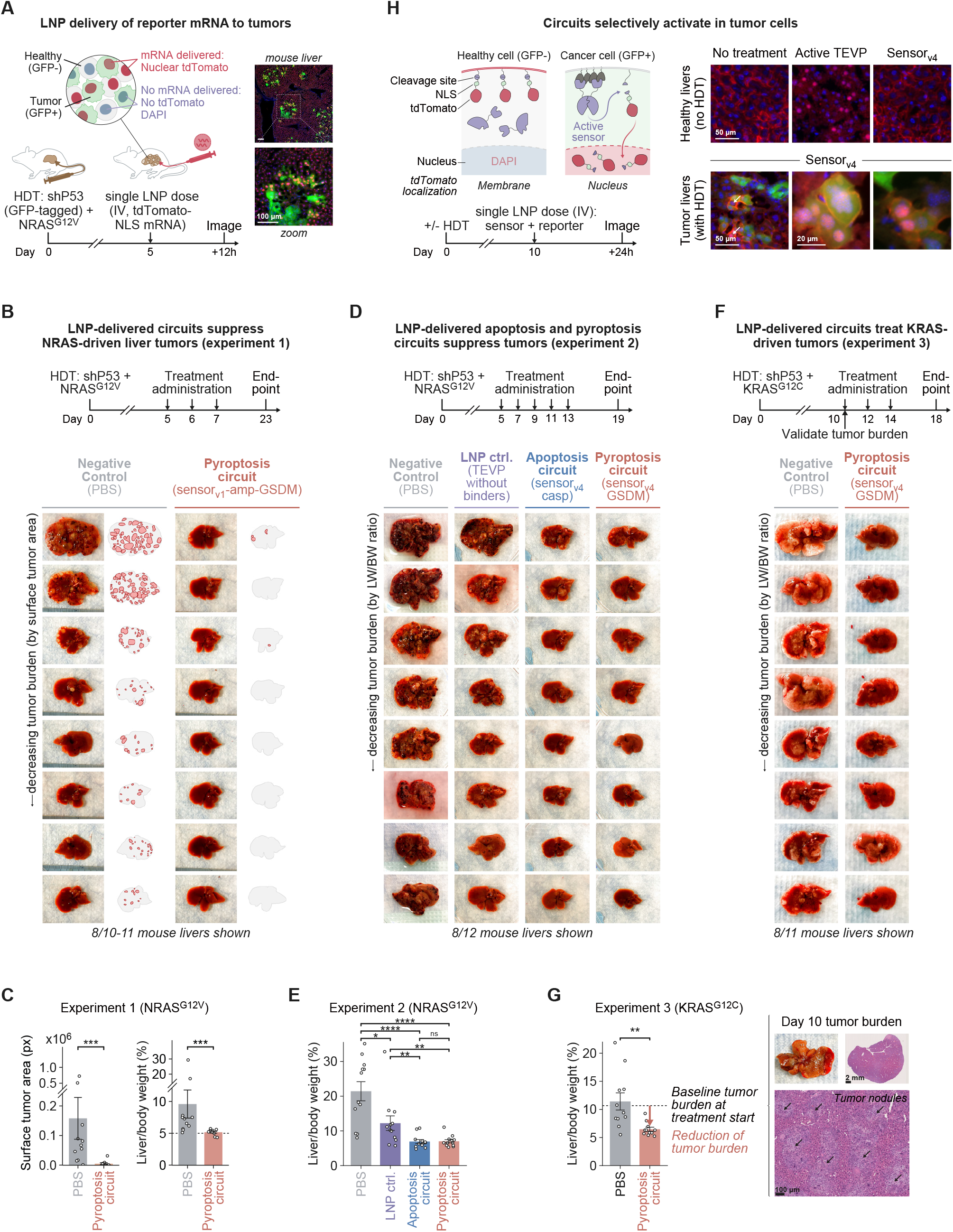
Systemically delivered circuits suppress RAS-driven liver tumors. **(A)** Single LNP administration enables delivery of nuclear-localized tdTomato mRNAs (red) to liver tumors (HDT with GFP-tagged shP53 + NRAS^G12V^, green). Images show a representative mouse liver stained with DAPI (blue). **(B)** Systemically delivered sensor_v1_-amp-gsdm circuit suppresses NRAS-driven liver tumors. Representative livers and blinded surface tumor annotations (red) are shown. **(C)** Quantification of tumor burden in the NRAS^G12V^ model from **(B). (D)** Apoptosis and pyroptosis circuits enable in vivo tumor suppression without an amplifier. **(E)** Liver-to-body weight ratios for the NRAS^G12V^ model from **(D). (F)** Circuits treat established KRAS-driven liver tumors. **(G)** Left: Liver-to-body weight ratios for the KRAS^G12C^ model from **(F).** Dashed line indicates baseline tumor burden at treatment start (day 10). Right: representative day-10 liver and H&E staining showing established tumor nodules. **(H)** Circuits selectively activate in tumor cells in vivo. Sensor activity was assessed using a membrane-tethered tdTomato reporter that relocates to the nucleus upon TEV protease cleavage, co-delivered with circuit sensors as mRNA-LNP. **(C)**, **(E)**, and **(G)**: Bars depict mean ± s.e.; data points represent individual animals.

To test the efficacy of systemically-delivered therapeutic circuits, we initiated tumors, allowed them to develop for five days, administered the sensor_v1_-amp-GSDM circuit as mRNA-LNP (1.5 mg/kg) three times, on days 5, 6, and 7, and sacrificed mice once control animals showed signs of distress (here, day 23) to examine livers and measure tumor burden (**Fig. 4B**). Strikingly, animals treated with the circuit showed near-complete tumor clearance compared to the PBS-treated control group (**Fig. 4B**, **fig. S20**). Circuit-treated animals were almost entirely free of surface tumor nodules (p = 0.00055) and exhibited liver-to-body weight ratios comparable to those of healthy animals (p = 0.00079; **Fig. 4C**, **fig. S21**, **table S2**).

Encouraged by these results, we performed a second round of *in vivo* efficacy experiments to determine whether the optimized and simplified versions of the circuits could be effective. We used sensor_v4_ along with apoptosis or pyroptosis effectors, omitting the amplifier entirely. We induced tumors as above, systemically delivered repeated doses of LNP circuits on days 5, 7, 9, 11, and 13 and analyzed livers on day 19 (**Fig. 4D**). Both the apoptosis (p = 6.7 x 10^-5^ versus PBS control) and pyroptosis (p = 5.3 x 10^-5^ versus PBS control) circuits strongly suppressed tumor development (**Fig. 4, D and E**, **fig. S22 and S23**, **table S3**). Negative control LNPs containing only the split TEV protease halves without RAS-binding domains or a functional effector failed to achieve comparable reductions in tumor burden, indicating that tumor suppression could not be explained by the LNP alone (LNP ctrl. vs. apoptosis circuit: p = 0.010; LNP ctrl. vs. pyroptosis circuit: p = 0.010; **Fig. 4, D and E**, **fig. S22 and S23**, **table S3**). Qualitatively, the apoptosis and pyroptosis effectors exhibited similar degrees of tumor suppression. This may reflect saturated therapeutic efficacy under these experimental conditions, or overlapping downstream responses from the two effectors, possibly through caspase activation of endogenous GSDME in hepatocytes ^66,81^. The circuit-treated, dissected liver tumors exhibited increased levels of cleaved PARP, a caspase-3 substrate, and T cell marker CD3, suggesting that both circuits activated caspase and stimulated immune recruitment (**fig. S24**). We did not observe substantial changes in phospho-ERK levels compared to PBS-treated controls, suggesting circuits function through direct induction of cell death rather than ERK suppression (**fig. S24**; *see further discussion below*).

We next sought to address two critical questions: whether circuits can be effective against more established tumors, and whether circuits could respond to G12C mutations in the more clinically relevant KRAS (rather than NRAS) isoform *in vivo*. We induced tumors using KRAS^G12C^, but otherwise retained the same HDT tumor initiation protocol. We also extended the delay before treatment to 10 days when signs of more advanced disease were clearly evident, including elevated liver-to-body weight ratios, grossly visible lesions covering the liver surface, and histologically confirmed aggressive, multifocal tumor nodules (**Fig. 4G**, **fig. S25**), and we confirmed that LNPs can access these nodules (**fig. S26**). On days 10, 12, and 14, we treated animals with the sensor_v4_-GSDM mRNA-LNP circuits, and harvested livers on day 18 (**Fig. 4F**). While PBS-treated negative control animals progressed compared to the day-10 baseline, circuit-treated animals showed near-normal liver-to-body weight ratios (p = 0.0014), grossly normal livers, and a reduction in the number of visible nodules (**Fig. 4, F and G**, **fig. S27 and S28**, **table S4**). Notably, the liver-to-body weight decline below the day-10 baseline in circuit-treated animals suggests active regression of established disease, rather than tumor suppression (**Fig. 4G**, **fig. S25**). Further *in vivo* studies that initiate LNP circuit treatment in mice with different tumor stages will be needed to more precisely define the extent to which circuit efficacy depends on disease stage.

Finally, we performed pharmacodynamic analysis to verify tumor-specific sensor activation *in vivo*. We engineered a membrane-tethered tdTomato reporter that relocates to the nucleus in response to TEV protease activity. At day 10 post tumor initiation, we co-delivered mRNA-LNPs encoding sensor_v4_ and the tdTomato reporter to KRAS^G12C^ tumor-bearing mice, and collected livers 24 hours later (**Fig. 4H**, **Methods**). As a control, we also delivered these LNPs to healthy mice that did not undergo HDT. In healthy livers, no detectable nuclear translocation was observed, even at high sensor mRNA doses (1 mg/kg), while a positive control construct encoding a constitutively active TEV protease exhibited clear and homogeneous nuclear translocation already at low mRNA doses (0.1 mg/kg; **Fig. 4H**). By contrast, in tumor-bearing livers, translocation did occur (**Fig. 4H**). Critically, translocation was exclusively observed in GFP-positive cells, confirming that circuit activation is restricted to tumor cells. These results confirm that the sensor_v4_ exhibits a low background, high specificity, and sufficient sensitivity *in vivo*.

Taken together, these experiments demonstrate that LNPs can deliver mRNA-encoded therapeutic circuits to tumor and normal cells, that circuits can selectively activate in tumor cells, and that they can treat NRAS-and KRAS-driven liver tumors.

### Circuits provide key advantages over inhibitors

Therapeutic circuits and inhibitors appear to represent distinct ways to achieve largely the same goal—tumor cell removal. However, they work through distinct mechanisms of action. To understand the implications of these mechanistic differences, we directly compared the LNP-delivered circuit (sensor_v4_-casp) with two RAS inhibitors: Sotorasib, the clinically approved covalent KRAS^G12C^-targeting drug ^82–84^, and RMC-7977, a tool compound of the breakthrough pan-RAS inhibitor Daraxonrasib ^85,86^.

We first compared the specificity and efficacy of the three treatments. We treated RAS wild-type and mutant cells with saturating concentrations of Sotorasib, RMC-7977, or the sensor_v4_-casp circuit for 5 days. As expected, the inhibitors reduced viability in RAS-addicted, RAS-mutant cell lines (Calu-1, MIA PaCa-2, NCI-H358), but did not perturb RAS-wild-type cells (HEK293; **Fig. 5A**). The sensor_v4_-casp circuit reduced RAS-mutant cell viability as well, and did so more strongly, to near background levels. This occurred without detectable inhibition of the off-target HEK293 cells (**Fig. 5A**). Thus, the circuit achieved comparable specificity and greater efficacy compared to the drugs against RAS-addicted cells.

**Fig. 5.**
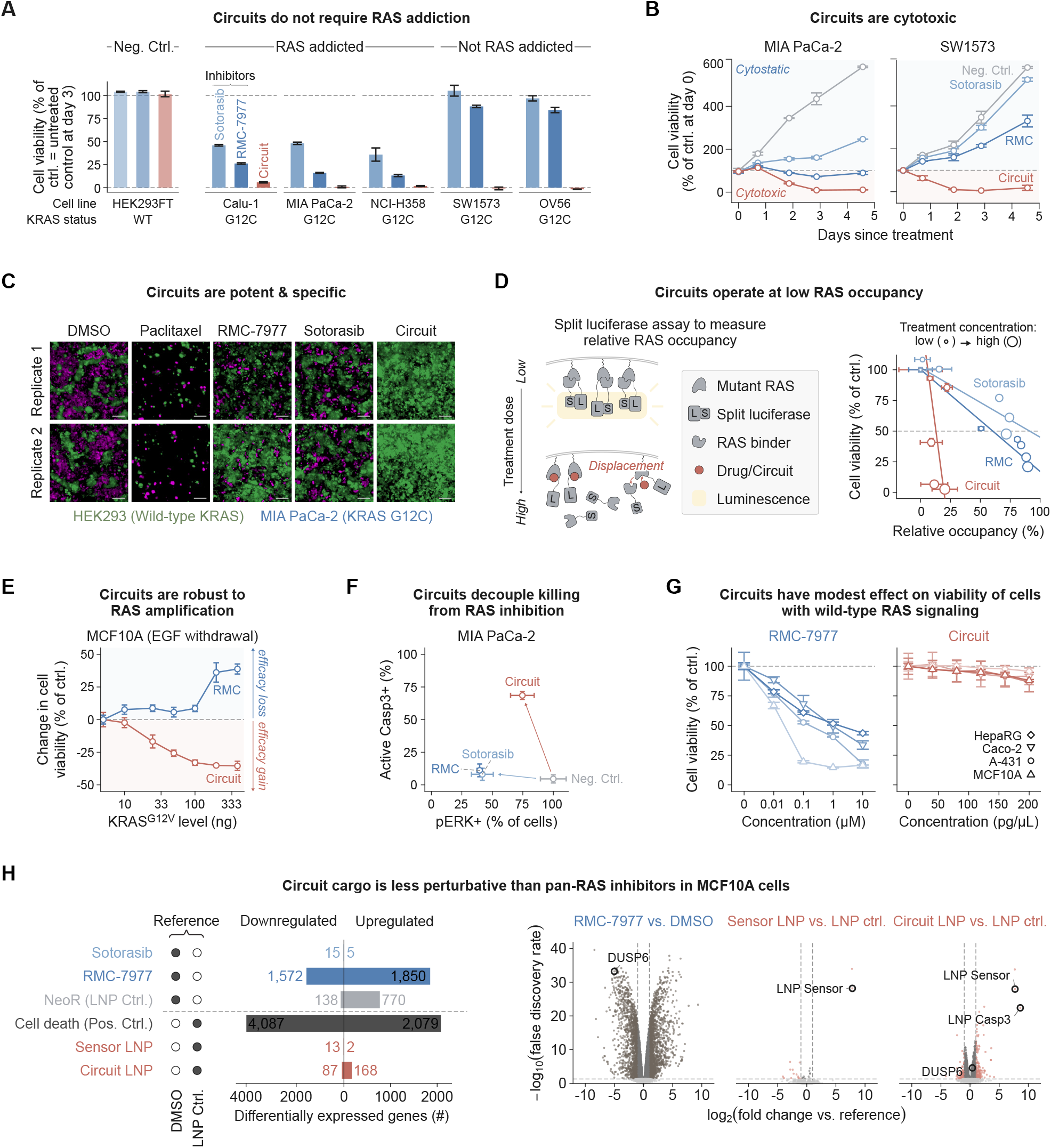
Circuits provide key advantages over RAS inhibitors. **(A)** Cell viability of RAS-addicted and non-addicted cancer cell lines treated with saturating concentrations of Sotorasib, RMC-7977, or sensor_v4_-casp circuit. **(B)** Cell viability over time in MIA PaCa-2 (KRAS^G12C^) and SW1573 (KRAS^G12C^) cells treated with negative control, Sotorasib, RMC-7977, or sensor_v4_-casp circuit. Viability expressed as a percentage of day-0 control. **(C)** Fluorescence images of co-cultured HEK293 (KRAS^WT^, green) and MIA PaCa-2 (KRAS^G12C^, magenta) cells treated with DMSO, Paclitaxel (30 µM), RMC-7977 (1 µM), Sotorasib (1 µM), or sensor_v4_-casp circuit (150 pg/µL engineered caspase-3 + 100 pg/µL sensor_v4_). Images acquired on day 5. Scale bar, 200 µm. **(D)** Left: schematic of split-luciferase RAS-RBD displacement assay measuring relative RAS occupancy. Right: Mean cell viability as a function of relative RAS occupancy in MIA PaCa-2 cells treated with Sotorasib, RMC-7977, or sensor_v4_-casp circuit at increasing concentrations (filled circles, low; open circles, high). **(E)** Change in MCF10A cell viability after EGF withdrawal as a function of ectopic KRAS^G12V^ expression level; dashed line separates efficacy loss (above) from efficacy gain (below). **(F)** Cleaved caspase-3 and pERK staining of MIA PaCa-2 cells treated with sensor_v4_-casp circuit, RMC-7977, Sotorasib, or negative control. **(G)** Cell viability of RAS wild-type cell lines representing tissues implicated in pan-RAS inhibitor toxicity. **(H)** RNA-seq analysis of MCF10A cells comparing transcriptomic perturbations across treatments. Left: number of differentially expressed genes for each condition relative to DMSO (Sotorasib, RMC-7977, NeoR LNP control) or LNP control (cell-death positive control, sensor LNP, circuit LNP). Right: volcano plots showing significantly down- and up-regulated genes. Error bars in **(A)**, **(B)**, **(D)**, **(E)**, and **(G)** represent mean ± s.d.

A key advantage of the circuit is that, unlike targeted inhibitors, it does not depend on oncogene addiction. In the case of RAS, oncogene addiction varies widely, as can be observed in the DepMap CRISPR essentiality database (**fig. S29**) ^87^. Consistent with their reliance on oncogene addiction, the two inhibitors minimally reduced viability of the non-addicted cell lines SW1573 and OV56 (**Fig. 5A**), despite potently suppressing pERK (**fig. S30**). By contrast, the circuit eliminated these non-addicted cell lines as effectively as the addicted ones (**Fig. 5A**), consistent with its sense-kill mechanism, which does not depend on cellular responses to pathway inhibition.

Circuits could also differ from inhibitors in terms of their dynamics. To compare the cell killing dynamics of circuits to those of inhibitors, we tracked cell viability over 5 days of treatment. In RAS-addicted MIA PaCa-2 cells, Sotorasib reduced proliferation, but maintained viable cell numbers near their pre-treatment baseline, while RMC-7977 induced a mixture of growth inhibition and partial cytotoxicity (**Fig. 5B**). By contrast, the circuit rapidly reduced cell numbers below Day 0 baseline, and achieved near-zero viability within three days through apoptotic killing, as confirmed by Annexin staining (**Fig. 5B**, **fig. S31**). These differences could be observed vividly in time-lapse movies of co-cultured RAS wild-type (HEK293) and RAS-mutant (MIA PaCa-2) cells, labeled with distinct fluorescent proteins. While saturating doses of Sotorasib and RMC-7977 reduced the growth of mutant cells, the circuit actively eliminated them within the first few days of the experiment (**Fig. 5C**, **fig. S32**, **movie S1**).

Together, these results highlight a fundamental difference between drugs and circuits. The response of a cell to an inhibitor depends on intrinsic cellular pathways (**Fig. 6F**). Many aspects of a cancer cell’s state, including but not limited to oncogene addiction, indirectly impact its response to drugs, which can range from slowed proliferation to eventual cell death. In contrast, the therapeutic outcome of a circuit—direct cell death—is encoded and executed directly by the circuit’s own effector components (**Fig. 6F**), which can act rapidly and potently.

**Fig. 6.**
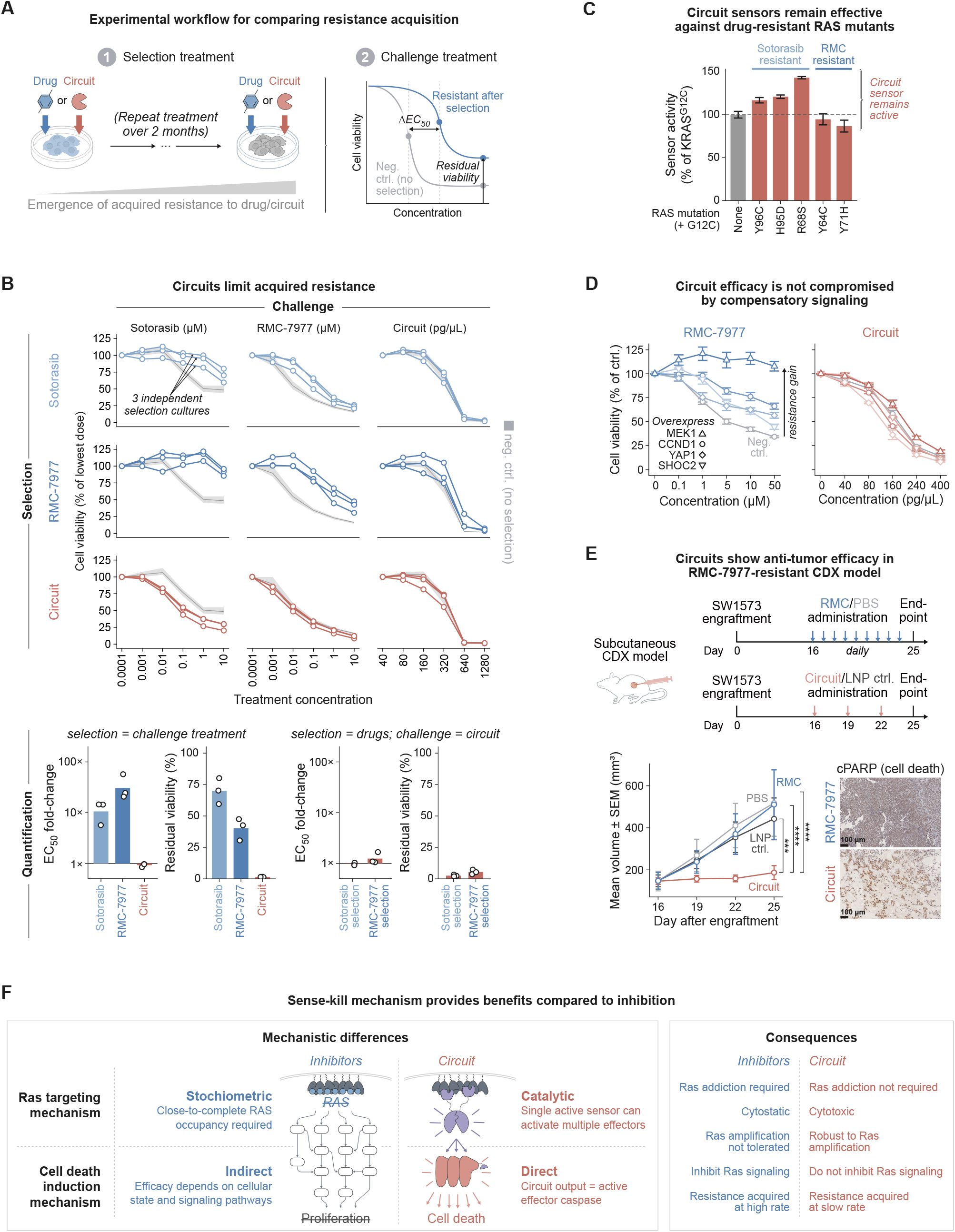
Circuits limit acquired resistance and remain potent against drug-resistance mechanisms. **(A)** Experimental workflow for comparing resistance acquisition. **(B)** Top: Dose-response curves for all selection-by-challenge combinations. Rows denote the selection, columns the challenge treatment. Each curve represents one of three independent selection cultures; shaded reference denotes the non-selected negative control. Bottom: quantification of EC_50_ fold-change (relative to non-selected control) and residual viability (%) for selection = challenge conditions (left) and for drug-selected cultures challenged with circuit (right). Individual data points represent independent selection cultures (based on n = 3 measurements). **(C)** Sensor_v4_ remains active against clinically relevant drug-resistant KRAS variants. **(D)** Circuit efficacy is not compromised by compensatory bypass signaling. Negative control represents cells without overexpression. **(E)** Top/left: Anti-tumor efficacy of circuits in an RMC-7977-resistant subcutaneous CDX model (SW1573 cells). Right: cleaved PARP (cPARP) immunohistochemistry in RMC-7977- and circuit-treated tumors. **(F)** Summary schematic and table comparing sense-kill circuits to inhibitors. The downstream network is a conceptual illustration and not intended to represent specific proteins.

### The catalytic circuit mechanism allows sensitive targeting of RAS-mutant cells

For targeted inhibitors, each drug molecule binds to and inhibits one target molecule. Because unoccupied target molecules can continue to signal ^88,89^, inhibition requires near-saturating occupancy. For example, clinical response to the BRAF inhibitor PLX4032 in melanoma patients required greater than 80% ERK phosphorylation inhibition ^88^. Circuits, on the other hand, function catalytically: each reconstituted sensor protease bound to a pair of mutant RAS proteins can activate multiple cell death effector proteins, which can then each cleave and activate multiple downstream substrates. This cascade and catalytic mechanism makes circuits potentially more sensitive to mutant RAS.

To test this hypothesis, we used a previously described split luciferase RAS-RBD displacement assay ^90,91^ to compare the activity of inhibitors to that of the sensor_v4_-casp circuit at equivalent levels of RAS occupancy (**Methods**, **Fig. 5D**, **fig. S33**). As expected, Sotorasib and RMC-7977 required high RAS occupancy to reduce the viability of MIA PaCa-2 cells (KRAS^G12C^). By contrast, the circuit eliminated nearly all cells at much lower occupancies (**Fig. 5D**), consistent with a catalytic mechanism in which sparse RAS engagement is amplified into a potent response.

The difference between stoichiometric and catalytic modes of action has implications for RAS amplifications, a common mechanism of resistance to RAS-inhibitors ^92,93^. For a catalytic sensor, but not a stoichiometric inhibitor, elevated RAS expression should enhance circuit-based killing by increasing productive sensor-reconstitution events. To test this, we titrated ectopic KRAS^G12V^ expression and measured RAS-dependent proliferation of MCF10A cells after EGF-withdrawal ^94^ in the presence of either RMC-7977 or the sensor_v4_-casp circuit (Methods). As anticipated, RMC-7977 efficacy declined with increasing KRAS^G12V^ expression, recapitulating amplification-driven resistance, while the sensor_v4_-casp circuit showed the opposite behavior, killing cells more efficiently at elevated KRAS^G12V^ expression (**Fig. 5E**, **fig. S34**). These results show how catalytic therapeutic circuits can efficiently target cancer cells with drug-resistant RAS amplifications.

The catalytic activity of circuits also led to non-intuitive effects on RAS signaling. Both Sotorasib and RMC-7977 strongly reduced pERK levels in MIA PaCa-2 (KRAS^G12C^) cells, consistent with their designed mechanisms, but only modestly induced cell death (**Fig. 5F**, **fig. S35**). By contrast, the sensor_v4_-casp circuit strongly activated caspase-3 and cell death, with only a modest reduction in pERK (**Fig. 5F**, **fig. S35**). Thus, the circuit kills cells with minimal inhibition, while the inhibitors inhibit signaling, with minimal cell killing.

Pan-RAS inhibitors cause on-target, off-tumor toxicity in RAS-dependent tissues ^95,96^. The mutant-specific 12VC1 monobody could reduce such off-tumor toxicity by restricting circuit activity to cancer cells. To test this, we treated RAS wild-type cell lines representing tissues implicated in RAS inhibitor toxicity with either RMC-7977 or the sensor_v4_-casp circuit. These lines included intestinal epithelial (Caco-2), keratinocyte (A-431), mammary epithelial (MCF10A), and hepatocyte-like (HepaRG) cells. RMC-7977 caused dose-dependent reductions in viability across the panel, whereas the circuit generated more modest effects (cell viability *≥* 85% at 200 pg/uL, **Fig. 5G**). Consistent with this selectivity, the circuit also did not reduce viability in cells where wild-type RAS was activated by EGF stimulation, even at concentrations that potently stimulated pERK activity (from ∼15% to ∼70%; **fig. S36**). This ability to discriminate between wild-type and mutant Ras is consistent with the exquisite structural specificity of the 12VC1 monobody.

Finally, to identify potential off-target perturbations, we performed a transcriptome-wide comparison of Sotorasib, RMC-7977, and the sensor_v4_-casp circuit in RAS-wild-type MCF10A cells (**Fig. 5H**, **fig. S37**, **Methods**). Sotorasib exhibited few detectable effects, due to its mutant-specificity. RMC-7977 strongly downregulated the canonical RAS target gene DUSP6 and induced reciprocal KRAS upregulation ^97,98^ at 24 hours, consistent with on-target pathway suppression and transcriptional rebound. This was accompanied by perturbations of MAPK/RAS signaling, cell cycle pathway signatures (**fig. S37, A and B**), as well as global transcriptome perturbations (1,572 downregulated, 1,850 upregulated; **Fig. 5H**, **fig. S37C**). To control for non-specific effects of LNP delivery, we analyzed negative control LNPs lacking the circuit. These induced genes associated with innate immune responses, including upregulation of interferon and mRNA-sensing pathways ^99–101^ (138 downregulated, 770 upregulated; **Fig. 5H**, **fig. S37**). We also confirmed that a constitutively active cell death effector (active TEVP + TEVP-activated caspase-3) could induce widespread transcriptome changes (4,087 downregulated, 2,079 upregulated; **Fig. 5H**, **fig. S37**). Critically, however, LNPs incorporating circuit cargo generated few additional effects beyond those observed in negative control LNPs, even though the delivered circuit mRNA was highly expressed (87 downregulated, 168 upregulated; **Fig. 5H**, **fig. S37**). This could reflect the limited impact of the circuit on signaling.

Taken together, these studies show that inhibitors and circuits use fundamentally distinct mechanisms that have non-intuitive yet critical therapeutic implications. Rather than passively relying on oncogene addiction, circuits sense RAS and use cell death effector proteins to actively and rapidly induce cell death, leading to increased efficacy and cytotoxicity with or without RAS addiction. Because circuits function catalytically, they are more sensitive to RAS amplifications, but have less effect on cell signaling and are less perturbative at the transcriptome level (**Fig. 6F**).

### Circuits limit acquired resistance

Acquired resistance is perhaps the most important limitation of targeted therapies ^102,103^. By actively executing cells, therapeutic circuits could limit the time available to acquire resistance. Additionally, because the circuits are less reliant on endogenous components for their effects, there may be fewer potential pathways to resistance. These considerations suggest the hypothesis that circuits could be less susceptible to resistance compared to inhibitors.

To test this hypothesis, we subjected MIA PaCa-2 cells to 2 months of continuous selection on either Sotorasib, RMC-7977, or the sensor_v4_-casp circuit, each at its own EC_70_. To quantify resistance and cross-resistance, we then challenged each culture with all treatments. Each selection was performed in three independent culture replicates, accompanied by non-selected negative controls (**Methods**, **Fig. 6A**).

Inhibitors rapidly lost potency and efficacy after selection, as quantified by increased EC_50_ and residual viability at saturating treatment doses (**Fig. 6B**, **fig. S38 and S39**). More specifically, Sotorasib selection increased EC_50_ by 5- to 15-fold across replicates, and increased the fraction of cells surviving at maximum drug concentration (residual viability) from 48% (no selection) to 60-80% (after selection). RMC-7977 also increased EC_50_ by 21- to 57-fold across replicates, while increasing the fraction of cells surviving at maximum drug concentration from 16% (no selection) to 31-43% (after selection). Notably, resistance to one drug conferred cross-resistance to the other. RMC-7977-selected cultures were near completely resistant to Sotorasib (survival at 10 µM: 91% versus 48% without selection), and Sotorasib-selected cultures were cross-resistant to RMC-7977 (up to 12-fold EC_50_ shift; **Fig. 6B**, **fig. S38 and S39**). This cross-resistance suggests that cells adapted to RAS inhibition, rather than to the drug *per se*.

Strikingly, circuit-selected cells showed no significant increase in EC_50_ or residual viability when challenged with circuit, and no loss of sensitivity to either drug (**Fig. 6B**, **fig. S38 and S39**). Sotorasib- and RMC-7977-resistant cells challenged with the circuit also showed minimal cross-resistance (**Fig. 6B**, **fig. S38 and S39**). The finding that circuits showed minimal to no resistance acquisition under continuous selection was consistent across independent experiments (**fig. S40 and S41**). Together, these results strongly suggest that, at least under these conditions, therapeutic circuits are less susceptible to resistance acquisition than inhibitors.

### Circuits circumvent drug-resistance mechanisms

An unsolved therapeutic challenge is targeting tumors that have evolved resistance to RAS inhibitors. Prevalent acquired resistance mechanisms include *cis* mutations in KRAS that prevent inhibitor binding ^104–106^. To determine whether the circuit could sense such mutants, we evaluated the response of the 12VC1 sensor to a panel of clinically relevant KRAS^G12C^ variants that confer resistance to Sotorasib (H95D, Y96C, R68S) and RMC-7977 (Y71H, Y64C) ^106^. The sensor maintained robust activity against all tested resistant variants, comparable to its response to parental KRAS^G12C^ (**Fig. 6C**).

Drug resistance can also occur through activation of parallel signaling proteins such as MEK1, CCND1, YAP1, and SHOC2 that restore proliferative signaling in the presence of RAS inhibition ^107–111^. As expected, overexpression of each of these proteins drastically reduced RMC-7977 efficacy (**Fig. 6D**). However, the same perturbations did not systematically reduce the sensor_v4_-casp circuit efficacy (**Fig. 6D**), consistent with the circuit’s sense-kill mechanism.

Finally, we asked whether these resistance advantages extend to an *in vivo* setting. We found that RMC-7977 efficiently cleared tumors in the NRAS^G12V^/shTP53 HDT model described above over a 28-day experiment (p = 0.022, Methods, **fig. S42 and S43**, **table S5**). Thus, this model did not permit analysis of potential differences in resistance.

We therefore turned to SW1573 xenografts, an established RAS-inhibitor-resistant model in which a co-occurring PIK3CA mutation is associated with RAS-independent pro-survival signaling ^112–115^. Animals received either a negative control (PBS), daily oral RMC-7977 (10 mg/kg), LNP negative control, or the LNP-delivered sensor_v4_-GSDM circuit. LNP conditions and PBS were administered by intratumoral injection every three days (**Fig. 6E**). While RMC-7977 had no detectable effect on tumor progression (p = 0.473 versus PBS), circuits actively triggered tumor cell death (IHC staining, caspase-3 substrate PARP), and strongly suppressed tumor growth (p = 6.72 x 10^-4^ versus RMC-7977; 9.69 x 10^-5^ versus PBS; **Fig. 6E**, **fig. S44, S45, and S46**, **table S6**).

Taken together, these results suggest that therapeutic circuits can target drug-resistant cells in culture and *in vivo*. This finding is particularly relevant as RAS inhibitors are likely to become standard-of-care for many cancers, and select for inhibitor-resistant tumors.

## Discussion

The astonishing effectiveness of natural tumor suppression circuits like p53 provokes the fundamental questions of whether it is possible to design analogous synthetic sense-and-respond circuits, how effectively such circuits could function in cells and organisms, and what advantages they might provide compared to existing treatment modalities. The answers to these questions emerge by considering intrinsic properties of the circuits themselves, interactions of the circuits with endogenous cellular components, and the broader physiological context in which circuit therapies operate.

Therapeutic circuits are based on two core circuit-intrinsic design principles: catalytic sensing and mutant-specificity. Catalytic sensing is made possible by split-protease reconstitution, which provides a sensitive mechanism for detecting membrane-bound, mutant RAS that should be applicable to other membrane-bound cancer-specific target proteins. The use of mutant-specific binders such as 12VC1 enables remarkable discrimination of cancer and normal cells (**Fig. 2A**). Rapid advances in binder design should allow design of mutant-specific binders to other cancer-specific target proteins. Through catalytic sensing, the circuits are able to respond at low RAS occupancies where inhibitors are ineffective (**Fig. 5D**) and—counterintuitively—are more, rather than less, effective against cells with RAS amplification (**Fig. 5E**), a common driver of resistance to RAS inhibitors.

The effectiveness of a therapy depends on the cellular responses it evokes. For inhibitors, the fate of a target cancer cell depends on how the cell’s signaling network responds to the loss of oncogenic input. When cells exhibit oncogene addiction, the eventual outcome can include proliferative arrest or eventual cell death. In contrast, the efficacy of therapeutic circuits does not depend on suppression of an endogenous pathway. By directly coupling sensing to engineered cell death effectors (**Fig. 6F**), therapeutic circuits function cytotoxically, actively eliminating cells instead of inhibiting growth (**Fig. 5B**). Thus, circuits are able to function independently of oncogene addiction (**Fig. 5A and 6E**), directly eliminating non-addicted RAS-mutant cells that are refractory to inhibitors in cells and *in vivo*.

Therapeutic circuits are capable of specifically eliminating cancer cells with reduced susceptibility to resistance compared to inhibitors. Prolonged selection with the circuit over two months did not drive acquired resistance, in sharp contrast to the large efficacy and potency shifts observed under equivalent inhibitor selection (**Fig. 6B**). This capability stems from two core features of the therapeutic circuit. First, it immediately kills mutant cells, limiting the time available for the cell to acquire resistance. Highly transfected cells were eliminated within hours, and most mutant RAS cells were eliminated within a few days (**movie S1**). Second, therapeutic circuits’ low reliance on endogenous pathways for cell killing provides fewer molecular pathways to resistance. For example, the circuit circumvents prevalent bypass signaling pathways such as MEK1 and YAP1 that confer RAS inhibitor resistance (**Fig. 6D**). In an inhibitor strategy, even a single copy of RAS with a *cis*-mutation that prevents inhibitor binding could provide sufficient signaling to confer drug-resistance. However, the situation with the circuit is reversed: the circuit can still respond as long as some copies of RAS can be bound by the sensor, making it more difficult to evade the circuit with a binding-resistant mutation in a single copy of RAS. While our data suggests that circuits can reduce susceptibility to resistance, we cannot rule out the possibility of resistance through mechanisms such as reduced LNP uptake or dominant suppressors of cell death effectors. We anticipate that resistance to circuits, if it occurs, will likely arise at the interfaces between engineered circuit proteins and endogenous components, potentially making it easier to identify and proactively circumvent.

Systemically delivered therapeutic circuits suppressed aggressive, multifocal RAS-driven liver tumors in immunocompetent mice and drove regression of established disease (**Fig. 4**). The ability to couple sensing to either apoptotic or pyroptotic cell death provides complementary therapeutic strategies: apoptosis enables cell-autonomous killing, while pyroptosis releases inflammatory signals that can recruit immune responses and extend tumor clearance beyond directly transfected cells. More *in vivo* studies will be required to comprehensively evaluate circuit function in cell- and patient-derived xenografts as well as in metastatic settings, in order to better understand their therapeutic potential across diverse RAS-driven cancers. In particular, in-depth characterization of the immune response elicited by pyroptotic circuits *in vivo*, and experiments testing how apoptosis and pyroptosis should be balanced, will be important to maximize tumor clearance while minimizing potential toxicity from hyper-inflammation.

While intravenous LNP delivery to the liver has clinical precedent, access to most other tissues has only been demonstrated in preclinical studies ^69,116^. LNP delivery is a rapidly evolving field, advanced by technologies such as selective organ targeting (SORT) ^57^, and thus the full scope of what systemic LNP delivery can and cannot achieve in humans has yet to be defined. Should certain tissues remain inaccessible through systemic LNP delivery, intratumoral administration provides a viable alternative in the context of cancer. Additionally, circuits could use other modalities such as oncolytic viruses, extracellular vesicles, or, in the future, cells themselves as mRNA delivery vehicles ^117–120^. Regardless, the circuit paradigm effectively shifts the burden of specificity from the delivery method to the circuit, making circuits compatible with higher penetrance but less tissue-specific delivery methods. Finally, a potential concern with any circuit that uses non-human proteins is immunogenicity. Here, we used a viral protease as a major component. However, more recent work shows how similar effects can be achieved without non-human protein domains.

Therapeutic circuits are a programmable technology. Replacing the sensor in the current circuits could retarget them to other cancer drivers such as MYC, *β*-catenin, or other oncogenes. Some potential targets may be more difficult to target through point-mutation-based selectivity and could require sensing of aberrant expression levels or cancer-specific post-translational modifications. In fact, circuits do not need to respond to oncogenes at all, but could be designed to sense proteins that provide accurate markers of disease state. Looking ahead, the growing power of computational protein design methods should enable rapid generation of effective binders for cancer-associated markers, broadening the therapeutic circuit repertoire. Circuit outputs could similarly be broadened to include cytokine release or other therapeutic responses ^15,121^.

Together, these considerations show that therapeutic circuits could provide higher efficacy and less opportunity for resistance than targeted therapies, greater molecular specificity than chemotherapies, and an expanded ability to sense intracellular proteins compared to CAR-T cells and antibody-drug conjugates. Looking ahead, circuits could provide a potent, specific, and programmable mechanism for treating cancer and other human diseases.

## Acknowledgements

We thank Duncan Chadly, Dongyang Li, Rongrong Du, Martin Tran, Jacob Parres-Gold, Jan Gregrowicz, Evan Mun, Ron Hadas, Ethan Richman, Jordan Lay, Sean Hsieh, Nikita Makarov, Roey Lazarovits, and Arun Chakravorty for advice on experimental design, technical support, insightful discussions, or critical proofreading of the manuscript; Inna-Marie Strazhnik for graphical design; Leah Santat, Jo Leonardo, and Rui Malinowski for administrative support.

## Author contributions

A.C.L., L.M., and M.B.E. conceived and designed the study. S.M., A.C.L., and L.M. conceived and designed *in vivo* experiments. M.B.E. directed and supervised the study. D.J.S., H.Z., and M.B.E. directed and supervised *in vivo* experiments. A.C.L., L.M., K.H., and A.Q. performed *in vitro* and computational experiments. V.T., E.Z., S.X., M.W.B., H.L., A.A.D., J.M.L., M.J.F., Q.I., and X.J.G. assisted with or offered guidance regarding *in vitro* or computational experiments. S.M., Z.L., and M.W. performed *in vivo* experiments. B.G. performed RNAseq experiments. L.K. manually annotated liver surface tumors. A.C.L., L.M., and S.M. analyzed data. A.C.L., L.M., and M.B.E. wrote the manuscript and designed figures with input from all authors.

## Data and materials availability

At the date of publication, data will be deposited at data.caltech.edu and become publicly available. Accession numbers will be listed in the key resources table. Raw RNA sequencing data will be deposited at the NCBI Sequence Read Archive. Raw microscopy images will also be deposited online. Any additional information required to reanalyze the data reported in this paper is available from the lead contact upon request. All material requests and correspondence should be addressed to M.B.E.

