## Supplemental Information for "Engineered protein circuits for cancer therapy"

#### Supplementary Materials

Materials and Methods

Tables S1-S6

Figures S1-S46

Movie S1 (separate)

Key Resource Table

Supplementary References

### Materials and Methods

#### Plasmid construction

Plasmids were generated via Gibson Assembly (NEBuilder HiFi DNA Assembly Master Mix; New England BioLabs) or KLD cloning (T4 Polynucleotide Kinase, T4 DNA Ligase, DpnI, T4 DNA Ligase Buffer; Thermo Scientific). Gene fragments were sourced from Twist Bioscience or Integrated DNA Technologies (IDT), or PCR-amplified from existing laboratory plasmid constructs. IDT synthesized all PCR primers. Plasmids were purified (QIAprep Spin Miniprep Kit or Qiacube machine; Qiagen), normalized (Thermo Scientific NanoDrop 8000), and sequence-verified (Plasmidsaurus or Genewiz) prior to experimental use.

#### Tissue culture and cell lines

Most cell lines were obtained from ATCC and cultured under standard conditions (37°C, 5% CO2, humidified Eppendorf CellXpert C170i incubator). Adherent cells were maintained in Dulbecco's Modified Eagle Media (DMEM; Thermo Fisher) supplemented with 10% fetal bovine serum (FBS; Avantor), penicillin (1 unit/ml), streptomycin (1 µg/ml), glutamine (2 mM), sodium pyruvate (1 mM), and 1X Minimal Essential Media Non-Essential Amino Acids (all Thermo Fisher). HEK293FT cells were cultured for up to 20 passages, and human cancer cell lines for up to 15 passages, maintaining 10%-90% confluency. Cells were passaged using 0.25% Trypsin-EDTA (Thermo Fisher). Viable cell numbers for seeding densities were determined using Trypan Blue (Invitrogen) and the Countess 3 automated cell counter (Thermo Fisher). Cells were routinely confirmed mycoplasma-free (MycoStrip kit; InvivoGen). Cell lines with stably integrated fluorescent proteins used for microscopy were purchased from FenicsBIO.

#### Transient transfection

Transient transfections were performed utilizing FuGENE HD (Promega), Lipofectamine 3000 (Thermo Fisher), or LNPs (described below) as transfection agents. Unless otherwise noted, we seeded 10k cells (96-well plate), 200k cells (24-well plate) for FuGENE or LNP transfections, and 275k cells (24-well plate) for Lipofectamine transfections. Cells were reverse-transfected immediately after seeding following the manufacturer's instructions. For FuGENE transfections, we incubated 1 µg plasmid DNA with 6 µl transfection reagent. For FuGENE transfections, 200k cells were transfected with up to 566 ng of DNA. For Lipofectamine transfections, 1.5 µl Lipofectamine 3000 and 1.5 µl reagent P3000 were used with a total of 500 ng DNA to transfect an equivalent of 275k cells. The total amount of plasmid DNA applied in each transfection was scaled according to seeding densities.

#### Flow cytometry

Cells were collected two days post-transient transfection for flow cytometry, filtered through a 40-µm cell strainer, and analyzed by flow cytometry (CytoFLEX S flow cytometer, Beckman Coulter). Unless otherwise noted, we used FITC-A channel (GFP of iTEVP reporter[1]; excitation 488 nm, emission 525/40 nm; gain of 1), ECD-A channel (mRuby3; excitation 561 nm, emission 610/20 nm; gain of 1), PB450-A channel (mTagBFP2; excitation 405 nm, emission 450/45 nm; gain of 1), and APC-A700-A channel (IFP of iTEVP reporter; excitation 638 nm, emission 712/25; gain of 100). Live cells and singlets were gated using FlowJo (version 10.10, BD Biosciences). iTEVP reporter activation was assessed by calculating the median IFP fluorescence (activation) for a narrow GFP expression regime ($10^{5.25}$-$10^{5.5}$ a.u., unless otherwise noted; reporter normalization), corresponding to the reporter expression regime with maximal dynamic range. As the reporter construct was always transfected as a separate polytransfection mix, selecting cells with high reporter expression did not bias subsequent analysis steps, *i.e.*, the expression distribution of other constructs (*e.g.*, reporter or ectopic RAS) remained unaffected.

#### Design and Screening of Protease-Activatable Proteases

We designed a panel of candidate protease-activatable proteases based on diverse natural and synthetic protein activation mechanisms. All constructs were expressed in HEK293 cells via transient transfection for characterization. Amplifier expression levels were assessed by fluorescence of co-transfected Ruby (low amplifier expression: $10^3$-$10^4$ a.u., medium expression: $10^4$-$10^5$ a.u., high expression: $10^5$-$10^6$ a.u.). Confidence estimates were obtained by bootstrapping (95% confidence intervals; 1,000 bootstrap iterations).

Candidates 1-3: To emulate autoinhibition-based activation mechanisms observed in proteins such as WASP, SNARE, and ERM, we designed constructs in which a non-cleavable, synthetic autoinhibitory peptide was tethered to the active site of TVMV protease (TVMVP)[2]. The tether included a cleavage site for an orthogonal protease (TEVP), enabling activation through proteolytic removal of the inhibitory domain. Constructs were tested with and without co-expression of the activating TEVP protease. To modulate inhibitory dynamics, we generated a series of variants with progressively shortened linkers between the autoinhibitory peptide and TVMVP.

Candidates 4-7: To improve autoinhibition, we developed a tethering strategy in which the autoinhibitory peptide was anchored at both termini via two TEVP cleavage sites. We circularly permuted the TVMVP sequence and rejoined the new termini using antiparallel leucine zippers to stabilize the conformation. To optimize access of TEVP to its cleavage sites, we also varied the length of the linker flanking the autoinhibitory domain.

Candidates 8-10: We next explored strain-based inhibition, drawing inspiration from previous studies where circular permutation introduces conformational strain that disrupts protein function (*e.g.*, CRISPR)[3]. We engineered circularly permuted variants of TVMVP in which the old termini were joined by short linkers, embedding the TEVP cleavage site to allow strain release upon protease input. Additional constructs included deletions of disordered residues at the native termini to increase the distance between ends and enhance strain potential.

Candidates 11-13: To sterically inhibit the TVMVP active site, we fused bulky de novo designed heterodimers to the protease, positioned to occlude the catalytic cleft[4]. These heterodimeric blocking domains were flanked by TEVP cleavage sites to enable conditional displacement and activation.

Candidates 14-18: We adopted a protease caging strategy using split TVMVP[4–6]. One protease half was caged by fusion to a catalytically inactive TVMVP fragment via designed coiled coils. The coiled coil associated with the inactive half was flanked by TEVP cleavage sites, enabling conditional release. Additional constructs extended this strategy by caging both protease halves with inactive TVMVP fragments, each released via TEVP cleavage. To enable protease activation through strand displacement, we substituted the coiled coils with lower-affinity, sequence-matched variants previously reported to allow displacement by higher-affinity interactors. These designs were intended to facilitate TEVP-dependent exchange of inhibitory domains with active TVMVP halves.

#### Computational identification of RAS binding domains

To identify candidate RAS binding domains, we first extracted human proteins known to physically associate with KRAS from the STRING database[7] (physical subnetwork, medium confidence). All proteins passing these filters were co-folded with wild-type KRAS-4B and GTP using AlphaFold3 (AlphaFold Server)[8]. Binders predicted to interact with the CAAX domain of KRAS-4B were manually removed. To identify candidate binders for experimental characterization, we filtered for binders with high-confidence RAS interactions (interface PAE score < 10 and interface plDDT score > 75) and manually extracted RAS-interacting domains from the full-length proteins. For the proteins ARAF, RAF1, RGL1, and RASA4, we generated multiple candidate binder versions corresponding to different truncations or domains of the full-length protein. We also mutated sequence regions known to act as nuclear localization signals (NLS). Furthermore, we used RFdiffusion[9] (*Base_ckpt* and *Complex_beta_ckpt* models with default parameters; no hotspot residue selection) to *de novo* design RAS binders between 50 and 100 amino acids in length. Briefly, target structures (KRAS, HRAS, NRAS, including GTP or analog nucleotide as ligand) were obtained from the Protein Data Bank (PDB)[10]. PDB assemblies were filtered and processed to provide a single RAS domain as input for RFdiffusion. Sequence design was performed with ProteinMPNN-FastRelax (*dl_binder_design* pipeline)[11,12]. After co-folding candidate binders with their corresponding target using AlphaFold2[13] (*dl_binder_design* pipeline), binders were filtered for high-confidence RAS interactions (interface PAE score < 10 and interface plDDT score > 80). Binders with a high likelihood of dimerization in the absence of RAS were removed (co-folding of binder homodimers with AlphaFold3). In parallel, we compiled a list of synthetic RAS-binding domains such as monobodies[14], DARPins[15,16], synthetic proteins[17], and nanobodies[18] from the available literature. All sequences were codon-optimized for expression in human cells (Genscript).

##### Sensor screening

To evaluate the sensitivity and specificity of candidate RAS sensors, we reverse transfected HEK293FT cells (275k seeding density) with three separate plasmid mixes encoding (1) binder-cTEVP (120 ng), binder-nTEVP-IRES-mTagBFP2 (150 ng); (2) wild-type or mutant RAS (15 ng), mRuby3 (45 ng); and (3) iTEVP reporter fused to a CAAX membrane tether (170 ng). Transfections were performed with Lipofectamine 3000 according to the manufacturer's instructions. To characterize the binders, we gated for medium sensor (mTagBFP2 expression between $10^4$ and $10^6$ a.u.) and high RAS expression (mRuby3 expression between $10^5$ and $10^6$ a.u.). To analyze sensor responses to varying amounts of RAS, we gated for medium sensor expression (mTagBFP2 expression between $10^4$ and $10^6$ a.u.) and binned RAS expression as shown in plots. All plots show median reporter activation for the corresponding gate or bin. Confidence estimates for sensor screening experiments were obtained by bootstrapping (95% confidence intervals; 1,000 bootstrap iterations).

##### Sensor targeting scope characterization

To evaluate the targeting scope of RAS sensors, we tested the sensor performance against a list of RAS variants compiled by selecting all KRAS, HRAS, and NRAS variants from the TCGA pan-cancer database (accessed via the cbioportal server) that are present in at least 10 cancer patients[19,20]. Analogous to our sensor screening procedure, we reverse-transfected HEK293FT cells (200k seeding density) with three separate plasmid mixes encoding (1) binder-cTEVP (133 ng), binder-nTEVP-IRES-mTagBFP2 (180 ng); (2) wild-type or mutant RAS (13 ng), mRuby3 (40 ng); and (3) iTEVP reporter fused to a CAAX membrane tether (200 ng). Transfections were performed with FuGENE HD according to the manufacturer's instructions. To calculate the fold-change of the response to mutant vs. wild-type RAS, we gated for medium sensor (mTagBFP2 expression between $10^4$ and $10^{5.5}$ a.u.) and high RAS expression (mRuby3 expression between $10^5$ and $10^6$ a.u.) and divided the median activation of a given mutant RAS variant by the median activation of the corresponding wild-type RAS isoform. Confidence estimates were obtained by bootstrapping (95% confidence intervals; 1,000 bootstrap iterations). A similar scheme was used for testing sensors with 12D4 binders (mTagBFP2 expression between $10^3$ and $10^5$ a.u.), as well as mechanistic characterization of sensors (mTagBFP2 expression between $10^4$ and $10^5$ a.u.).

##### Mutant-specificity of the 12VC1-based sensor

In *Teng et al. 2021*, the 12VC1 monobody was originally characterized as binding $KRAS^{G12C}$ and $KRAS^{G12V}$ tightly but KRAS-G12D only marginally, too weak to inhibit RAS signaling. However, when we systematically profiled the 12VC1 sensor against a panel of KRAS, HRAS, and NRAS mutations in HEK293 cells, we found that it responded robustly to mutations beyond G12C and G12V, including G12D (**Fig. 1E**, **fig. S8**). Additionally, 12VC1 was previously characterized to inhibit RAS signaling and viability of mutant cells when stably expressed, while our 12VC1-based sensor alone did not detectably affect cell viability or pathway activity. We believe these two discrepancies reflect the same underlying principle. Circuit sensing, as we show in **Fig. 5D**, is catalytic and requires only a transient encounter with RAS, brief enough to reconstitute a protease. Thus, a binder that is too weak for RAS pathway inhibition could, in principle, remain sufficient for circuit sensing. These differences could also reflect differences in the sensor construct itself (*e.g.*, its inclusion of TEV domains that do not appear in *Teng et al. 2021*). We further confirmed that, at high expression, the 12VC1 monobody from *Teng et al.* 2021 does reduce cell viability. We delivered the monobody alone as mRNA-LNP and observed dose-dependent reductions in viability, reducing viability by ~70% at the highest monobody concentrations (fig. S9). This is consistent with *Teng et al.'s* findings: at sufficiently high expression, 12VC1 stoichiometrically occupies RAS, blocks RAF engagement, suppresses ERK signaling, and kills cells. The monobody, in this regime, functions as an inhibitor. In contrast, we confirm that even at the highest doses tested, the sensor produced only a modest reduction in cell viability (**fig. S9**).

##### Single-cell classification performance of sensors

To evaluate the ability of our sensors to discriminate between wild-type (WT) and mutant RAS proteins at the single-cell level, we conducted receiver operating characteristic (ROC) curve analysis. First, we selected all cells with medium sensor (mTagBFP2 expression between $10^4$ and $10^{5.5}$ a.u.) and high RAS expression (mRuby3 expression between $10^5$ and $10^6$ a.u.). Next, for each sensor, we compared normalized reporter activation between cells expressing mutant RAS variants and those expressing the corresponding wild-type proteins. ROC curves were generated for each sensor by setting thresholds across the range of observed normalized reporter values. At each threshold, we classified cells into true positives (mutant cells above threshold), false positives (wild-type cells above threshold), true negatives (wild-type cells below threshold), and false negatives (mutant cells below threshold). The true positive rate (sensitivity) and false positive rate (1 - specificity) were calculated accordingly. The area under the ROC curve (AUC) was determined using the trapezoidal rule, providing a quantitative measure of each sensor's discriminatory performance. Higher AUC values indicated better sensor performance in distinguishing mutant from wild-type RAS proteins.

##### Sensor characterization in human cancer cell lines

To characterize the performance of sensors in human cancer cell lines, we reverse-transfected 250k cancer cells in a 24-well plate with two separate plasmid mixes encoding (1) binder-cTEVP (100 ng), binder-nTEVP-IRES-mTagBFP2 (135 ng); and (2) iTEVP reporter fused to a CAAX membrane tether (150 ng). After 48h, cells were analyzed by flow cytometry. Median sensor activation was calculated as the ratio of the iTEVP activation signal to the iTEVP expression level (IFP/GFP). Confidence estimates were obtained by bootstrapping (95% confidence intervals; 1,000 bootstrap iterations). We used the same setup to characterize single-protein sensors. Median reporter activity (GFP expression between $10^5$ and $10^{5.5}$ a.u.) was calculated after gating for all sensor-transfected cells. Single-protein sensors were tested using a similar scheme (mTagBFP2 expression between $10^4$ and $10^5$ a.u.).

##### mRNA production

DNA templates containing a 5' T7 promoter sequence followed by the dinucleotide sequence AG were linearized by PCR (Q5 High-Fidelity DNA Polymerase, NEB). A 3' end 120-base-pair poly (A) tail was added using PCR. The linear DNA templates were purified by Ampure beads or gel purification (Zymoclean Gel DNA Recovery Kit, Zymo). mRNA was subsequently produced via *in vitro* transcription (IVT) using NEB's HiScribe T7 High Yield RNA Synthesis Kit. The IVT reaction mix contained 10X Reaction Buffer (1X final), 1 µg DNA template, 4 mM CleanCap AG (TriLink), 2 µl T7 RNA polymerase mix, and the nucleotides ATP, GTP, CTP, and *N1*-Methyl-Pseudouridine-5'-Triphosphate (TriLink) at a final concentration of 5 mM. After an incubation period of 2 hours at 37 °C, DNase I (NEB) was added, and reactions were incubated for another 15 minutes at 37 °C. Finally, mRNA was purified using Zymo's RNA Clean and Concentrator Kit, and concentrations were read out by Nanodrop or Qubit RNA High Sensitivity or Broad Range kits. The concentration-normalized mRNAs were stored at -80 °C.

##### Lipid nanoparticle (LNP) mRNA encapsulation

We adopted a previously described 4-component LNP formulation[21] (4A3-SC8, DOPE, Cholesterol, DMG-PEG) that enables transfection of cells in culture and *in vivo*.

To prepare the lipid stock solutions, a full 25 mg tube of 4A3-SC8 compound was dissolved in 167 µL of pure ethanol to yield a 150 mg/mL stock solution. Separately, 10 mg of DOPE was dissolved in 1.0 mL of pure ethanol to produce a 10 mg/mL stock solution (alternatively, 100 mg in 10 mL ethanol). Similarly, 10 mg of cholesterol was dissolved in 1.0 mL of pure ethanol (or 50 mg in 5 mL ethanol), and 10 mg of DMG-PEG was dissolved in 1.0 mL of ethanol (or 50 mg in 5 mL ethanol), resulting in 10 mg/mL stock solutions for each. A 20 mM working lipid mixture was prepared by combining 6.7 µL of the 4A3-SC8 solution (23.8%, 4.76 mM), 50.7 µL of the DOPE solution (23.8%, 4.76 mM), 52.7 µL of the cholesterol solution (47.6%, 9.52 mM), and 34.2 µL of the DMG-PEG solution (4.8%, 0.96 mM).

The working lipid mixture was used to encapsulate mRNAs in the lipid nanoparticle (LNP) formulation. Initially, the lipid mixture was equilibrated at room temperature for at least 5 minutes and then vortexed at speed 10 for 5 seconds. A lipid mastermix was subsequently prepared by mixing 12 µL of the lipid mixture with 18 µL of 200-proof ethanol, yielding 30 µL per reaction. This mastermix was distributed into individual tubes. Each RNA mixture was prepared by combining 40 µL of RNA solution (250 ng/µL; total 10 µg RNA input) with 32 µL of nuclease-free water and 18 µL of 50 mM citrate buffer, yielding a total volume of 90 µL per reaction. To assemble the LNPs, 30 µL of the lipid mastermix was placed on a Vortex-Genie 2 vortex mixer set at speed level 1. While vortexing, 90 µL of the RNA mixture was rapidly pipetted into the lipid mastermix in a single action, and vortexing was continued for 30 seconds.

The resulting dispersion was incubated at room temperature for 5 minutes, and dialysis was initiated within 15 minutes of mixing. Dialysis was performed using Pur-A-Lyzer Midi 3500 dialysis tubes. Each tube was preconditioned by adding 900 µL of water, incubating for 5 minutes, and then removing the water. Approximately 120 µL of each LNP sample was transferred into the dialysis tubes, which were placed into a styrofoam holder and submerged in 1X PBS within a beaker. Dialysis was conducted either for 1 hour at room temperature or overnight at 4°C. After dialysis, each sample was transferred into an RNase-free 1.5 mL microcentrifuge tube, and the final volume was measured. Samples were adjusted to a total volume of 500 µL by adding the appropriate volume of 1X PBS. All samples were stored at 4°C.

The resulting mRNA-LNP particles exhibited nearly complete transfection efficiency of human MIA PaCa-2 cells and allowed titration of expression over four orders of magnitude (**fig. S14A**). This approach was also compatible with multi-component delivery of separately encapsulated mRNAs (**fig. S14B**).

##### mRNA-LNP transfection of human cell lines

LNPs were reverse-transfected by first adding LNPs to culture plates, followed by the addition of cells on top. To maximize transfection efficiency *in vitro*, mRNA-containing LNPs were pre-complexed with ApoE, a naturally secreted liver protein that binds lipids in vivo and recruits lipid particles to cells expressing the low-density lipoprotein (LDL) receptor.

##### Cell viability assay with CellTiter-Glo

Cell viability was assessed using the CellTiter-Glo (CTG) Luminescent Cell Viability Assay (Promega). Cells were seeded into 96-well plates at 100 µL medium per well and incubated with treatments as described for each experiment. Nunc Edge 2.0 plates were used to minimize edge effects. On the day of the viability assay, plates were removed from the incubator and equilibrated at room temperature for 30 minutes prior to CTG reagent addition. Subsequently, 100 µL of CTG reagent was directly added into each well. Plates were then placed on an orbital shaker in the Promega GloMax instrument and shaken for 2 minutes to facilitate cell lysis and thorough mixing of the reagent. Following shaking, plates were incubated at room temperature for an additional 10 minutes to stabilize the luminescent signal. During this incubation, 180 µL of the mixture from each well was transferred into a white luminescence-compatible plate. Luminescence was measured using the GloMax according to the manufacturer's protocol. Cell viability was calculated by normalizing raw luminescence measurements of treated to untreated samples (at endpoint, unless otherwise noted). For time-course experiments, cell viability was normalized to luminescence measurements of untreated cells at time 0.

##### DepMap RAS-dependence analysis

To analyze whether RAS oncogene dependency varies across cancer cell lines, we analyzed genome-wide CRISPR-Cas9 loss-of-function screening data from the Broad Institute Cancer Dependency Map (DepMap; Project Achilles). Gene-level dependency was quantified by the Chronos gene-effect score (DepMap CRISPRGeneEffect dataset), where 0 denotes no fitness effect and increasingly negative values denote stronger dependency; by DepMap convention, a score near $-1$ corresponds to the median effect of common-essential genes. For each of the 1,186 cancer cell-line models with Chronos data, we tabulated gene-effect scores for KRAS and the core-essential ribosomal gene RPL17 (used as a pan-essential reference), together with model metadata (Oncotree lineage and primary disease) from the corresponding DepMap Model annotation file. Cell lines were classified as KRAS-mutant or KRAS-wild-type by the presence of a KRAS hotspot mutation.

##### KRAS occupancy measurement

KRAS occupancy was measured with a NanoBiT split-luciferase RAS–RBD displacement reporter in which KRAS-G12C was fused to LgBiT and the cRAF RBD to SmBiT. Their association reconstitutes luminescence, which is reduced when the drug or circuit sensor displaces the RBD from KRAS. MIA PaCa-2 cells were transfected with the two reporter constructs (5 ng each) using Lipofectamine 3000 in a 10 cm dish, the medium was changed after 6 h, and cells were re-plated two days later into 96-well plates (20,000 cells per well, 100 µL) and allowed to attach overnight. Sotorasib, RMC-7977, and the sensor$_{v4}$-casp circuit were then added (20 µL into the 100 µL per well) as a serial dilution per treatment in 3 replicate wells per condition. To read out luminescence: the medium was removed and replaced with Nano-Glo Live Cell reagent (furimazine substrate, 20-fold dilution) in Opti-MEM + 10% FBS. Luminescence was read on a Promega Glomax. Viability of MIA PaCa-2 cells was measured in parallel by CellTiter-Glo across matched dose ranges, and viability was plotted against the corresponding occupancy for each treatment.

##### Cell viability assay with flow cytometry

CellTiter-Glo (CTG) provides accurate measurements only within a specific range of cell densities. Thus, for experiments testing varying initial cell densities and their effects on drug versus circuit efficacy, we used flow cytometry to quantify cell viability. Annexin V staining was used to gate and exclude dead cells from analysis.

##### Quantification of pERK and caspase-3 activation

MIA PaCa-2 cells were treated with DMSO, Sotorasib, RMC-7977, or mRNA LNPs for 24 h, then assayed for phospho-ERK and active caspase-3 by intracellular flow cytometry. Cells were stained using a two-step fixation/methanol intracellular protocol. Culture medium was removed and discarded, and the remaining adherent cells were harvested by trypsinization and pelleted. Cells were fixed by adding an equal volume of 2X eBioscience IC Fixation Buffer (~20 min at room temperature, protected from light), permeabilized in ice-cold 90–100% methanol ($\geq$30 min at 2–8 °C), and washed twice in eBioscience Flow Cytometry Staining Buffer. Cells were then co-stained for 30–60 min at room temperature (protected from light) with a directly conjugated active caspase-3 antibody (clone C92-605.rMAb) and an Alexa Fluor 647-conjugated phospho-p44/42 MAPK (Erk1/2) (Thr202/Tyr204) antibody (clone E10) (~1 µL each per 100 µL), washed twice in staining buffer, and analyzed by flow cytometry.

##### Analysis of RAS circuit response to EGF

HEK293 cells (100k cells per well) were seeded in a 24-well plate and treated with epidermal growth factor (EGF) at concentrations of 0, 0.1, 1, 10, and 100 ng/mL for 1 hour. Following treatment, cells were processed for phospho-ERK staining and analyzed by flow cytometry. Specifically, cells were fixed by adding an equal volume (600 µL) of 2X eBioscience™ IC Fixation Buffer directly to each well, followed by gentle vortexing to mix thoroughly. Samples were incubated for 20 minutes at room temperature in the dark. After fixation, cells were scraped thoroughly from each well using a pipette tip, collected, and centrifuged at 600 × g for 5 minutes at room temperature, followed by removal of the supernatant. Cell pellets were then resuspended in the residual volume and permeabilized by adding 200 µL ice-cold 90–100% methanol (HPLC grade), followed by vortexing and incubation for at least 30 minutes on ice. Subsequently, cells were washed by adding 600 µL eBioscience™ Flow Cytometry Staining Buffer, centrifuged at 600 × g for 5 minutes, and the supernatant discarded. The wash step was repeated, leaving approximately 100 µL of residual staining buffer. Cells were stained with 5 µL (0.125 µg) directly conjugated anti-phospho-ERK1/2 antibody (Catalog #12-9109-42) for 30 minutes at room temperature in the dark. After staining, cells underwent two additional washes with 600 µL Flow Cytometry Staining Buffer, each followed by centrifugation at 400–600 × g for 4–5 minutes, with final resuspension in 100 µL of staining buffer for analysis. Finally, cells were analyzed by flow cytometry using the entire remaining volume (100 µL). The fraction of pERK-positive cells was calculated as the fraction of cells with ECD-A signal > $10^3$ a.u. Cell viability was measured as described above.

##### RNA-sequencing

For the transcriptome comparison, MCF10A cells were treated with Sotorasib, RMC-7977, the sensor$_{v4}$-casp circuit, a negative-control LNP (NeoR mRNA), or a constitutively active cell-death effector (caspase-3 + active TEV protease) in triplicate. Treatments were applied for 24h before cells were washed with PBS three times, trypsinized, and resuspended in Zymo DNA/RNA Shield. Sample preparation and sequencing were performed by Plasmidsaurus. Differential expression was assessed with edgeR using a quasi-likelihood F-test; genes were considered differentially expressed at a false-discovery rate below 0.05 and an absolute log2 fold-change of at least 1. Each treatment was compared both to the DMSO vehicle control and to the negative-control LNP, the latter to separate circuit-specific effects from the general response to LNP delivery. Gene-set enrichment of the differentially expressed genes was then performed with Enrichr against the MSigDB Hallmark and Reactome gene-set collections.

##### Drug treatment *in vitro*

Cells were treated with Sotorasib/AMG-510 (10 mM, MedChemExpress, Cat. No.: HY-114277), RMC-7977 (10 mM, MedChemExpress, Cat. No.: HY-156498), or Paclitaxel (10 mM, MedChemExpress, Cat. No.: HY-B0015) at the indicated concentrations per experiment.

##### Time-lapse microscopy

HEK293 and MIA PaCa-2 cell lines were engineered to stably express mRuby3 and GFP, respectively. A co-culture containing 50,000 total cells at a 3:1 ratio of HEK293 to MIA PaCa-2 was seeded into black-walled, glass-bottom 24-well plates (Ibidi) and incubated overnight. The following day, drug or circuit treatments were applied, and plates were imaged using an Olympus automated microscope controlled by MetaMorph software. Time-lapse image acquisition was performed every 2 hours. Images were processed using a custom pipeline in Python. Raw image tiles were first stitched with 10% overlap to reconstruct full fields of view. Stitched images underwent three main processing steps. First, Gaussian filtering (7×7 kernel) was applied to denoise background speckles. Second, image contrast was normalized using percentile-based intensity scaling, with intensities rescaled between the 1st and 99.9th percentiles to account for field-to-field variation. Finally, a dual-threshold segmentation strategy was used for cell detection and quantification. Pixels exceeding the high threshold (20th percentile of the normalized intensity range) were classified as healthy cells and assigned a fixed intensity value of 180. Pixels between the high (20th percentile) and low (15th percentile) thresholds were preserved with their scaled intensities to capture cells with reduced fluorescence signal, indicative of potential cell stress or death. This method enabled consistent detection of both bright and dim cells while preserving biologically relevant signal variation.

##### Sequential resistance selection with drugs and circuit

To assess resistance acquisition, MIA PaCa-2 cells were placed under continuous selection for approximately two months with Sotorasib, RMC-7977, or the sensor$_{v4}$-casp circuit, each at its own concentration that enabled killing of 70% of cells before selection. Each treatment arm was run in three independent replicates per cell line, alongside non-selected control cultures. Cells were maintained in 6-well plates seeded at 60,000 cells per well (2 mL cell suspension plus 200 µL treatment per well) and re-treated approximately every 6 days: at each round, cells were trypsinized, counted (Countess with trypan blue), resuspended to the original seeding density, and re-plated with freshly prepared treatment as on day 1, over ~2 months. At baseline ('no selection') and after selection, each population was challenged in a 96-well format (2,000 cells per well) with a six-point dose range of all three treatments, and viability was read by CellTiter-Glo three days later. Dose-response curves (four-parameter log-logistic) were fit to extract $EC_{50}$ (half-maximum as set by upper and lower asymptote in log-logistic model) and residual viability at saturating dose. Upper asymptote was set to cell viability and lower asymptote to cell viability at the highest treatment concentration (**fig. S38 and S40**). Three-parameter modeling was used as a fallback option when convergence of four-parameter models failed.

##### MCF10A EGF-withdrawal assay

To test how increased RAS expression affects circuit vs inhibitor efficacy, we used the EGF-dependent MCF10A transformation system. MCF10A cells were seeded at 2,500 cells per well (100 µL) in 96-well plates under EGF withdrawal and transfected with a titration of ectopic $KRAS^{G12V}$ mRNA-LNP, along with NeoR mRNA-LNP added as filler to hold the total mRNA-LNP constant at 500 pg/µL for 24 hours. Cells were then treated with a single concentration of the sensor$^{v4}$-casp circuit mRNA-LNPs (300 pg/uL total; 70 pg/uL of sensor + 70 pg/uL of effector) or RMC-7977 (1 nM), across three replicate 96-well plates. KRAS-dependent proliferation was read by CellTiter-Glo three days later.

##### Bypass-signaling resistance assay

To test how the ectopic expression of inhibitor-resistance-inducing genes affects circuit efficacy, we transfected MiaPaCa2 cells with the following genes: NeoR (negative control), CCND1 T286A, YAP1 S127A, SHOC2 S2G, and MEK1 Q56P and analyzed cell viability upon treatment with circuit and

RMC-7977. MiaPaCa-2 cells were seeded at 2,000 cells per well (100 µL) in 96-well plates transfected with a single concentration of ectopic gene mRNA-LNP (300 pg/µL) except for MEK1 Q56P, which was transfected at 150 pg/uL with an additional 150 pg/uL of NeoR to bring the final LNP amount up to 300 pg/uL, for 24 hours. Cells were then treated with sensor$_{v4}$-casp circuit mRNA-LNPs at 0,40, 80, 160, 240, 400 pg/uL (normalized to 300 pg/uL total LNP with NeoR) or RMC-7977 at 0, 0.1, 1, 5, 10, 50 nM. Proliferation was read by CellTiter-Glo three days later. Experiments were performed with three biological replicates.

##### Animal studies

All animal handling, care, and treatment procedures were carried out in accordance with the applicable regulations and guidelines established by the relevant Institutional Animal Care and Use Committee (IACUC). Animals were housed in polycarbonate cages within an environmentally controlled, well-ventilated room maintained at a constant temperature of 20-26°C and a relative humidity of 40-80%. Fluorescent lighting was provided on a 12-hour light/dark cycle.

##### *In vivo* hydrodynamic tail vein injection liver tumor models

FVB/NJ mice (Jackson Laboratory, 001800) were used for HDT tumor models due to their enhanced susceptibility to tumor development. Hydrodynamic transfection was used to introduce transposable vectors expressing mutant NRAS$^{G12V}$, TP53 shRNA, and Sleeping Beauty transposase (SB100) into FVB mice. Mice were injected at 6-8 weeks of age, when their body weights were ~18-25 g. HDT plasmids were suspended at the noted concentrations in 2 mL of saline and administered via tail-vein injection over 7 seconds. A 10:1 mass ratio of combined HDT plasmids to SB100 transposase plasmid was used. In the genetic tumor rescue experiments, PT2-NRAS$^{G12V}$ was used, and 10 ug/mouse of each plasmid was added to the HDT volume. LNP studies substituted PT3-NRAS$^{G12V}$, and 1 ug of this plasmid/mouse was used due to increased tumor induction efficiency. In the *in vivo* mRNA-LNP experiments, LNPs were dosed at 1.5 mg total RNA per kg body weight dissolved in 0.2 mL PBS via the lateral tail vein at the noted time points. For the KRAS-driven model, the NRAS$^{G12V}$ transposon was replaced with a KRAS$^{G12C}$ transposon, retaining the shTP53 and SB100 components and the HDT procedure above. For the *in vivo* delivery experiment, GFP was added to the shTP53 plasmid to label transformed clones. To assess LNP delivery to tumor tissue, mice received mRNA-LNPs encoding a nuclear-localized tdTomato reporter. Mice were sacrificed 12 h later and livers processed for fluorescence imaging. Treatment schedules varied by experiment and are given in the corresponding text and figures.

##### *In vivo* pharmacodynamics analysis

To verify tumor-specific sensor activation *in vivo*, we used a membrane-tethered tdTomato reporter engineered to translocate to the nucleus upon TEV protease activity. mRNA-LNPs encoding sensor$_{v4}$ together with this reporter were delivered IV to KRAS$^{G12C}$ tumor-bearing mice on day 10 post-HDT. A construct encoding constitutively active TEV protease served as a positive control. Healthy (non-HDT) mice received the same LNPs as a negative control. Livers were collected 24 h later, sectioned, and imaged to score nuclear vs membrane tdTomato localization in tumor vs adjacent normal cells.

##### Liver and body weight measurements

Animals were euthanized by cervical dislocation under isoflurane anesthesia. Livers were removed, weighed, and imaged at the time of sacrifice. Liver/body weight ratios were calculated as follows: (whole liver weight)/(intact body weight) × 100. Extracted livers were imaged (anterior and posterior views), and corresponding images were used to assess the surface tumor burden of each animal. More specifically, tumor nodules were annotated manually in a blinded fashion. Surface tumor burden was calculated as accumulated surface tumor area over anterior and posterior liver views.

##### Liver surface tumor area annotation and masking

Liver images were manually annotated for nodules by an independent annotator with no involvement in the study. The annotation process was blinded: image files were assigned numeric identifiers, and the annotator was not provided with any information regarding experimental conditions. Tumors were manually segmented using NimbusImage on a total of 284 images of mouse livers. Each image received three separate masks: (1) “white nodes” for tumors that appeared lighter than surrounding tissue, (2) “black nodes” for tumors that appeared darker, and (3) a whole-liver mask outlining the total liver area. Only abnormalities larger than approximately 1 mm were included. Both raised masses and regions of distinct discoloration were classified as nodules. Non-tumor features such as fat deposits, blood vessels, bubbles, and specular reflections were excluded from annotation.

##### Kaplan-Meier curve

Tumors were induced in FVB mice as described above, with or without the noted circuit plasmids. Tumor development was allowed to proceed until reaching any of the three following clinical endpoints: a body condition score of 1, difficulty breathing, or decreased motility resulting in inability to obtain food or water, at which point mice were immediately euthanized at the recommendation of veterinary staff. The log-rank test was used to determine if differences in survival were significant.

##### SW-1573 flank xenograft model

SW-1573 cells were implanted subcutaneously into both flanks, yielding two independent tumors per animal. Mice were shaved and prepared one day before implantation. Once tumors reached approximately 100–200 $mm^3$ (day 16), animals were randomized by tumor size into cohorts of similar mean tumor volume, and treatment began the same day. Animals received vehicle control (PBS), an LNP negative control, daily oral RMC-7977 (10 mg/kg by oral gavage), or the LNP-delivered 12VC1-based sensor-GSDMA pyroptosis circuit. The two LNP arms (circuit and LNP control) were administered by intratumoral injection every three days, and control animals received an equal volume of vehicle or control LNP on the same schedule. Tumor volume was measured by caliper (length and width of each flank tumor). Tumors were harvested on day 25. Tumor growth was analyzed with a linear mixed model (LMM) accounting for repeated measurements and two flank tumors per mouse: log(tumor volume) ~ treatment x day + (1 | mouse). Pairwise treatment comparisons were performed on (i) estimated tumor volume at endpoint and (ii) estimated log-volume growth rates over time; raw p-values within each set of six pairwise contrasts were adjusted using the Benjamini-Hochberg procedure. Model assumptions were assessed using residual and random-effect diagnostic plots (**fig. S45**).

##### LNP formulation and physical characterization for mouse studies

The following lipids were dissolved in 100% ethanol at a 15:30:15:3:7 molar ratio: 4A3-SC8 lipid (synthesized in-house), cholesterol (Sigma Aldrich Cat. No. C3045), DOPE (Avanti Polar Lipids Cat. No. 850725), DMG-PEG-2000 (Avanti Polar Lipids Cat. No. 880151), and DOTAP (Avanti, 890890P) with a total lipid:RNA mass ratio of 40:1. RNA was dissolved in 10 mM citrate buffer (pH 4.5) at a 3:1 v/v aqueous:organic phase ratio. LNPs were formed by microfluidic mixing of the lipid and RNA solutions using a Precision Nanosystems NanoAssemblr Benchtop Instrument, in accordance with the manufacturer’s protocol. LNPs were dialyzed in PBS overnight at 4°C and stored at 4°C for up to 72 hours. Particle size and dispersity were measured by dynamic light scattering (DLS) using a Malvern Zetasizer DLS instrument.

#### Software

The following software was used in this study:

Data collection: AlphaFold 3 (AlphaFold Server), Colabfold (v1.5.2), RFdiffusion (v1.1.0), ProteinMPNN-FastRelax (dl_binder_design v1.0.0), MetaMorph (version 6.2.6)

Data analysis: PyMol (version 2.5.4), ChimeraX (version 1.7.1), FlowJo (version 10.10.0), SnapGene (version 8.0.2), Geneious (version 2023.0.4), Bases2Fastq (Element Biosciences, https://github.com/Elembio/bases2fastq-dx), QuPath (0.5.1), NimbusImage, conda (v4.14.0), python (v3.12.3), bioconductor-biomart (v2.58.0), bioconductor-biostrings (v2.70.1), bioconductor-deseq2 (v1.42.0), cairo (v1.18.0), imagemagick (v7.1.1_33), ipykernel (v6.29.5), ipython (v8.26.0), jupyter_server (v2.14.2), jupyterlab (v4.2.1), r-base (v4.3.3), r-cowplot (v1.1.3), r-dplyr (v1.1.4), r-essentials (v4.3), r-ggplot2 (v3.5.1), r-ggpubr (v0.6.0), r-ggrepel (v0.9.5), r-jsonlite (v1.8.8), r-magick (v2.8.3), r-rcolorbrewer (v1.1_3), r-tidyverse (v2.0.0), pandas (v2.2.3), matplotlib (v3.10.1), numpy (v2.2.4), scipy (v1.15.2), scikit-learn (v1.6.1), opencv-python (v4.7.0.72), scikit-image (v0.20.0), pillow (v9.5.0), Claude, Cursor

#### Statistics and reproducibility

Unless otherwise noted, all experiments were conducted with three to four technical or biological replicates.

#### Supplementary Tables

**Table S1: Statistical analysis of tumor prevention experiment (Fig. 3).** Liver-to-body-weight ratios (LW/BW) and tumor surface area were compared across conditions using pairwise Mann-Whitney U tests and adjusted for multiple testing using the Benjamini-Hochberg procedure.

| Condition | Comparison | Metric | p-value | Adjusted p-value |
|---|---|---|---|---|
| PBS | $\text{sensor}_{v1}$-casp | LW/BW | $2.98 \times 10^{-3}$ | $2.98 \times 10^{-3}$ |
| PBS | $\text{sensor}_{v1}$-casp | Tumor surface area | $4.12 \times 10^{-3}$ | $4.12 \times 10^{-3}$ |
| PBS | $\text{sensor}_{v1}$-amp-casp | LW/BW | $1.18 \times 10^{-7}$ | $3.54 \times 10^{-7}$ |
| PBS | $\text{sensor}_{v1}$-amp-casp | Tumor surface area | $1.96 \times 10^{-6}$ | $3.20 \times 10^{-6}$ |
| $\text{sensor}_{v1}$-casp | $\text{sensor}_{v1}$-amp-casp | LW/BW | $1.87 \times 10^{-5}$ | $2.80 \times 10^{-5}$ |
| $\text{sensor}_{v1}$-casp | $\text{sensor}_{v1}$-amp-casp | Tumor surface area | $2.13 \times 10^{-6}$ | $3.20 \times 10^{-6}$ |

**Table S2: Statistical analysis of NRAS tumor suppression experiment (Fig. 4B).** Liver-to-body-weight ratios (LW/BW) and tumor surface area were compared between PBS- and $\text{sensor}_{v1}$-amp-GSDM-treated groups using a pairwise Mann-Whitney U test.

| Condition | Comparison | Metric | p-value | Adjusted p-value |
|---|---|---|---|---|
| PBS | $\text{sensor}_{v1}$-amp-GSDM | LW/BW | 0.000788 | NA |
| PBS | $\text{sensor}_{v1}$-amp-GSDM | Tumor surface area | 0.000551 | NA |

**Table S3: Statistical analysis of NRAS tumor suppression experiment with apoptosis and pyroptosis circuits (Fig. 4D).** Liver-to-body-weight ratios (LW/BW) were compared across conditions using pairwise Mann-Whitney U tests and adjusted for multiple testing using the Benjamini-Hochberg procedure.

| Condition | Comparison | Metric | p-value | Adjusted p-value |
|---|---|---|---|---|
| PBS | TEVP-LNP | LW/BW | $2.42 \times 10^{-2}$ | 0.0290 |
| PBS | $\text{sensor}_{v4}$-casp | LW/BW | $2.22 \times 10^{-5}$ | $6.66 \times 10^{-5}$ |
| PBS | $\text{sensor}_{v4}$-gsdm | LW/BW | $8.88 \times 10^{-6}$ | $5.33 \times 10^{-5}$ |
| TEVP-LNP | $\text{sensor}_{v4}$-casp | LW/BW | $6.81 \times 10^{-3}$ | 0.0102 |
| TEVP-LNP | $\text{sensor}_{v4}$-gsdm | LW/BW | $6.81 \times 10^{-3}$ | 0.0102 |
| $\text{sensor}_{v4}$-casp | $\text{sensor}_{v4}$-gsdm | LW/BW | 0.713 | 0.713 |

**Table S4: Statistical analysis of KRAS tumor treatment experiment (Fig. 4F).** Liver-to-body-weight ratios (LW/BW) were compared between PBS and $\text{sensor}_{v4}$-gsdm-treated groups using a pairwise Mann-Whitney U test.

| Condition | Comparison | Metric | p-value | Adjusted p-value |
|---|---|---|---|---|
| PBS | $\text{sensor}_{v4}$-gsdm | LW/BW | 0.0014 (**) | NA |

**Table S5: Statistical analysis of NRAS tumor suppression experiment with RMC-7977 (fig. S42).** Liver-to-body-weight ratios (LW/BW) were compared between PBS- and RMC-7977-treated groups using a pairwise Mann-Whitney U test.

| Condition | Comparison | Metric | p-value | Adjusted p-value |
|---|---|---|---|---|
| PBS | RMC-7977 | LW/BW | 0.0221 (*) | NA |

**Table S6: Statistical analysis of CDX treatment experiment (Fig. 6E).** Pairwise treatment comparisons were performed based on endpoint tumor volumes and estimated log-volume growth rates from a linear mixed model (LMM, see Methods). p-values were adjusted for multiple testing using the Benjamini-Hochberg procedure.

| **Condition** | **Comparison** | **Metric** | **p-value** | **Adjusted p-value** |
|---|---|---|---|---|
| PBS | LNP ctrl. | Growth trajectory (LMM) | 0.627 (ns) | 0.747 (ns) |
| PBS | RMC-7977 | Growth trajectory (LMM) | 0.430 (ns) | 0.645 (ns) |
| PBS | $Sensor_{v4}$-GSDM | Growth trajectory (LMM) | 0.00163 (**) | 0.00980 (**) |
| LNP ctrl. | RMC-7977 | Growth trajectory (LMM) | 0.747 (ns) | 0.747 (ns) |
| LNP ctrl. | $Sensor_{v4}$-GSDM | Growth trajectory (LMM) | 0.00445 (**) | 0.0133 (*) |
| RMC-7977 | $Sensor_{v4}$-GSDM | Growth trajectory (LMM) | 0.0113 (*) | 0.0225 (*) |
| PBS | LNP ctrl. | Endpoint tumor volume (LMM) | 0.960 (ns) | 0.960 (ns) |
| PBS | RMC-7977 | Endpoint tumor volume (LMM) | 0.394 (ns) | 0.473 (ns) |
| PBS | $Sensor_{v4}$-GSDM | Endpoint tumor volume (LMM) | $3.20 \times 10^{-5}$ (****) | $9.69 \times 10^{-5}$ (****) |
| LNP ctrl. | RMC-7977 | Endpoint tumor volume (LMM) | 0.392 (ns) | 0.473 (ns) |
| LNP ctrl. | $Sensor_{v4}$-GSDM | Endpoint tumor volume (LMM) | $1.22 \times 10^{-5}$ (****) | $7.35 \times 10^{-5}$ (****) |
| RMC-7977 | $Sensor_{v4}$-GSDM | Endpoint tumor volume (LMM) | $3.36 \times 10^{-4}$ (***) | $6.72 \times 10^{-4}$ (***) |

#### Supplementary Figures

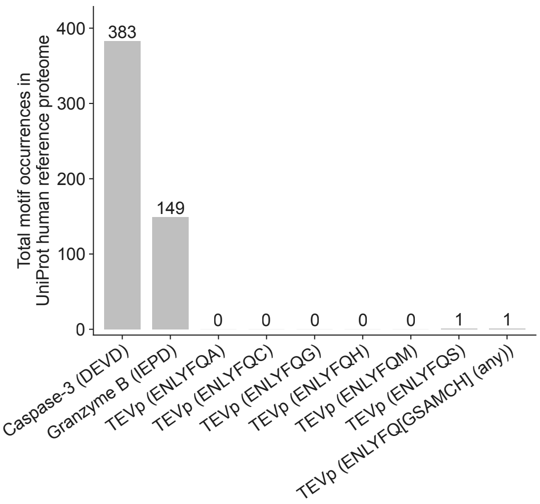


**Fig. S1. Abundance of protease cleavage motifs in the human proteome.** Bar chart showing the total number of exact cleavage-motif occurrences for caspase-3 (DEVD), granzyme B (IEPD), and TEV protease (ENLYFQ followed by A, C, G, H, M, or S individually, or any residue in the set [GSAMCHI]) identified by scanning the UniProt human reference proteome. Endogenous protease motifs for caspase-3 and granzyme B occur frequently (383 and 149, respectively), whereas TEV protease recognition sequences are nearly absent: five of six individual ENLYFQX variants yield zero occurrences, one (ENLYFQS) yields a single match, and the union of all tested P1' residues yields only one occurrence total. These data confirm that TEV protease cleavage activity is orthogonal to the human proteome.

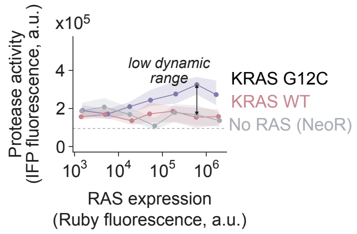


**Fig. S2. The RAF1-RBD-based sensor (sensor$_{v1}$) insufficiently discriminated between mutant and wild-type RAS.** Protease activity (IFP fluorescence, a.u.) of sensor$_{v1}$ is plotted as a function of RAS expression level (Ruby fluorescence, a.u.) for overexpressed KRAS$^{G12C}$, KRAS$^{WT}$, and a no-RAS control (NeoR). The sensor exhibited low dynamic range between mutant and wild-type KRAS responses, limiting its ability to detect endogenous mutant RAS in cancer cell lines. Shaded regions represent bootstrap 95% confidence intervals of the median (Methods).

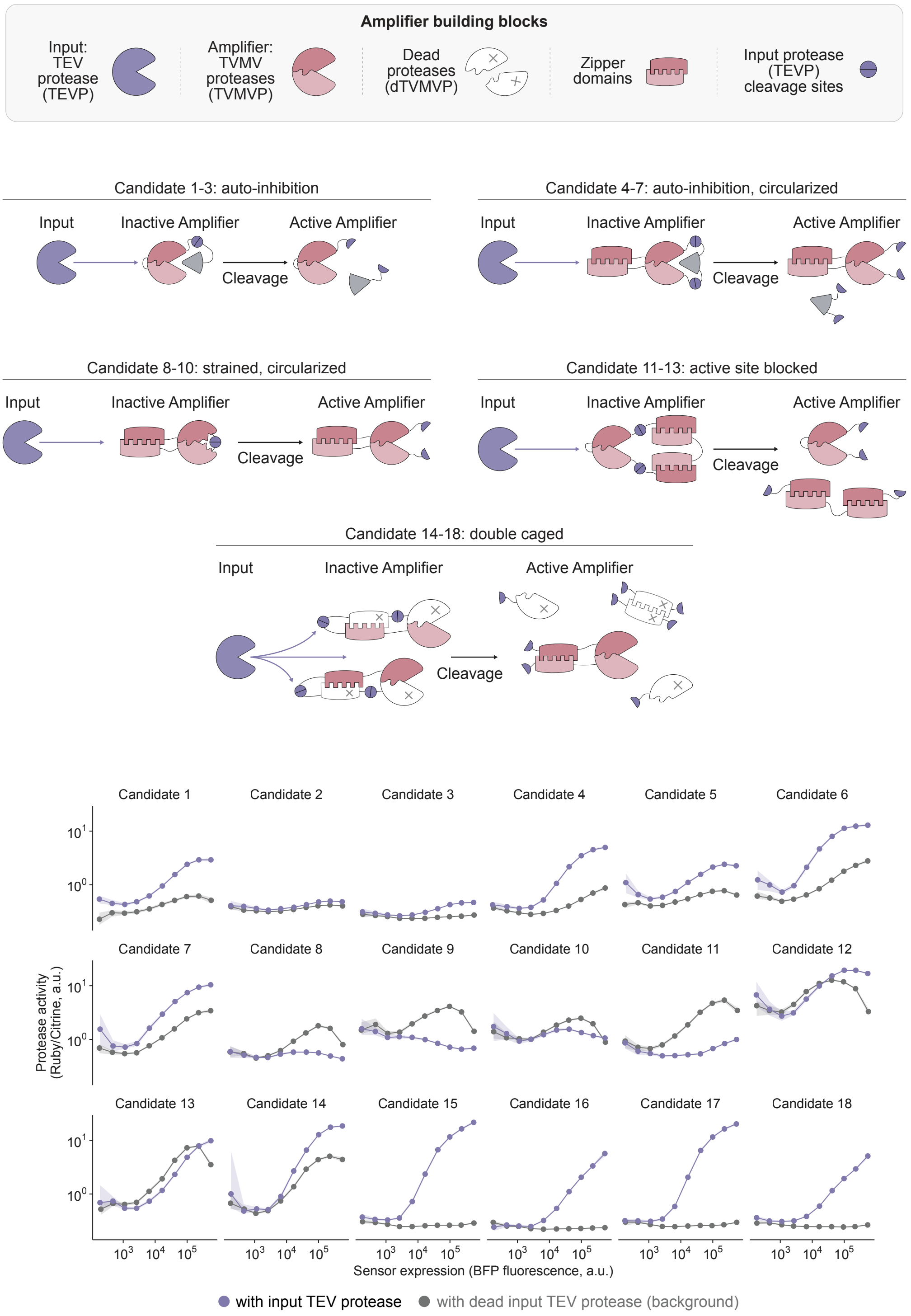


**Fig. S3. Characterization of candidate protease-activatable protease amplifiers.** Top: schematics of five amplifier design strategies for engineering TEV protease (TEVP)-activatable tobacco vein mottling virus protease (TVMVP) amplifiers. Building blocks include active TVMVP, catalytically dead TVMVP (dTVMVP), coiled-coil zipper domains, and TEVP cleavage sites. Candidates 1 to 3 use auto-inhibition by fusion of each TVMVP half to its catalytically dead complementary half. Candidates 4 to 7 extend this design with backbone circularization. Candidates 8 to 10 use strained, circularized architectures. Candidates 11 to 13 block the active site with a removable domain. Candidates 14 to 18 use a dual-caged design in which each split TVMVP half is fused to its dead complementary half and to a coiled-coil dimerization domain, with TEVP cleavage sites enabling release and competitive displacement of the caging domains. Bottom: protease activity (Ruby/Citrine ratio, a.u.) as a function of sensor expression (BFP fluorescence, a.u.) for each of the 18 candidate amplifiers, measured in the presence of active input TEV protease or catalytically dead input TEV protease (background control). Most designs provided little amplification or de-amplification; the dual-caged candidates (14 to 18) showed the greatest separation between active and dead input conditions, consistent with the design selected for further development (Fig. 1B).

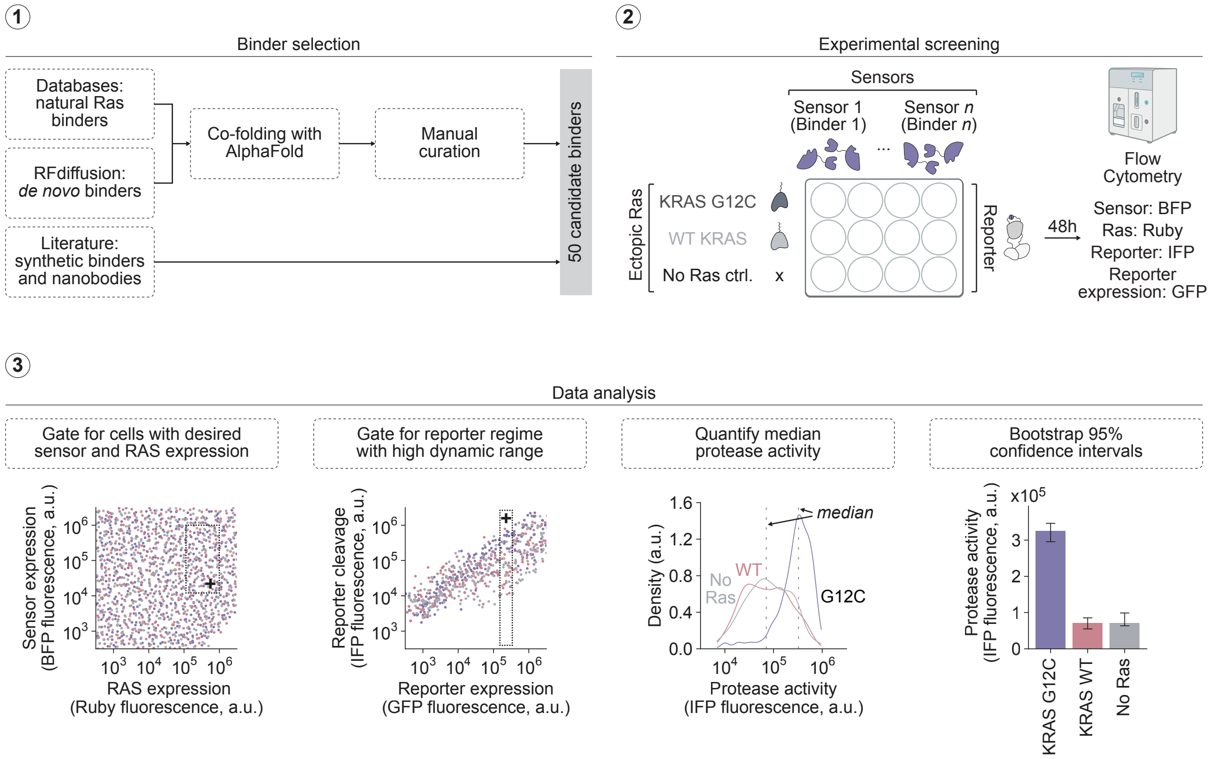


**Fig. S4. Sensor screening workflow and flow-cytometry quantification.** (Step 1) Binder selection: candidate RAS-binding domains were compiled from databases of natural RAS binders, *de novo* designed binders (RFdiffusion), and previously published synthetic binders and nanobodies. Candidates were co-folded with AlphaFold and manually curated to yield 50 candidate binders. (Step 2) Experimental screening: each candidate binding domain was fused to complementary split-TEV protease domains to create a sensor. HEK293 cells were polytransfected with each sensor (marked by BFP), ectopic RAS variants (KRAS$^{G12C}$, KRAS$^{WT}$, or no-RAS control; marked by mRuby3), and a protease-activatable fluorescent reporter (IFP, with GFP marking reporter expression). Cells were analyzed by flow cytometry 48 h after transfection. (Step 3) Data analysis: cells were gated for desired sensor (BFP) and RAS (mRuby3) expression levels, then gated for the reporter expression regime (GFP) with the highest dynamic range. Median protease activity (IFP fluorescence) was quantified for each RAS condition, and bootstrap 95% confidence intervals of the median were computed (Methods). Error bars represent bootstrap 95% confidence intervals of the median.

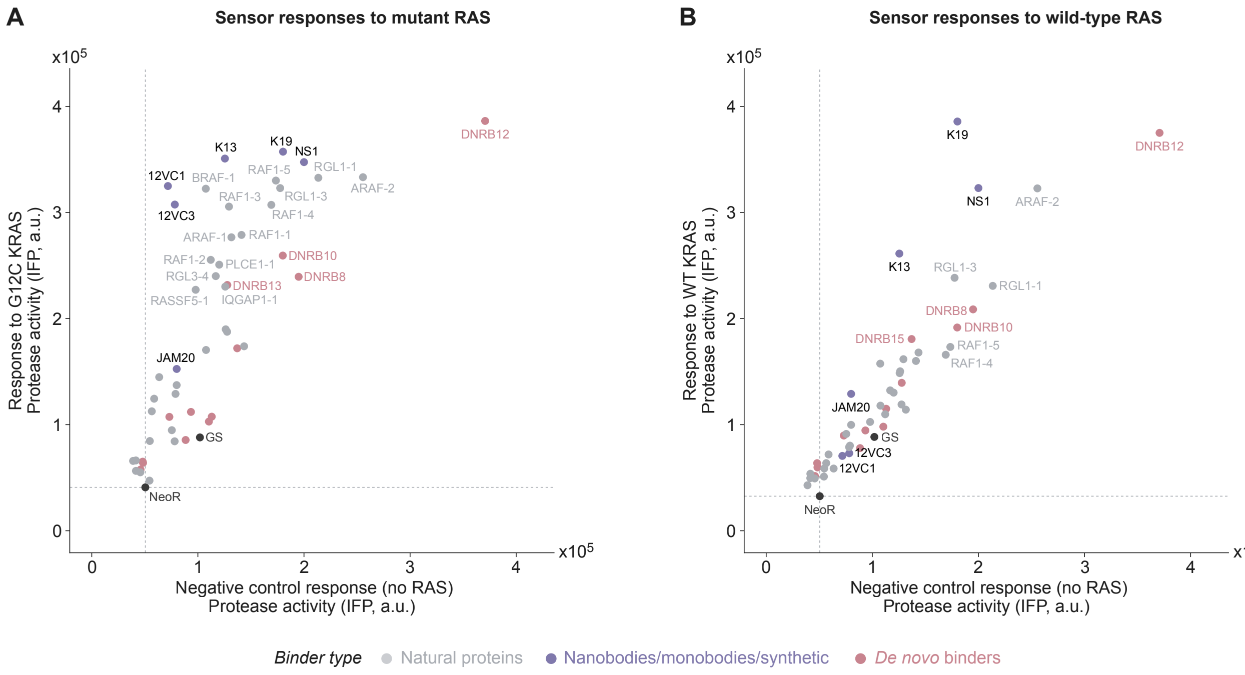


**Fig. S5. RAS binder screen identifies mutant-specific sensors. (A)** Library screen enables characterization of sensor protease activity against mutant $KRAS^{G12C}$ compared to a no-RAS negative control. **(B)** Library screen enables characterization of sensor protease activity against wild-type KRAS compared to a no-RAS negative control. **(A)**, **(B)** Each data point represents the median protease reporter activation (IFP, a.u.) for a candidate sensor in the high RAS and medium sensor expression regime (Methods). Sensors are color-coded by the category of RAS-binding domain used. Text labels denote the binding domain used for each sensor. GS denotes the GSGSGS negative control peptide used as a binder. Dashed lines indicate the baseline protease activity of the NeoR negative control (no sensor).

A

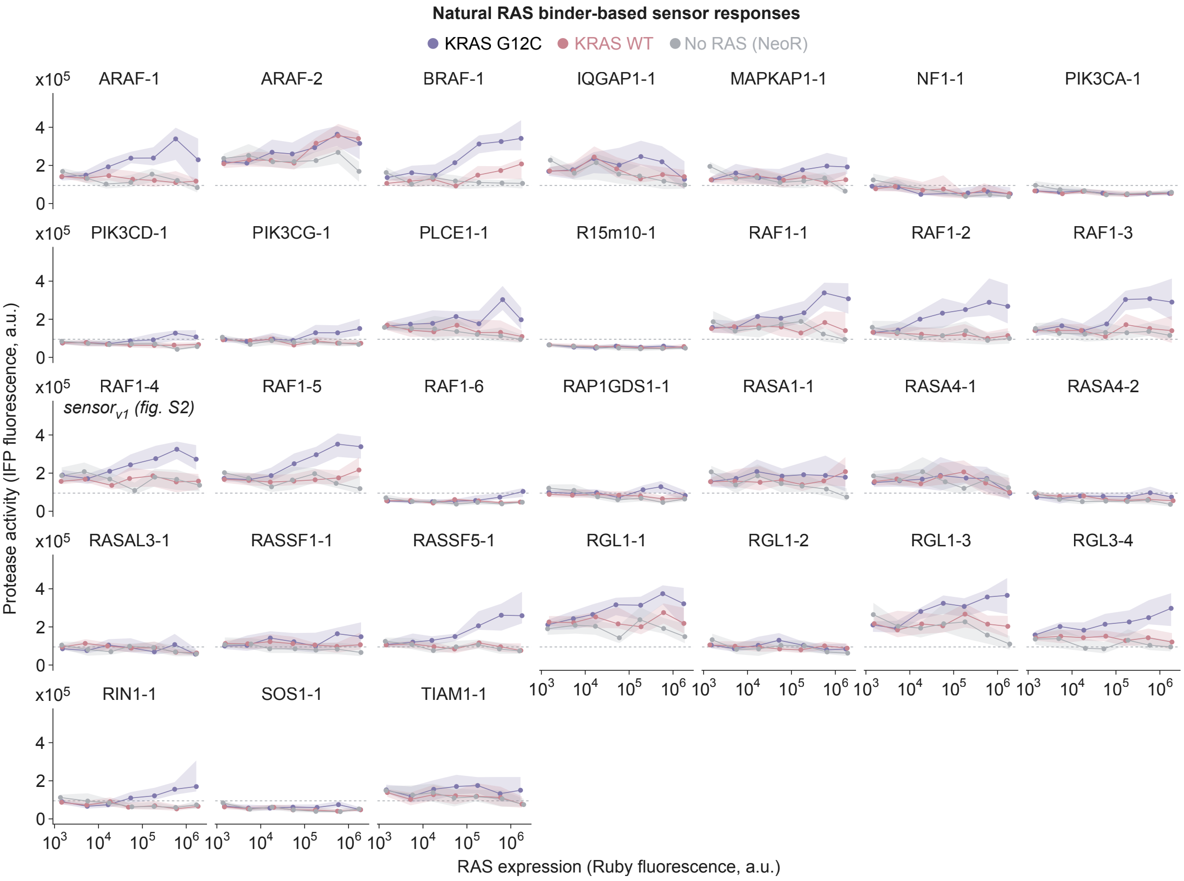


B

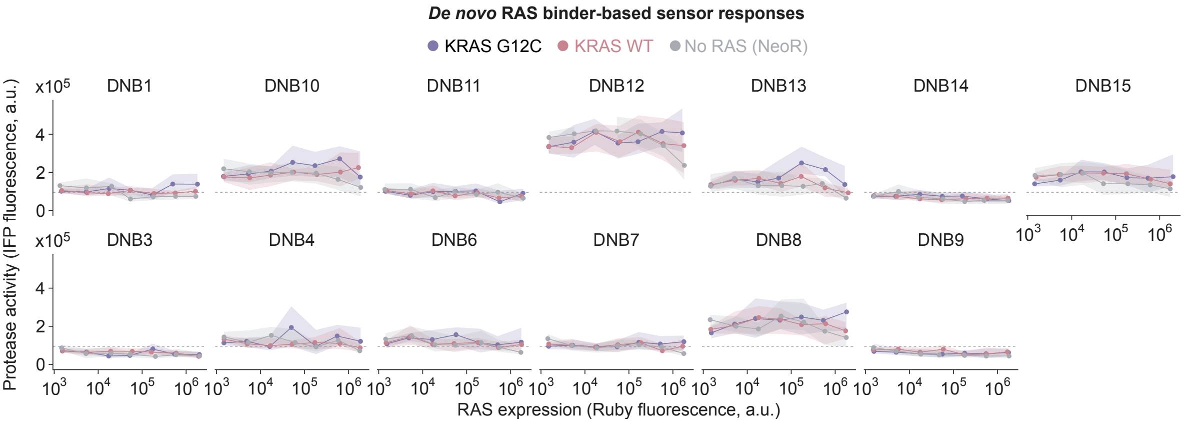


C

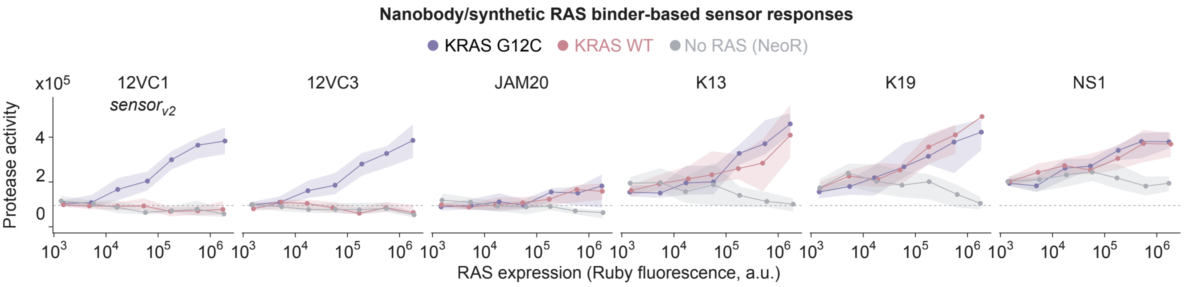


**Fig. S6. RAS expression dependency of individual sensor responses. (A)** Sensor responses of natural RAS binder-based sensors. Headings specify binding domains used for each sensor. RAF1-4 corresponds to $sensor_{v1}$. **(B)** Responses of sensors incorporating *de novo* RAS binders (DNB). **(C)** Responses of sensors utilizing nanobodies, monobodies, or other synthetic RAS binders. **(A)**-**(C)** Each subpanel plots median protease activity (IFP fluorescence, a.u.) as a function of RAS expression (Ruby fluorescence, a.u.) for $KRAS^{G12C}$, $KRAS^{WT}$, and a no-RAS control (NeoR). Data points represent median reporter activation in the intermediate sensor expression regime; dashed green line represents the activity of a negative control sensor containing GSGSGS peptides as binders. Shaded regions depict bootstrap 95% confidence intervals of the median (1,000 iterations). Sensor performance was evaluated following the experimental workflow described in Fig. 1C.

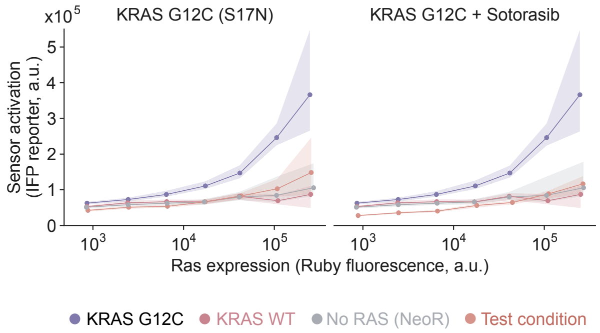


**Fig. S7. $Sensor_{v2}$ is RAS-dependent.** Sensor activation (IFP reporter, a.u.) is plotted as a function of RAS expression level (Ruby fluorescence, a.u.) for $KRAS^{G12C}$, $KRAS^{WT}$, no RAS (NeoR negative control), and the indicated test condition. Left: co-expression of the dominant-negative $KRAS^{G12C}$ S17N mutation reduces sensor activation to near-background levels, comparable to $KRAS^{WT}$ and no-RAS controls. Right: treatment of $KRAS^{G12C}$-expressing cells with Sotorasib similarly abrogates sensor activation. Data points represent median sensor activation. Shaded areas depict 95% confidence intervals from bootstrapping (1,000 iterations). Sensor performance was evaluated following the experimental workflow outlined in Fig. 1C.

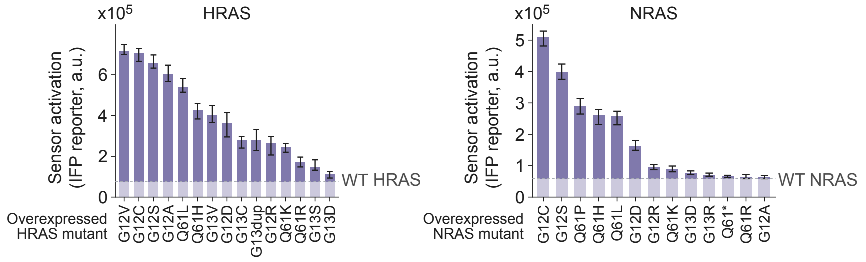


**Fig. S8. Sensor$_{v2}$ responds to diverse oncogenic HRAS and NRAS mutations.** Sensor activation (IFP reporter, a.u.) was measured in HEK293 cells co-transfected with the sensor$_{v2}$ construct, the indicated overexpressed HRAS (left) or NRAS (right) mutant variants, and a protease-activatable fluorescent reporter, following the experimental workflow described in Fig. 1C. Bars represent median reporter activation after gating for high RAS expression (Methods). The WT HRAS and WT NRAS reference bars indicate sensor activation in the presence of overexpressed wild-type RAS. Companion KRAS mutant data are shown in Fig. 1E. Error bars represent bootstrap 95% confidence intervals of the median (Methods).

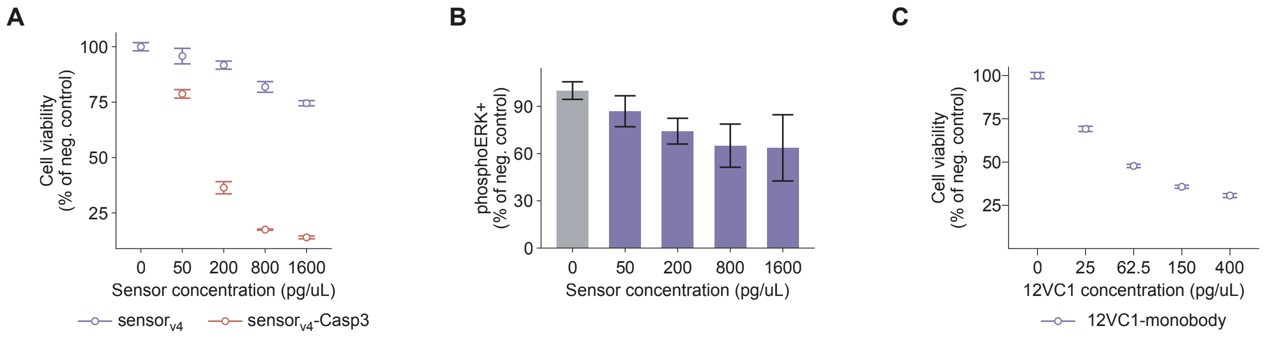


**Fig. S9. Effects of individual circuit components on cell viability and RAS signaling. (A)** Cell viability (% of negative control) as a function of LNP-delivered mRNA concentration for $sensor_{v4}$ alone or $sensor_{v4}$-casp. $Sensor_{v4}$ alone is largely tolerated, whereas the complete $sensor_{v4}$-casp circuit induces potent, dose-dependent cell death. **(B)** Percentage of phospho-ERK-positive (pERK+) cells (% of negative control) as a function of sensor concentration (pg/µL). Increasing sensor concentration leads to dose-dependent reduction of pERK. **(C)** Cell viability (% of negative control) as a function of 12VC1 monobody mRNA concentration (pg/µL). The 12VC1 monobody delivered alone reduces viability in a dose-dependent manner. Error bars in **(A)**, **(B)**, and **(C)** represent mean ± s.d.

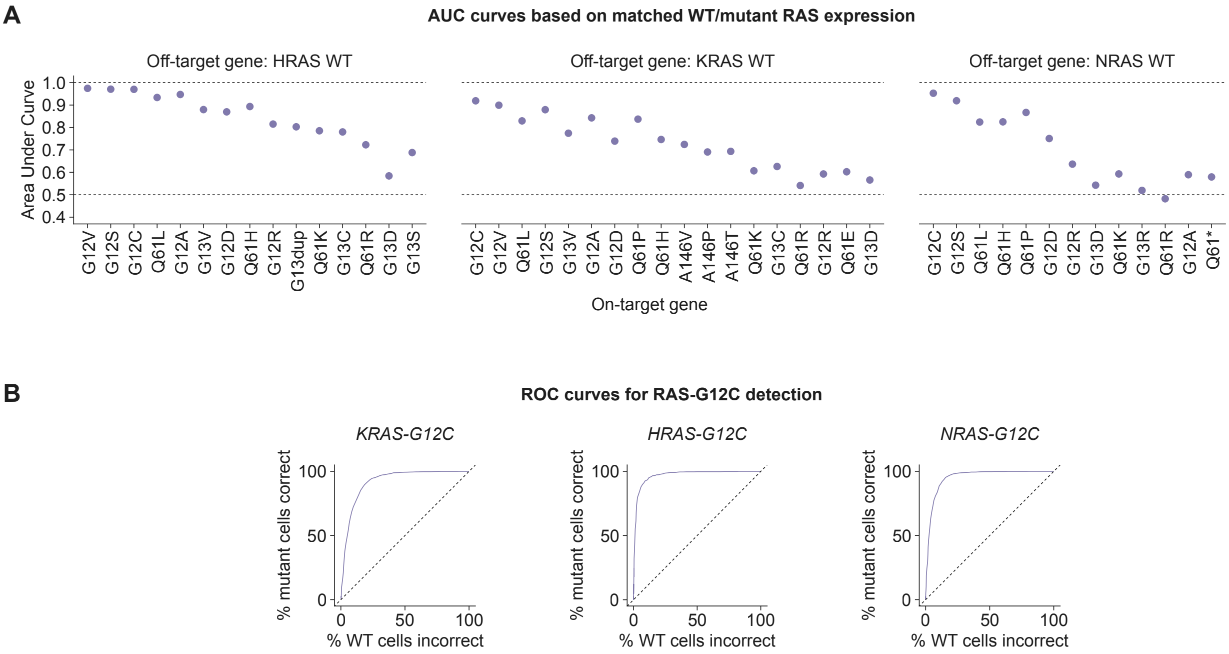


**Fig. S10. Sensor$_{v2}$ accurately classifies the RAS mutational status of single cells. (A)** Area under curve (AUC) values from receiver operating characteristic (ROC) analysis of sensor$_{v2}$ discriminating mutant from wild-type RAS-expressing cells at matched RAS expression levels. Each dot represents one oncogenic mutation. Subpanels show classification performance against HRAS$^{WT}$ (left), KRAS$^{WT}$ (center), and NRAS$^{WT}$ (right) as the off-target (wild-type) reference. Dotted lines indicate perfect classification (AUC = 1.0) and random classifier performance (AUC = 0.5). **(B)** ROC curves for G12C detection across all three RAS isoforms: KRAS$^{G12C}$ (left), HRAS$^{G12C}$ (center), and NRAS$^{G12C}$ (right). The y-axis shows the true positive rate (% mutant cells correctly classified) and the x-axis shows the false positive rate (% wild-type cells incorrectly classified), computed by varying the protease reporter activation threshold at matched RAS expression levels. Diagonal dotted line represents the performance of a random classifier.

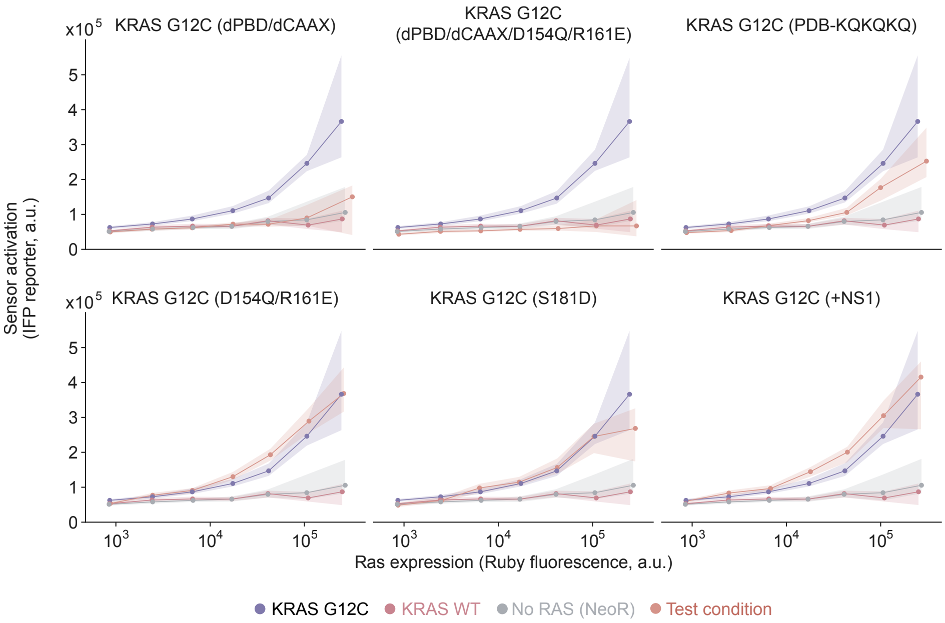


**Fig. S11. RAS membrane localization, but not clustering, is required for sensor$_{v2}$ activation.** Sensor$_{v2}$ response was measured against KRAS$^{G12C}$ variants carrying mutations that disrupt membrane targeting or dimerization/clustering. Each panel plots sensor activation (IFP reporter, a.u.) as a function of RAS expression (Ruby fluorescence, a.u.) for KRAS$^{G12C}$, KRAS$^{WT}$, no RAS (NeoR), and the indicated test condition. Deletion of the polybasic domain and CAAX motif (dPBD/dCAAX), combined deletion with dimer-interface mutations (dPBD/dCAAX/D154Q/R161E), and replacement of the polybasic domain with a weakened sequence (PBD-KQKQKQ) abolished or reduced sensor activity. In contrast, disrupting the dimer interface alone (D154Q/R161E), introducing a clustering-modifying mutation (S181D), or co-expressing the nanoclustering inhibitor NS1 had little effect on sensor sensitivity. Data points represent median sensor activation; shaded regions depict 95% confidence intervals from bootstrapping (1,000 iterations). Sensor performance was evaluated following the experimental workflow outlined in Fig. 1C.

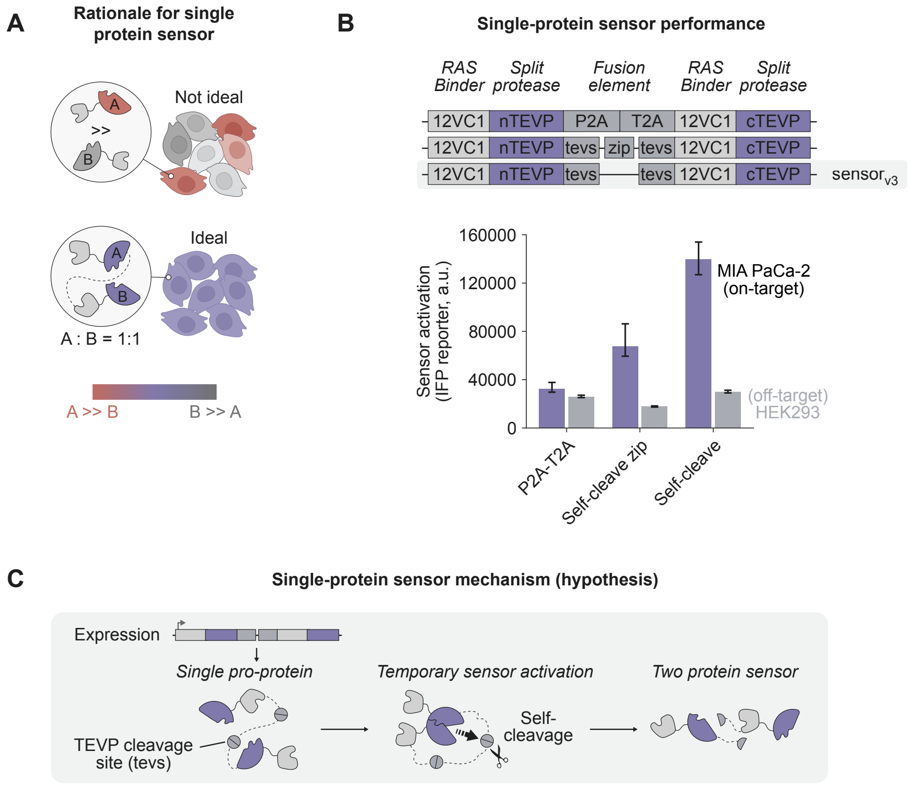


**Fig. S12. Self-cleaving polyprotein design improves sensor activation. (A)** Rationale for a single-protein sensor. Expressing both sensor halves from a single open reading frame ensures 1:1 stoichiometry between the two split-protease components, which is expected to improve sensor performance compared to designs where one component is expressed in excess. **(B)** Single-protein sensor performance. Top: three polyprotein architectures linking the two 12VC1-based sensor halves (12VC1-nTEVP and 12VC1-cTEVP) via different fusion elements: P2A-T2A (ribosomal skipping peptides), self-cleave zip (TEV cleavage sites flanking a coiled-coil zipper domain), and self-cleave (TEV cleavage sites without zipper; sensor$_{v3}$). Bottom: median sensor activation (IFP reporter, a.u.) measured in on-target MIA PaCa-2 (KRAS$^{G12C}$) and off-target HEK293 (KRAS$^{WT}$) cells. The self-cleave design (sensor$_{v3}$) achieves the highest on-target activation while maintaining low off-target signal. Error bars represent 95% confidence intervals from bootstrapping (1,000 iterations). **(C)** Proposed mechanism for the self-cleaving single-protein sensor. A single pro-protein is expressed and folds; the embedded TEV protease transiently activates upon RAS-dependent co-localization, cleaving the internal TEV cleavage sites (tevs) to liberate the two sensor halves as independent proteins that then function as a conventional two-component split-protease sensor.

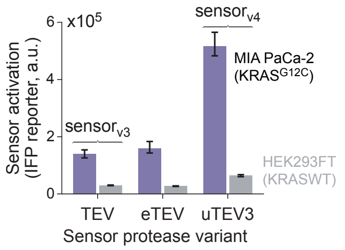


**Fig. S13. Catalytically enhanced TEV protease (uTEV3) improves sensor performance.** Bar plot shows median sensor activation (IFP reporter, a.u.) for sensor$_{v3}$ (containing wild-type TEV or eTEV protease) and sensor$_{v4}$ (containing uTEV3 protease) in on-target MIA PaCa-2 cells (KRAS$^{G12C}$) and off-target HEK293FT cells (KRAS$^{WT}$). Incorporation of the uTEV3 protease variant markedly increases sensor activation in KRAS-mutant cells while maintaining low background in wild-type cells. Error bars denote bootstrap 95% confidence intervals of the median.

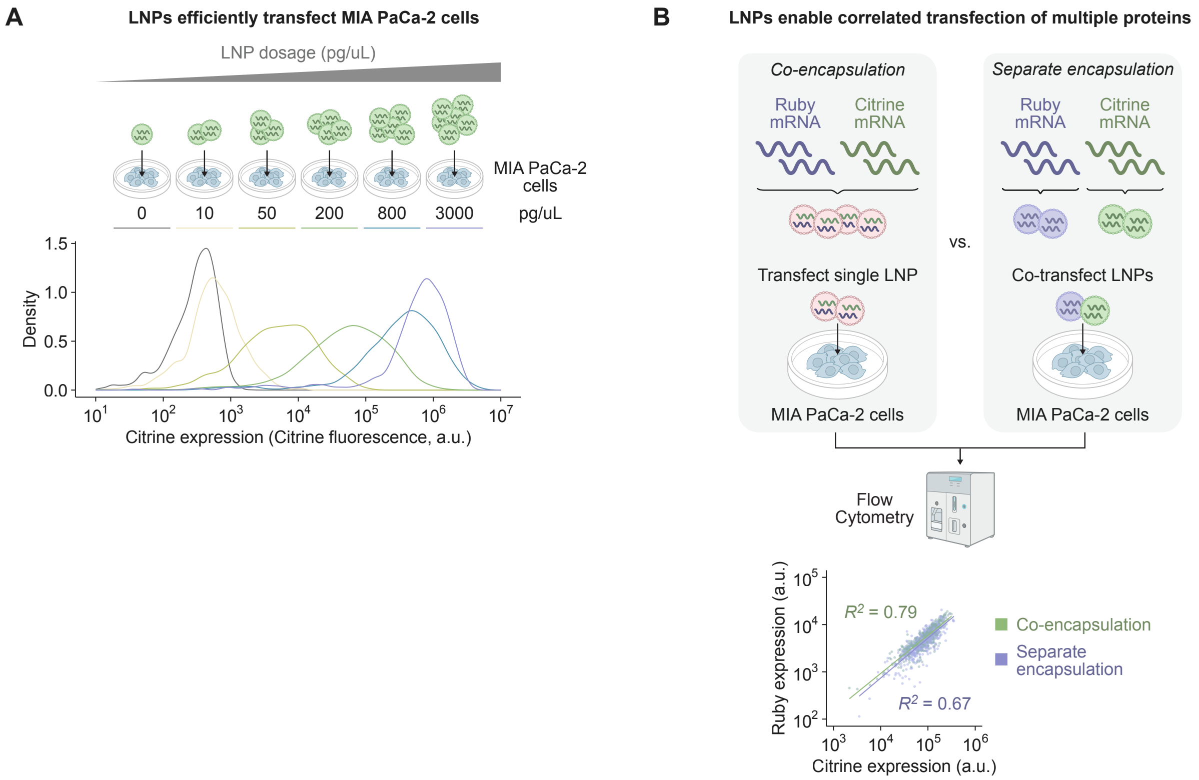


**Fig. S14. LNPs efficiently deliver mRNA to cells *in vitro* in a dose-dependent fashion. (A)** MIA PaCa-2 ($KRAS^{G12C}$) cells were treated with Citrine-encoding mRNA encapsulated in LNPs (4A3-SC8/DOPE formulation) at the indicated concentrations (0, 10, 50, 200, 800, and 3000 pg/µL). Density distributions of Citrine fluorescence (a.u.) measured by flow cytometry show dose-dependent increases in expression. **(B)** Co-encapsulation of two mRNAs (mRuby3 and Citrine) within a single LNP produces correlated expression of both proteins in individual cells. MIA PaCa-2 cells were transfected with LNPs containing co-encapsulated mRuby3 and Citrine mRNAs or with separately encapsulated LNPs co-delivered to the same wells, and analyzed by flow cytometry. Scatter plot shows single-cell Citrine versus mRuby3 expression (a.u.) for co-encapsulation ($R^2$ = 0.79) and separate encapsulation ($R^2$ = 0.67).

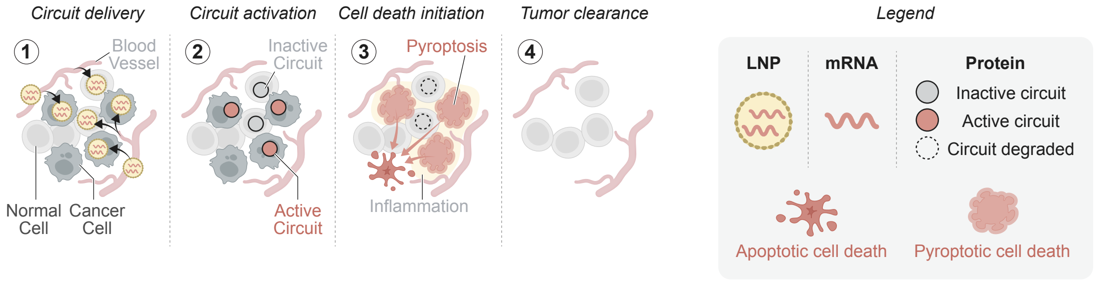


**Fig. S15. Schematic of pyroptosis-mediated bystander killing by therapeutic circuits.** (1) Therapeutic circuits encoding a RAS sensor and a pyroptosis effector are delivered as mRNA in lipid nanoparticles (LNPs) to both normal and cancer cells via the vasculature. (2) In normal cells, the circuit remains inactive and is degraded; in RAS-mutant cancer cells, sensor activation by clustered mutant RAS reconstitutes the protease. (3) Active circuits trigger pyroptotic cell death, releasing inflammatory signals that recruit immune cells and promote bystander killing of neighboring untransfected tumor cells. (4) Pyroptosis-induced inflammation enables tumor clearance even when only a fraction of cancer cells are directly transfected. Legend defines symbols for LNPs, mRNA, and protein states (inactive circuit, active circuit, circuit degraded) as well as apoptotic and pyroptotic cell death.

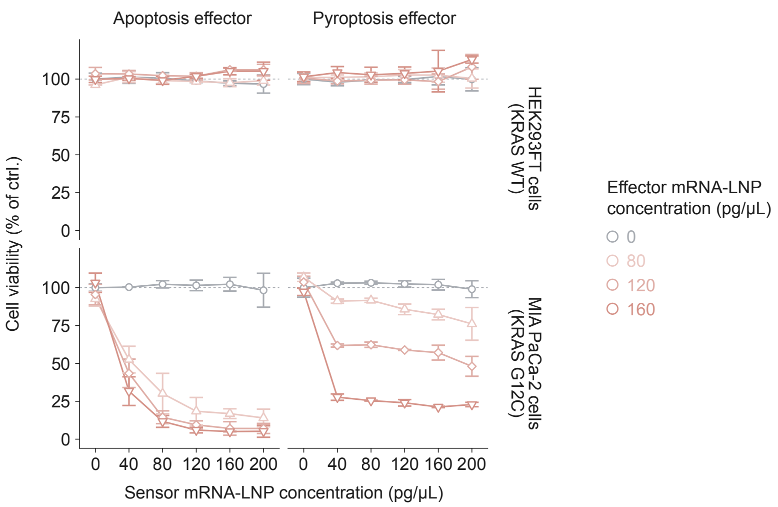


**Fig. S16. Dose-response characterization of apoptosis and pyroptosis circuits across sensor and effector concentrations.** Cell viability (% of control) of HEK293FT ($KRAS^{WT}$; top row) and MIA PaCa-2 ($KRAS^{G12C}$; bottom row) cells treated with $sensor_{v4}$-casp (apoptosis effector; left column) or $sensor_{v4}$-GSDM (pyroptosis effector; right column) mRNA-LNP circuits. Sensor mRNA-LNP concentration was titrated along the x-axis (0 to 200 pg/μL) at four fixed effector mRNA-LNP concentrations (0, 80, 120, and 160 pg/μL). Both circuits induced potent, dose-dependent killing of RAS-mutant MIA PaCa-2 cells while sparing RAS-wild-type HEK293FT cells across all tested concentrations. Cell viability was measured by CellTiter-Glo 3-5 days post-treatment. Error bars represent mean ± s.d. (n = 2 replicates). Dashed line indicates 100% viability.

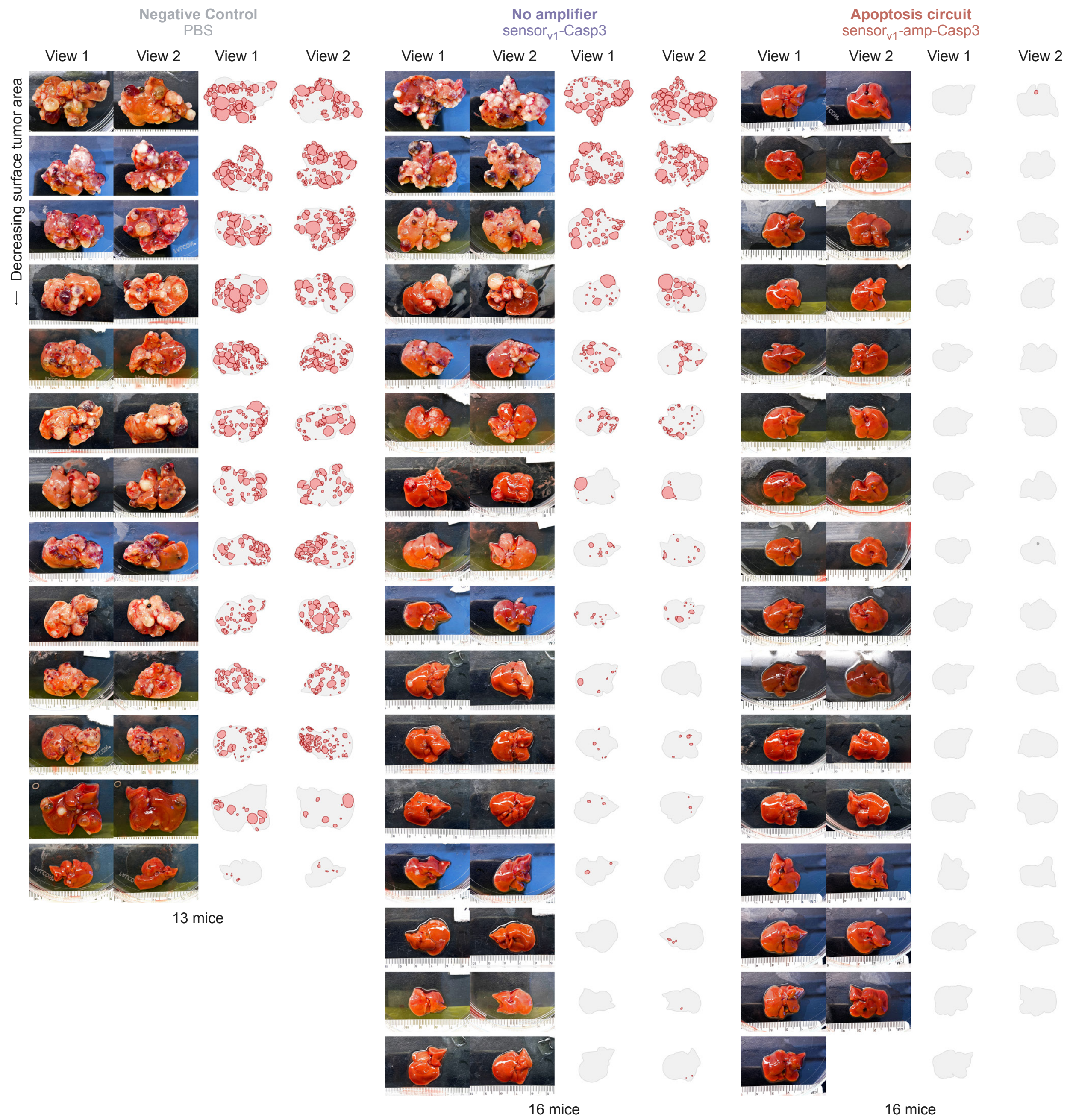


**Fig. S17. Extended liver images from *in vivo* tumor-prevention experiment.** Photographs and corresponding manually annotated surface tumor maps for all mice in the tumor-prevention cohort (Fig. 3B). Tumors were induced by hydrodynamic tail-vein injection (HDT) of plasmids encoding NRAS$^{G12V}$, shTP53, and SB100 transposase into female mice. Livers were collected 36-40 days after tumor induction. Three treatment groups are shown: negative control (PBS; n = 13), sensor$_{v1}$-casp without amplifier (n = 16), and sensor$_{v1}$-amp-casp apoptosis circuit with amplifier (n = 16). For each animal, two views of the excised liver are displayed alongside matched images with manually annotated surface tumor nodules (red). Animals are ordered by decreasing surface tumor area within each group.

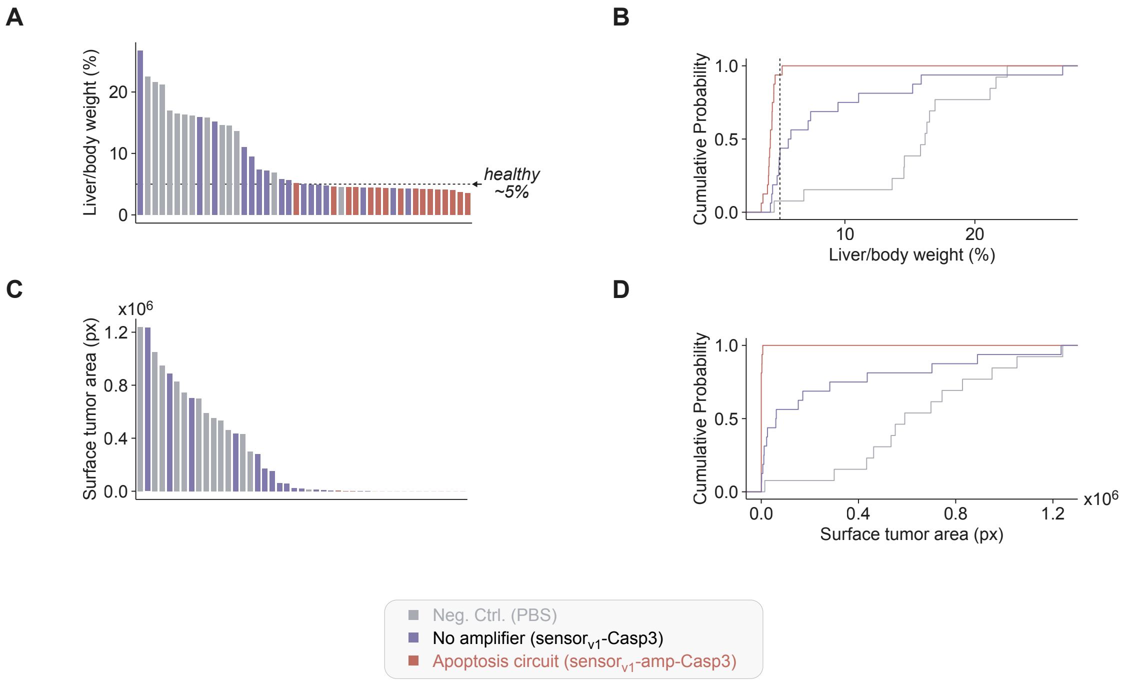


**Fig. S18. Extended quantification of tumor burden for *in vivo* tumor-prevention model. (A)** Waterfall plot of liver-to-body weight ratio (%) for individual mice treated with PBS (negative control), sensor$_{v1}$-casp (no amplifier), or sensor$_{v1}$-amp-casp (apoptosis circuit), sorted by decreasing tumor burden. The dashed line indicates the healthy liver-to-body weight ratio (~5%). **(B)** Cumulative probability distribution of liver-to-body weight ratios for the same three treatment groups. The dashed vertical line marks the ~5% healthy reference. **(C)** Waterfall plot of surface tumor area (px) for individual mice, sorted by decreasing tumor burden. **(D)** Cumulative probability distribution of surface tumor area for the same three treatment groups.

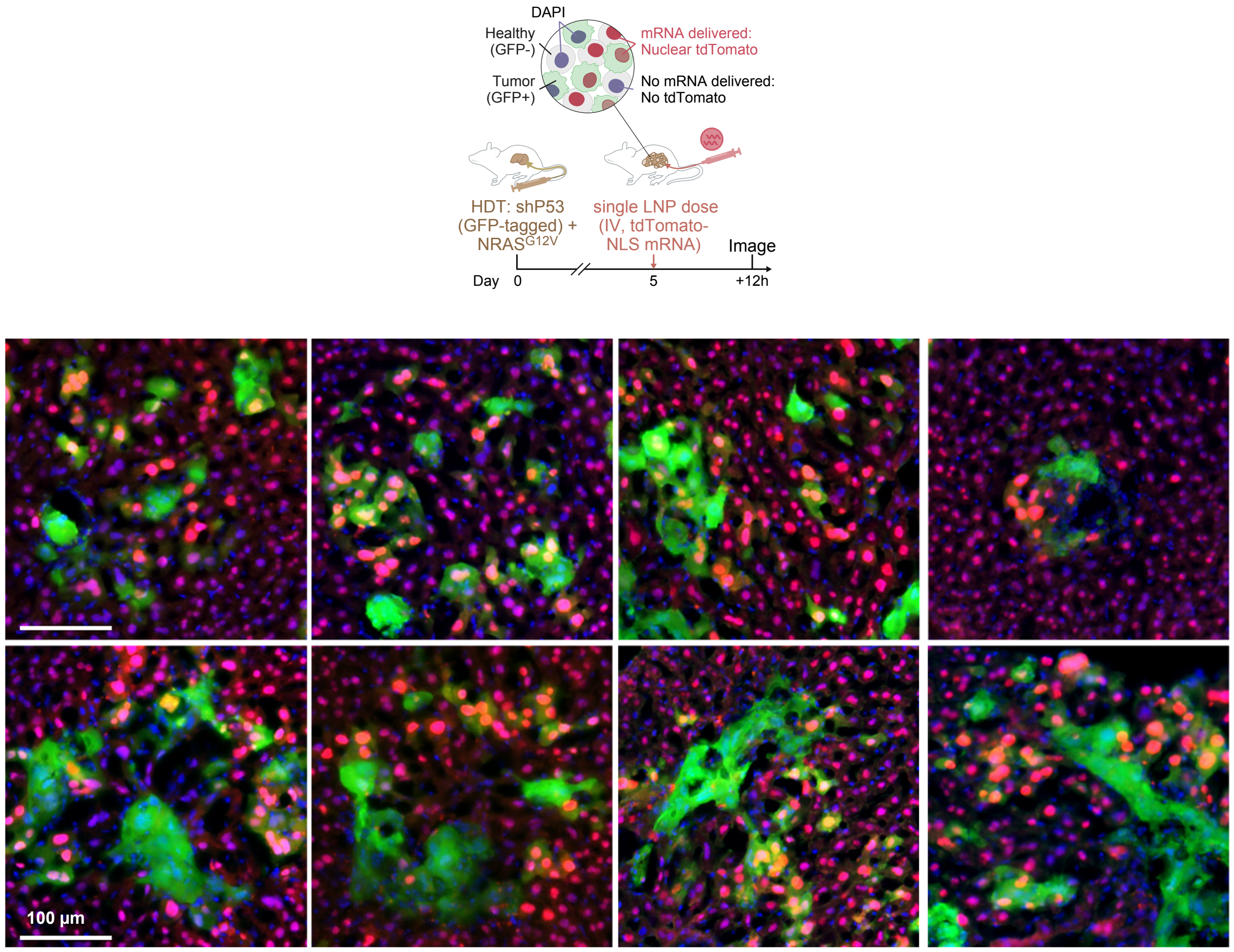


**Fig. S19. Systemically delivered LNPs access nascent liver tumor cells *in vivo*.** Top: schematic of the experimental workflow. Tumors were initiated on day 0 by hydrodynamic tail-vein injection (HDT) of GFP-tagged shP53 and $NRAS^{G12V}$ constructs with SB100 transposase; on day 5, a single intravenous dose of LNPs encapsulating nuclear-localized tdTomato (tdTomato-NLS) mRNA was administered, and livers were collected 12 hours later for fluorescence imaging. Healthy hepatocytes (GFP-negative) that received LNP cargo display nuclear tdTomato fluorescence (red); tumor cells are marked by GFP (green); DAPI marks nuclei (blue). Bottom: representative fluorescence micrographs of liver sections from tumor-bearing mice. Scale bars, 100 µm.

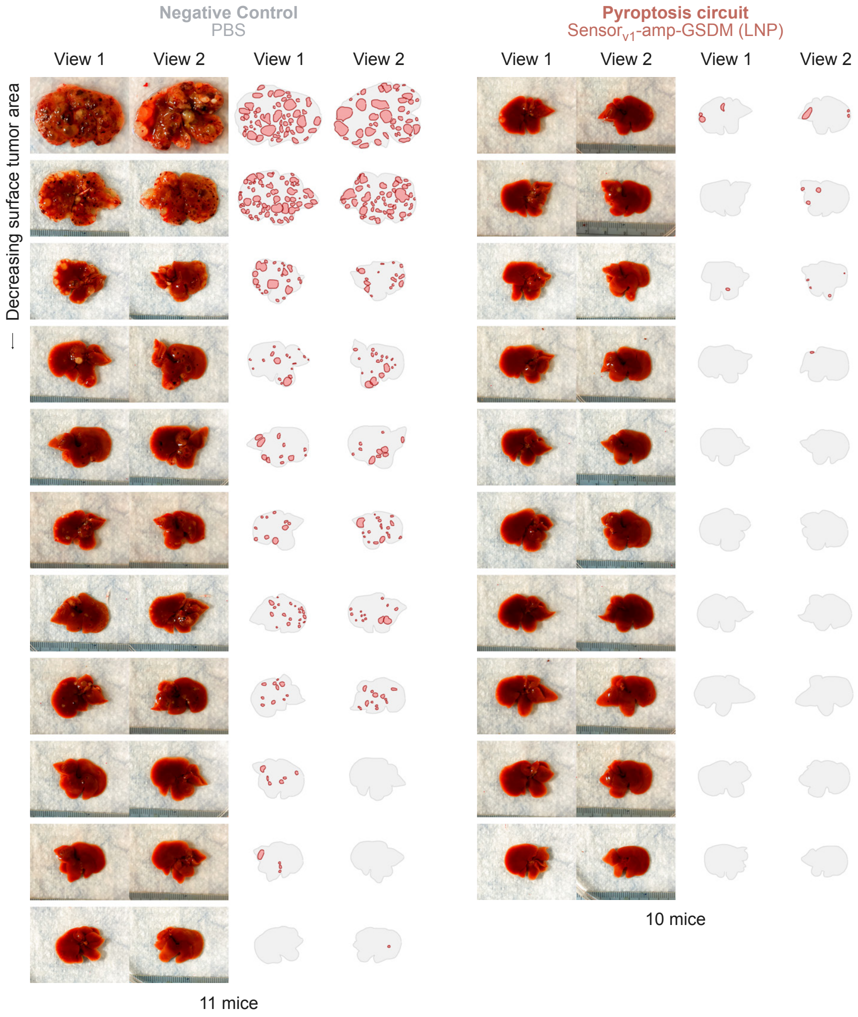


**Fig. S20. Pyroptosis circuit suppresses nascent NRAS-driven liver tumors (extended).** Representative liver photographs and corresponding blinded surface-tumor annotations for all individual mice from the experiment shown in Fig. 4B. Left: negative control (PBS, n = 11 mice). Right: pyroptosis circuit (sensor$_{v1}$-amp-gsdm, n = 10 mice). For each mouse, two views of the liver are shown alongside manually annotated surface-tumor nodules (red outlines). Mice are ordered by decreasing surface tumor area from top to bottom.

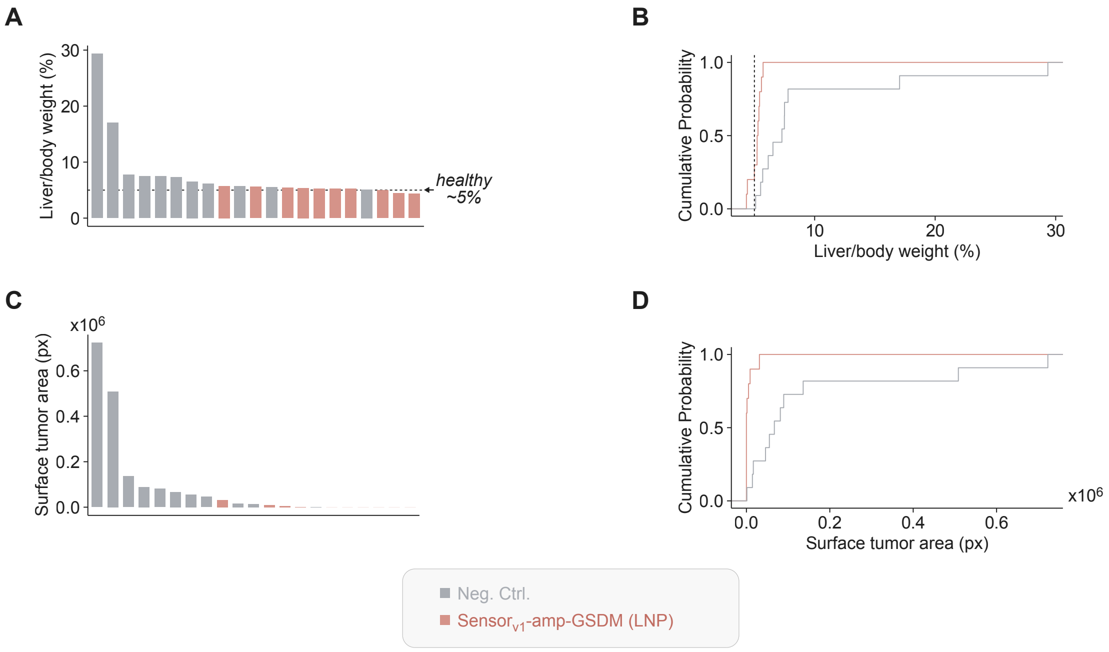


**Fig. S21. Quantification of liver tumor burden for fig. S20 (extended). (A)** Waterfall plot of liver-to-body weight ratio (%) for individual mice; dashed line indicates the healthy baseline (~5%). **(B)** Cumulative probability distribution of liver-to-body weight ratios for the same treatment groups; dashed vertical line marks the healthy baseline (~5%). **(C)** Waterfall plot of surface tumor area (px) for individual mice. **(D)** Cumulative probability distribution of surface tumor area for the two groups. Representative liver images for all animals are shown in fig. S20. Experimental details are shown in Fig. 4B.

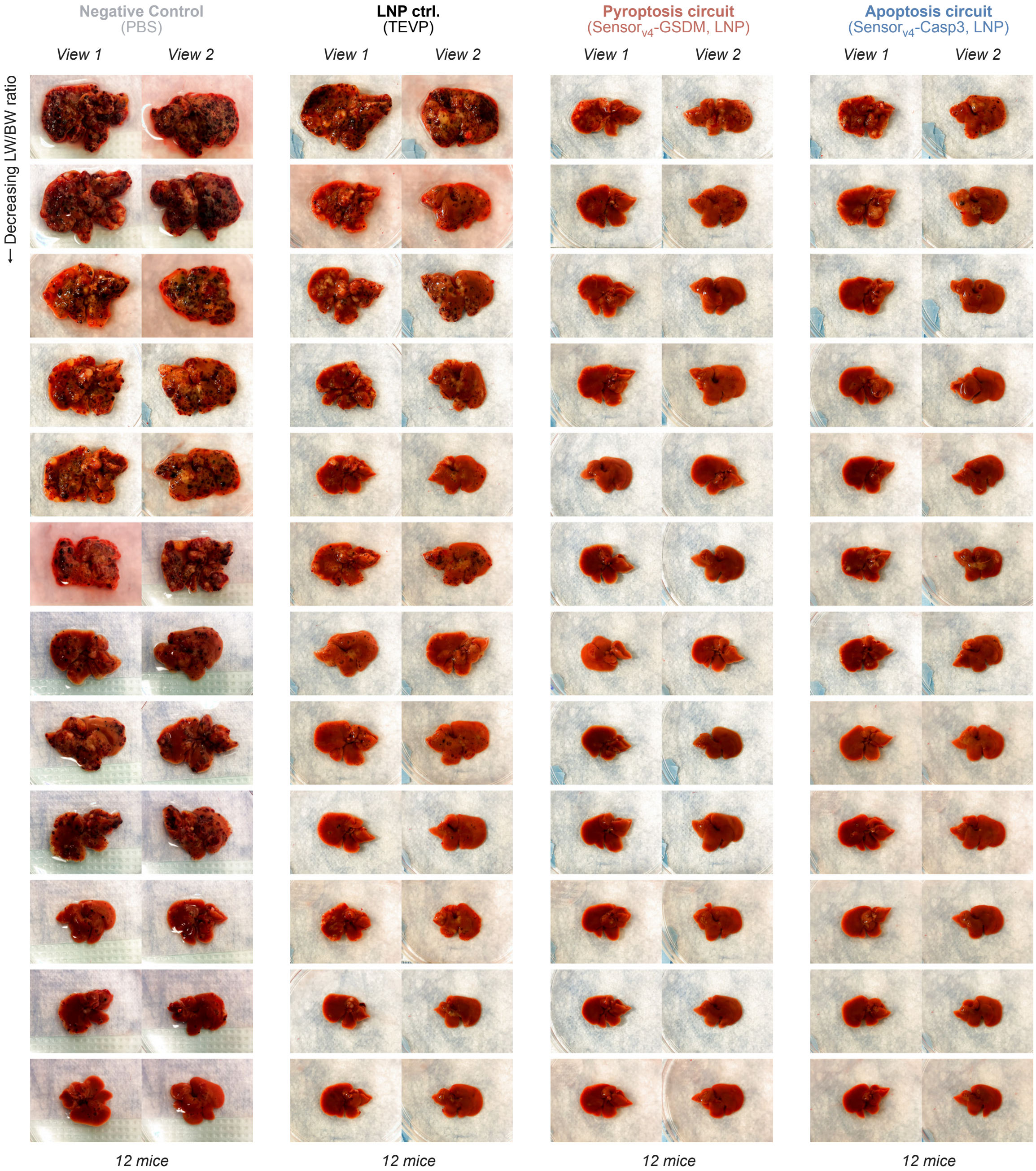


**Fig. S22. Apoptosis and pyroptosis circuits suppress NRAS-driven liver tumors (extended).** Livers from all mice in each treatment group (n = 12 female mice per condition) are shown from two viewing angles (View 1 and View 2), sorted by decreasing liver-to-body weight (LW/BW) ratio. Treatment groups: PBS (negative control), LNP control encoding split TEV protease halves without RAS-binding domains or a functional effector (TEVP), sensor$_{v4}$-GSDM (pyroptosis circuit), and sensor$_{v4}$-casp (apoptosis circuit). Both circuits used sensor$_{v4}$ paired with the indicated effector, delivered without an amplifier. Experimental details are shown in Fig. 4D.

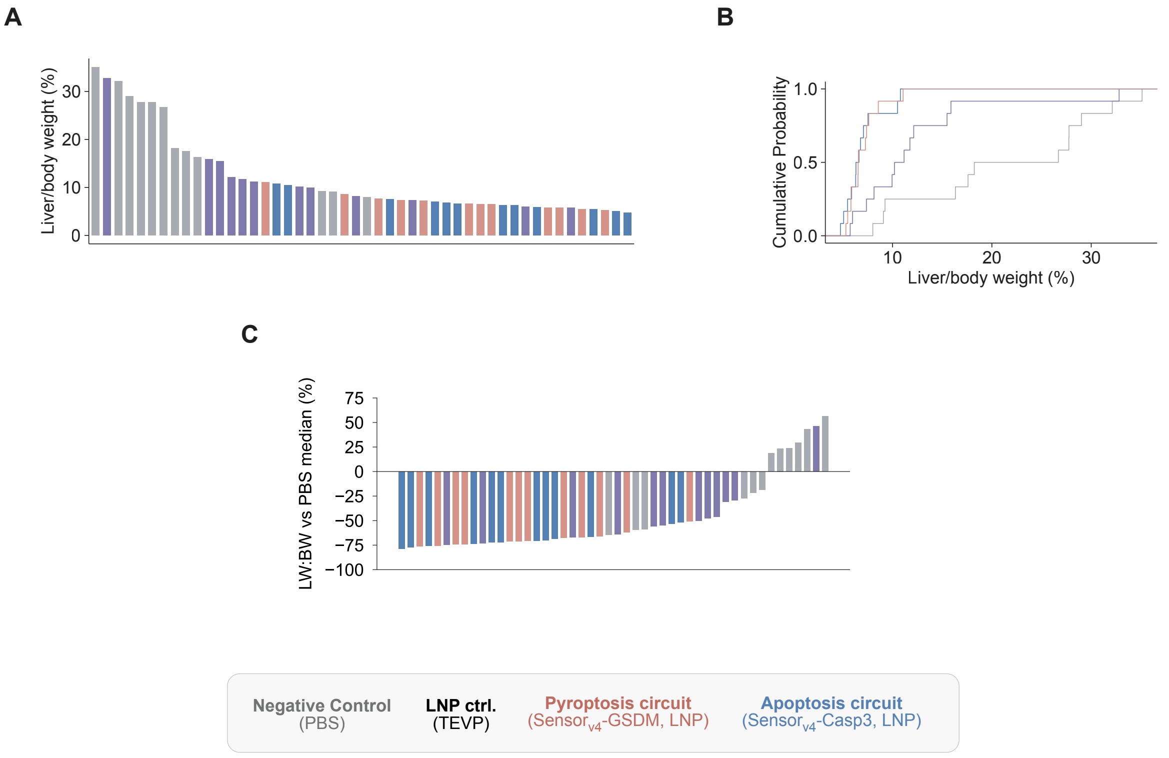


**Fig. S23. Quantification of liver tumor burden for fig. S22 (extended). (A)** Liver-to-body weight ratio (%) for individual animals, ranked in decreasing order. **(B)** Empirical cumulative distribution of liver-to-body weight ratios across treatment groups. **(C)** Percent change in liver-to-body weight ratio relative to the PBS control group median. Representative liver images for all animals are shown in fig. S22. Experimental details are shown in Fig. 4D.

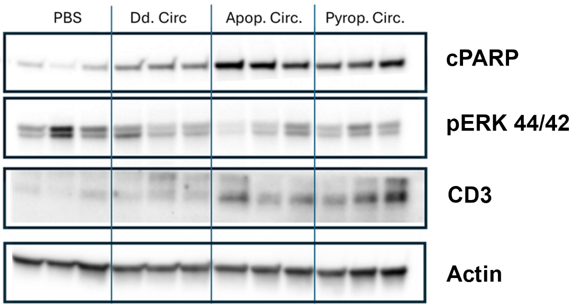


**Fig. S24. Circuit-treated liver tumors exhibit increased caspase activity and immune cell infiltration without suppression of ERK signaling.** Western blots of dissected $NRAS^{G12V}$ liver tumors from mice treated with PBS, a deactivated circuit control (Dd. Circ.), the $sensor_{v4}$-casp apoptosis circuit (Apop. Circ.), or the $sensor_{v4}$-GSDM pyroptosis circuit (Pyrop. Circ.). Western blots show cleaved PARP (cPARP), phospho-ERK 44/42 (pERK 44/42), the T cell marker CD3, and Actin (loading control). Each lane represents an individual mouse (n = 3 per condition).

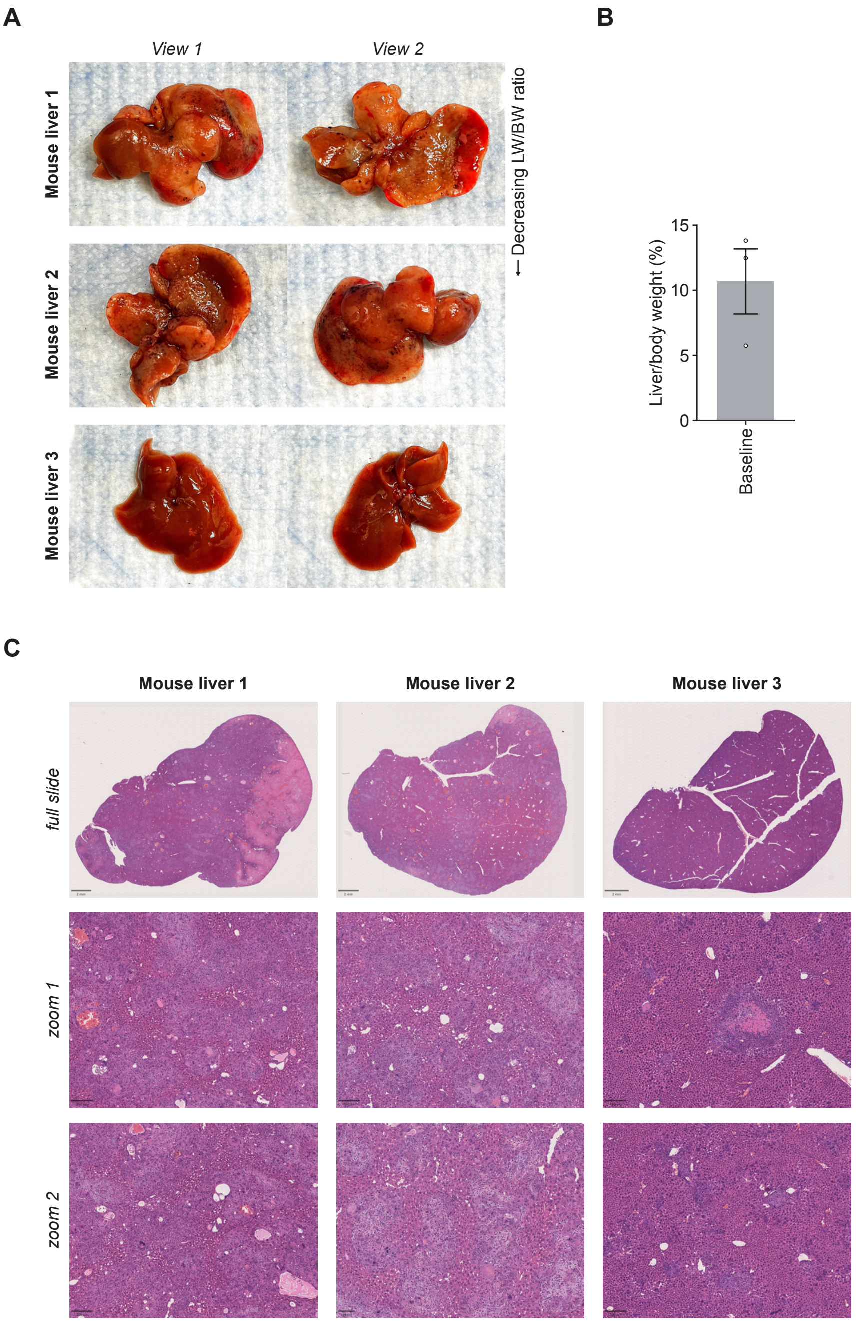


**Fig. S25. Day-10 baseline tumor burden in the KRAS$^{G12C}$ hydrodynamic tail-vein injection liver cancer model. (A)** Representative gross photographs of livers from three baseline mice harvested at day 10 post-tumor induction (HDT with KRAS$^{G12C}$, shTP53, and SB100 transposase), shown from two viewing angles and arranged by decreasing liver-to-body weight (LW/BW) ratio. **(B)** Liver-to-body weight ratio (%) for baseline mice at day 10 post-HDT. Error bars represent mean ± s.e. (n = 3 mice). **(C)** H&E-stained sections of the three baseline livers shown in (A).

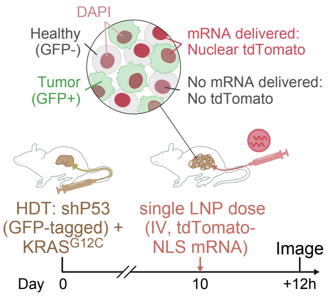


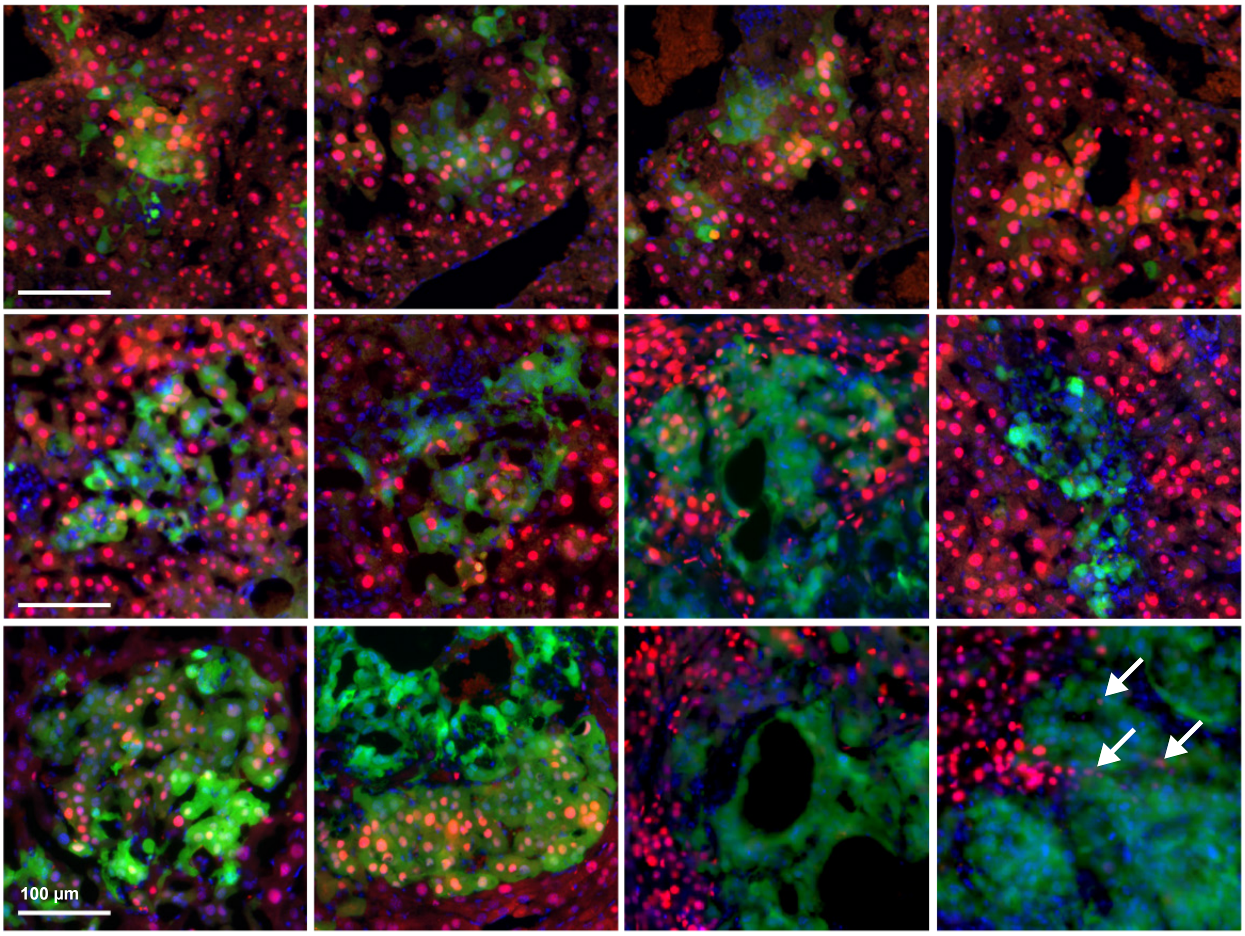


**Fig. S26. LNPs enable mRNA delivery to established KRAS$^{G12C}$ tumor nodules *in vivo*.** Top: experimental schematic. Liver tumors were initiated by hydrodynamic tail-vein injection (HDT) of GFP-tagged shP53 and KRAS$^{G12C}$ constructs at day 0. At day 10, when tumors were established, a single intravenous dose of LNPs encapsulating nuclear-localized tdTomato (tdTomato-NLS) mRNA was administered, and livers were collected 12 hours later for fluorescence imaging. Tumor cells are identified by GFP expression (green); successful mRNA delivery is indicated by nuclear tdTomato fluorescence (red); DAPI marks all nuclei (blue). Bottom: representative fluorescence micrographs of liver sections from tumor-bearing mice showing GFP-positive tumor nodules with co-localized nuclear tdTomato signal, confirming that intravenously administered LNPs access cells within established tumor nodules. White arrows highlight tumor cells (GFP-positive) with co-localized tdTomato signal, indicating successful mRNA delivery. Scale bar, 100 µm.

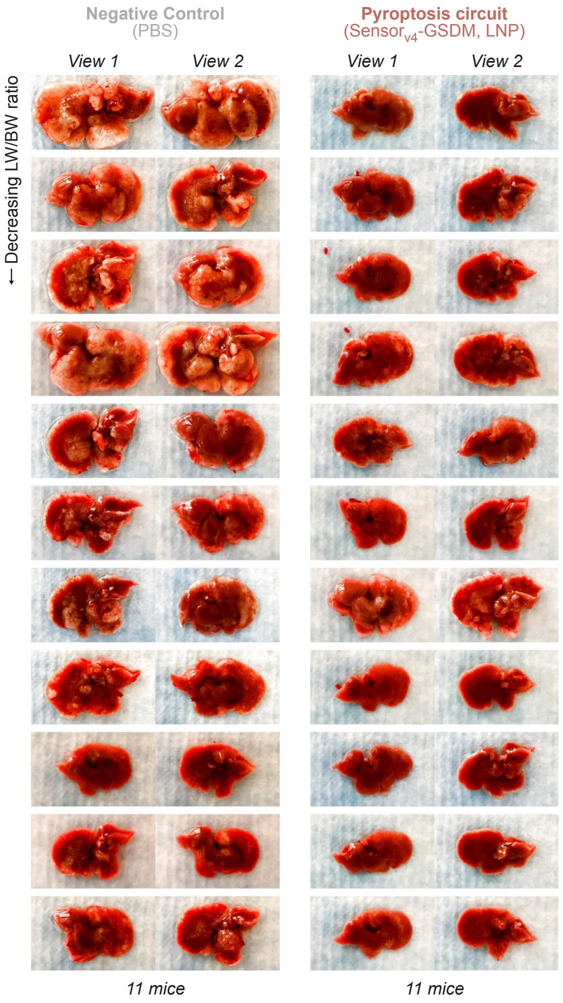


**Fig. S27. Individual liver images from KRAS$^{G12C}$ established-tumor treatment experiment (extended).** Representative photographs of all livers from PBS-treated negative control (left) and sensor$_{v4}$-GSDM pyroptosis circuit-treated (right) mice, shown from two views per animal. Livers are ordered by decreasing liver-to-body weight (LW/BW) ratio. Experimental details in Fig. 4F; full quantification in fig. S28.

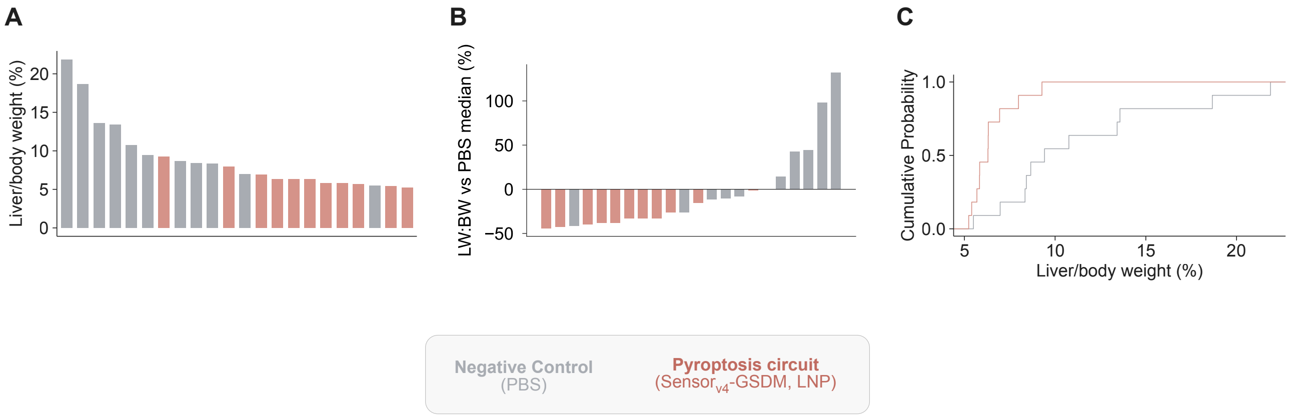


**Fig. S28. Quantification of liver tumor burden for fig. S27 (extended). (A)** Waterfall plot of liver-to-body weight ratio (%) for individual mice treated with PBS (gray) or the sensor$_{v4}$-GSDM pyroptosis circuit delivered as mRNA-LNP (green), sorted in descending order. **(B)** Percent change in liver-to-body weight ratio for each animal relative to the PBS group median. **(C)** Empirical cumulative distribution of liver-to-body weight ratios for PBS-treated and circuit-treated groups. Experimental details in Fig. 4F.

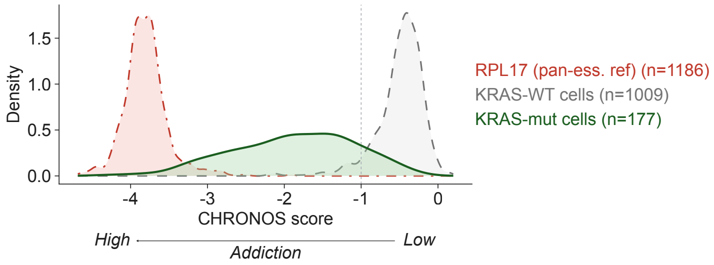


**Fig. S29. RAS-oncogene addiction is not universal across KRAS-mutant cancer cell lines.** Density distributions of CHRONOS CRISPR essentiality scores for KRAS across KRAS-mutant cell lines (green, n = 177) and KRAS-wild-type cell lines (gray, n = 1009) from the DepMap database. RPL17, a pan-essential gene, is shown as a reference (red, n = 1186). Lower CHRONOS scores indicate greater dependency on the gene for cell viability.

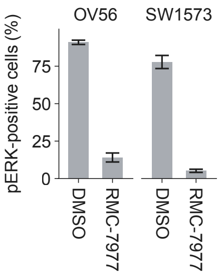


**Fig. S30. RMC-7977 potently suppresses pERK in non-addicted KRAS$^{G12C}$ cell lines.** Bar plots show the percentage of pERK-positive cells in OV56 (KRAS$^{G12C}$) and SW1573 (KRAS$^{G12C}$) cells treated with DMSO or RMC-7977, as measured by antibody staining and flow cytometry. Despite potent suppression of pERK signaling, these cell lines are minimally sensitive to RAS inhibitor-mediated killing (Fig. 5A), consistent with their low RAS oncogene addiction. Error bars represent mean ± s.d.

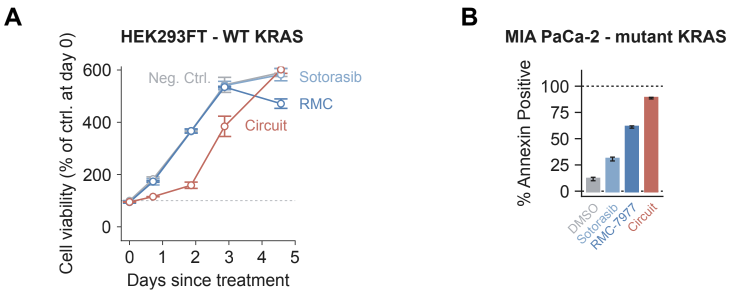


**Fig. S31. (A) Cell killing dynamics of RAS inhibitors and the therapeutic circuit in wild-type KRAS cells.** Cell viability of HEK293FT ($KRAS^{WT}$) cells treated with Sotorasib, RMC-7977, or the $sensor_{v4}$-casp circuit, measured over 5 days. Viability is expressed as a percentage of the matched control at day 0. Error bars represent mean ± s.d. **(B) Circuits induce apoptosis in RAS-mutant cancer cells.** Apoptotic cell death was confirmed by Annexin V staining and flow cytometry. Bars depict mean ± s.d.

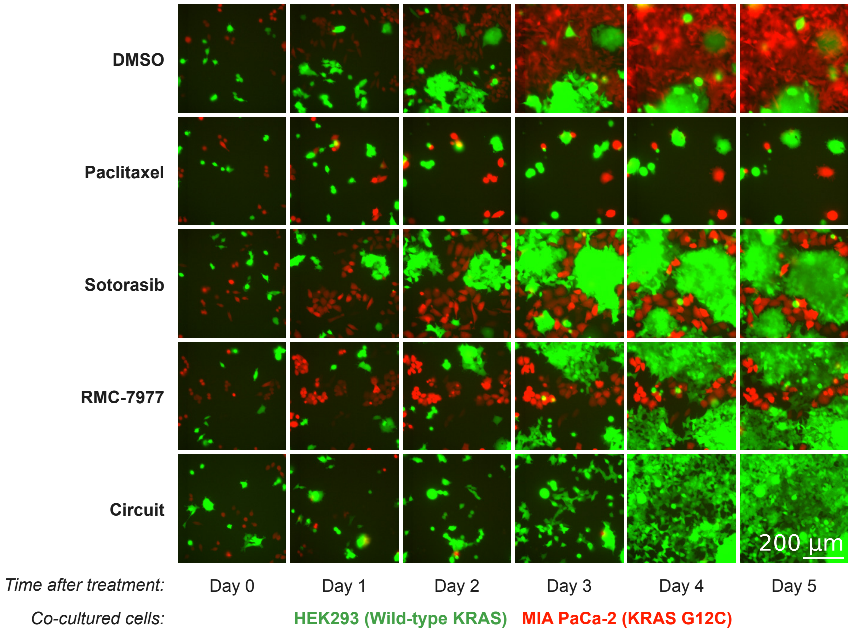


**Fig. S32. Extended time-lapse fluorescence imaging of co-cultured RAS wild-type and RAS-mutant cells treated with RAS inhibitors or the therapeutic circuit.** MIA PaCa-2 ($KRAS^{G12C}$) and HEK293 (wild-type KRAS) cells stably expressing mRuby3 (red) and GFP (green), respectively, were co-cultured and treated with DMSO (vehicle control), Paclitaxel (30 µM), Sotorasib (1 µM), RMC-7977 (1 µM), or the $sensor_{v4}$-casp apoptosis circuit (150 pg/µL engineered caspase-3 and 100 pg/µL $sensor_{v4}$ mRNA-LNP). Fluorescence images were acquired daily from day 0 through day 5 after treatment. While Sotorasib and RMC-7977 reduced the growth of mutant cells, the circuit actively eliminated MIA PaCa-2 cells within the first few days while sparing HEK293 cells. Scale bar, 200 µm.

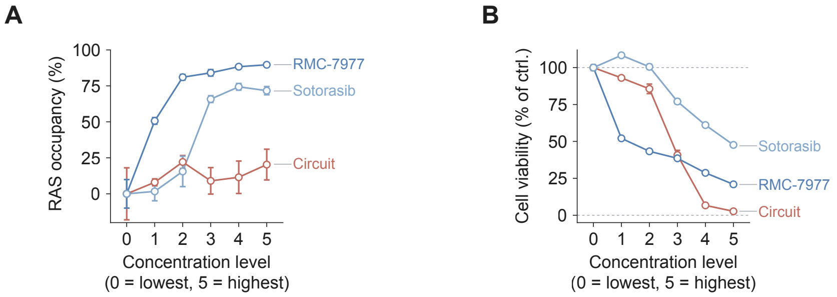


**Fig. S33. RAS occupancy and cell viability measurements across matched concentration levels for RAS inhibitors and the sensor$_{v4}$-casp circuit.** MIA PaCa-2 (KRAS$^{G12C}$) cells were treated with increasing concentrations of Sotorasib, RMC-7977, or the sensor$_{v4}$-casp circuit, and RAS occupancy was measured using a split luciferase RAS-RBD displacement assay. **(A)** RAS occupancy (%) as a function of concentration level (0 = lowest, 5 = highest) for RMC-7977, Sotorasib, and the circuit. Both Sotorasib and RMC-7977 achieved high RAS occupancy at elevated concentrations, whereas the circuit occupied a small fraction of RAS across all concentration levels. **(B)** Cell viability (% of control) as a function of concentration level. The circuit achieved near-complete cell killing at concentration levels where RAS occupancy remained low, whereas Sotorasib and RMC-7977 required higher concentrations to reduce viability. Error bars represent mean ± s.d.

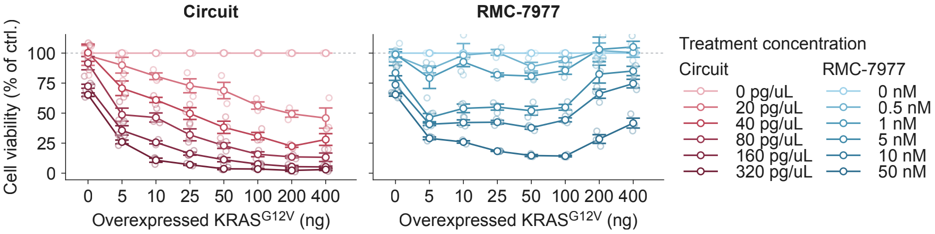


**Fig. S34. Therapeutic circuits are robust to RAS amplification. (A)** Cell viability of MCF10A cells following EGF withdrawal as a function of overexpressed KRAS$^{G12V}$ levels and treatment concentration for the sensor$_{v4}$-casp circuit (left; 0 to 320 pg/µL) or RMC-7977 (right; 0 to 50 nM). Increasing KRAS$^{G12V}$ expression reduces circuit-treated cell viability, whereas RMC-7977 efficacy declines at elevated KRAS$^{G12V}$ levels, recapitulating amplification-driven resistance. Error bars represent mean ± s.d.

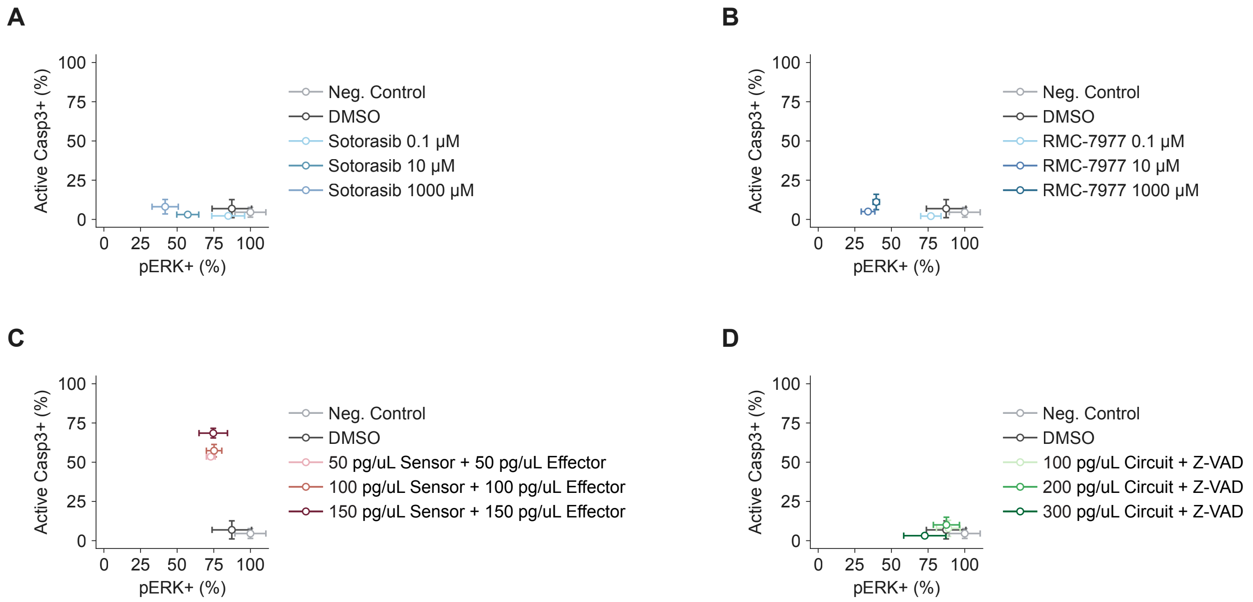


**Fig. S35. RAS inhibitors and the sensor$_{v4}$-casp circuit have distinct effects on RAS signaling and caspase-3 activation in MIA PaCa-2 cells.** pERK and active caspase-3 were quantified by intracellular antibody staining and flow cytometry. Error bars represent mean ± s.d.

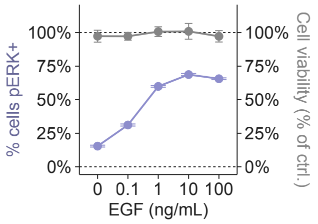


**Fig. S36. The sensor$_{v4}$-casp circuit does not reduce the viability of wild-type RAS cells upon EGF-mediated RAS pathway stimulation.** HEK293 (KRAS$^{WT}$) cells were treated with the sensor$_{v4}$-casp circuit and stimulated with increasing concentrations of EGF (0 to 100 ng/mL). Left y-axis (purple): percentage of pERK-positive cells, quantified by antibody staining and flow cytometry. Right y-axis (gray): cell viability measured by CellTiter-Glo, expressed as percentage of untreated control. EGF stimulation increased the fraction of pERK-positive cells from ~15% to ~70%, yet circuit-treated cell viability remained near 100% across all EGF concentrations. Error bars for pERK measurements represent 95% confidence intervals derived from bootstrapping (1,000 iterations). Error bars for cell viability represent mean ± s.d. (n = 2 replicates).

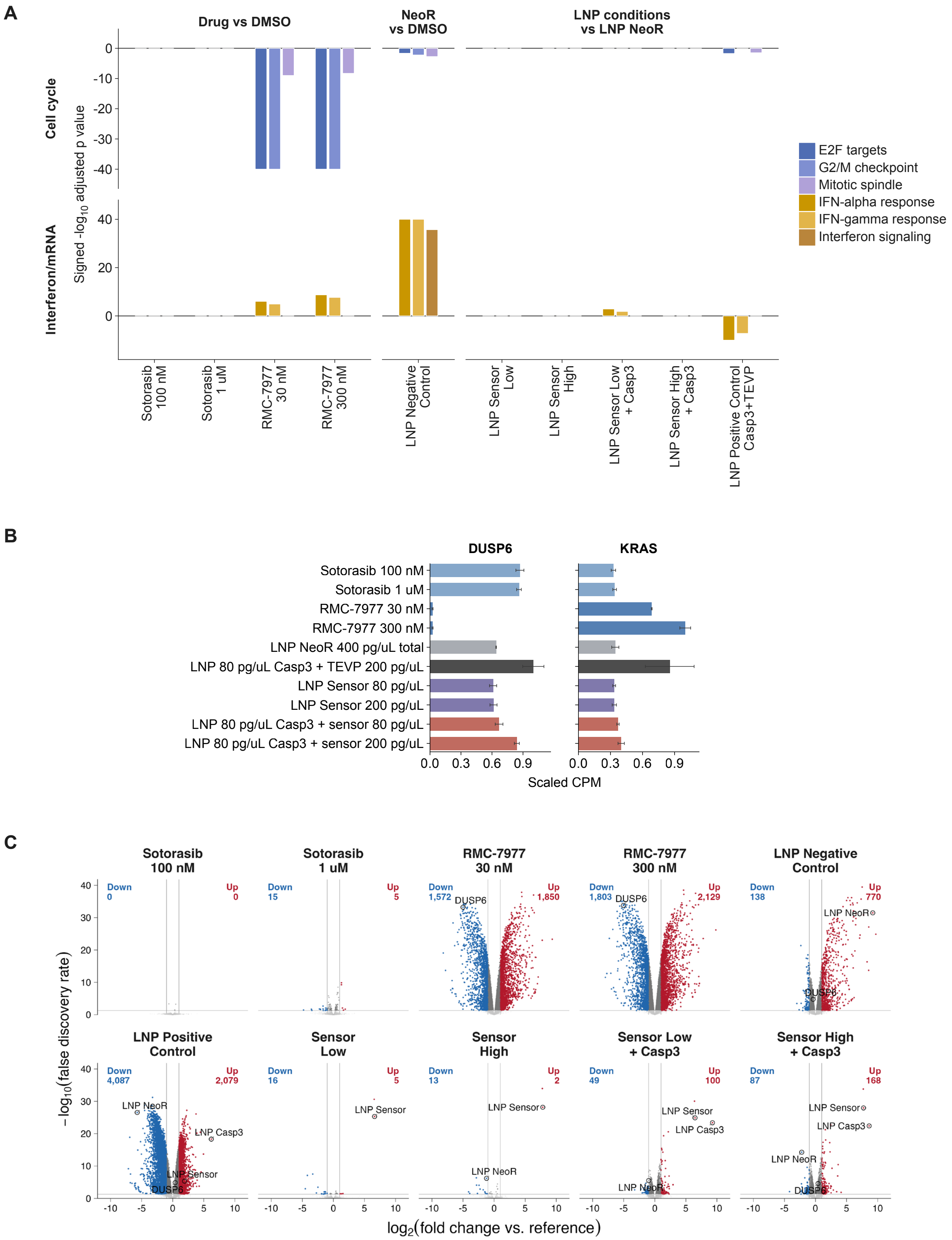


**Fig. S37. Extended transcriptomic analysis of RAS inhibitors and the sensor$_{v4}$-casp circuit in MCF10A (KRAS$^{WT}$) cells. (A)** Pathway-level enrichment of cell cycle (left) and interferon/mRNA-sensing (right) Hallmark gene sets across treatment conditions. Bars depict the signed -log$_{10}$(adjusted *p* value) for E2F targets, G2/M checkpoint, and mitotic spindle (cell cycle), and IFN-alpha response, IFN-gamma response, and interferon signaling (interferon/mRNA). Drug conditions (Sotorasib at 100 nM and 1 µM; RMC-7977 at 30 nM and 300 nM) are compared to DMSO; LNP negative control (NeoR) is compared to DMSO; remaining LNP conditions (sensor at low and high concentrations, sensor + engineered caspase-3 at low and high concentrations, and positive control Casp3 + TEVP) are compared to LNP NeoR. **(B)** Change of DUSP6 (left) and KRAS (right) transcript levels across conditions. LNP conditions are labeled with mRNA concentrations (pg/µL). Error bars represent mean ± s.d. **(C)** Volcano plots of differentially expressed genes for each condition. Numbers of significantly downregulated (blue) and upregulated (red) genes are indicated per panel. Key transcripts (DUSP6, KRAS, and LNP cargo mRNAs) are labeled. RNA-seq was performed 24 hours after treatment.

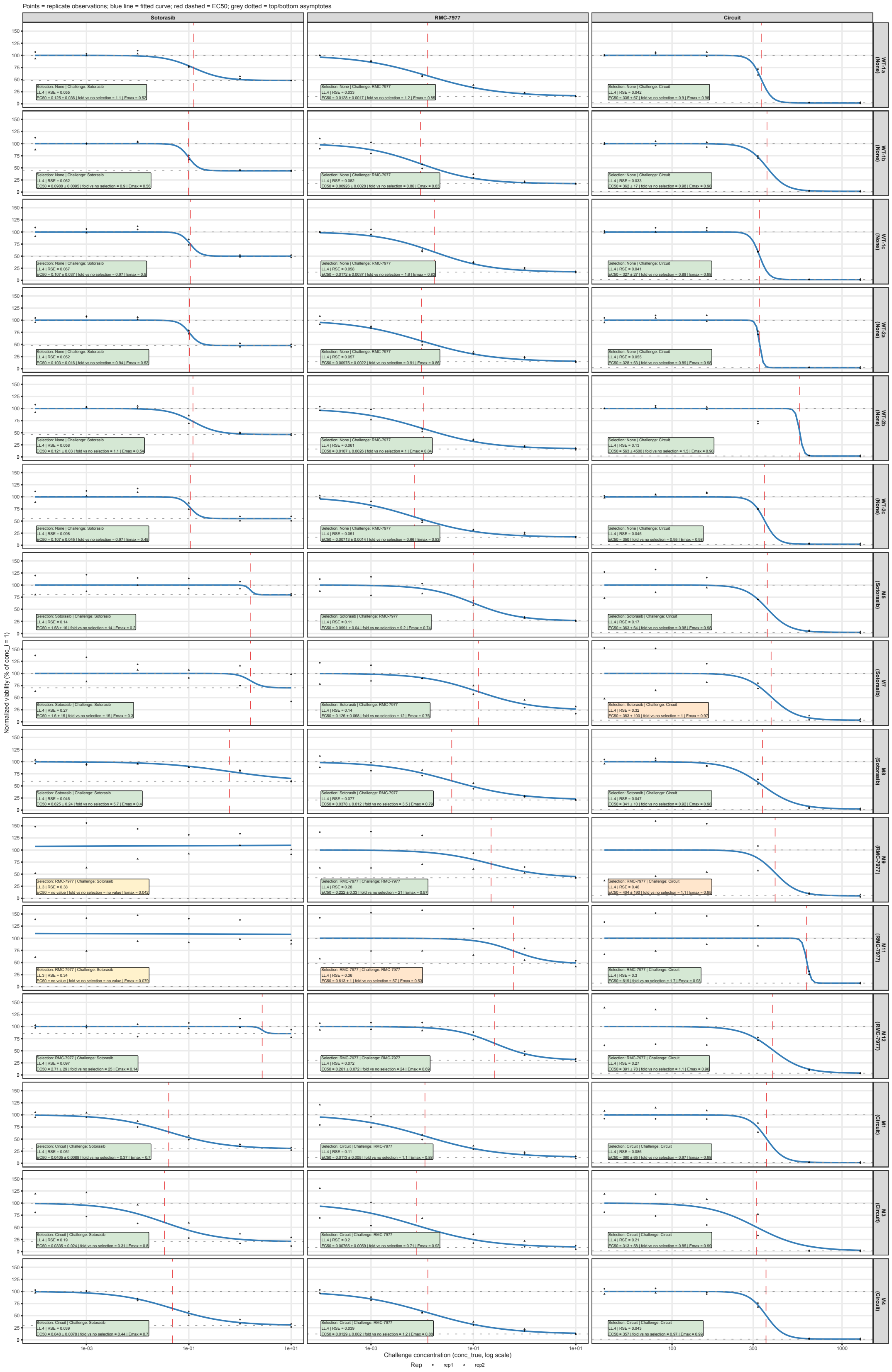


**Fig. S38. Individual dose-response curves for the acquired resistance experiment I.** MIA PaCa-2 ($KRAS^{G12C}$) cells were subjected to 2 months of continuous selection with Sotorasib, RMC-7977, or the $sensor_{v4}$-casp circuit at each treatment's $EC_{70}$, alongside non-selected controls (three independent selection replicates per condition). Each selected and non-selected culture was then challenged with Sotorasib (left column), RMC-7977 (center column), or the $sensor_{v4}$-casp circuit (right column) across a dose range. Each panel shows one selection replicate challenged with one treatment. Points represent replicate observations (n = 2 technical replicates per concentration, distinguished by shape). Blue lines are fitted four-parameter log-logistic curves; red dashed vertical lines indicate the fitted $EC_{50}$; grey dotted horizontal lines indicate the fitted top and bottom asymptotes. Fitted model parameters ($EC_{50}$, slope, and residual viability) are annotated within each panel. These individual curves underlie the summary quantification presented in Fig. 6B.

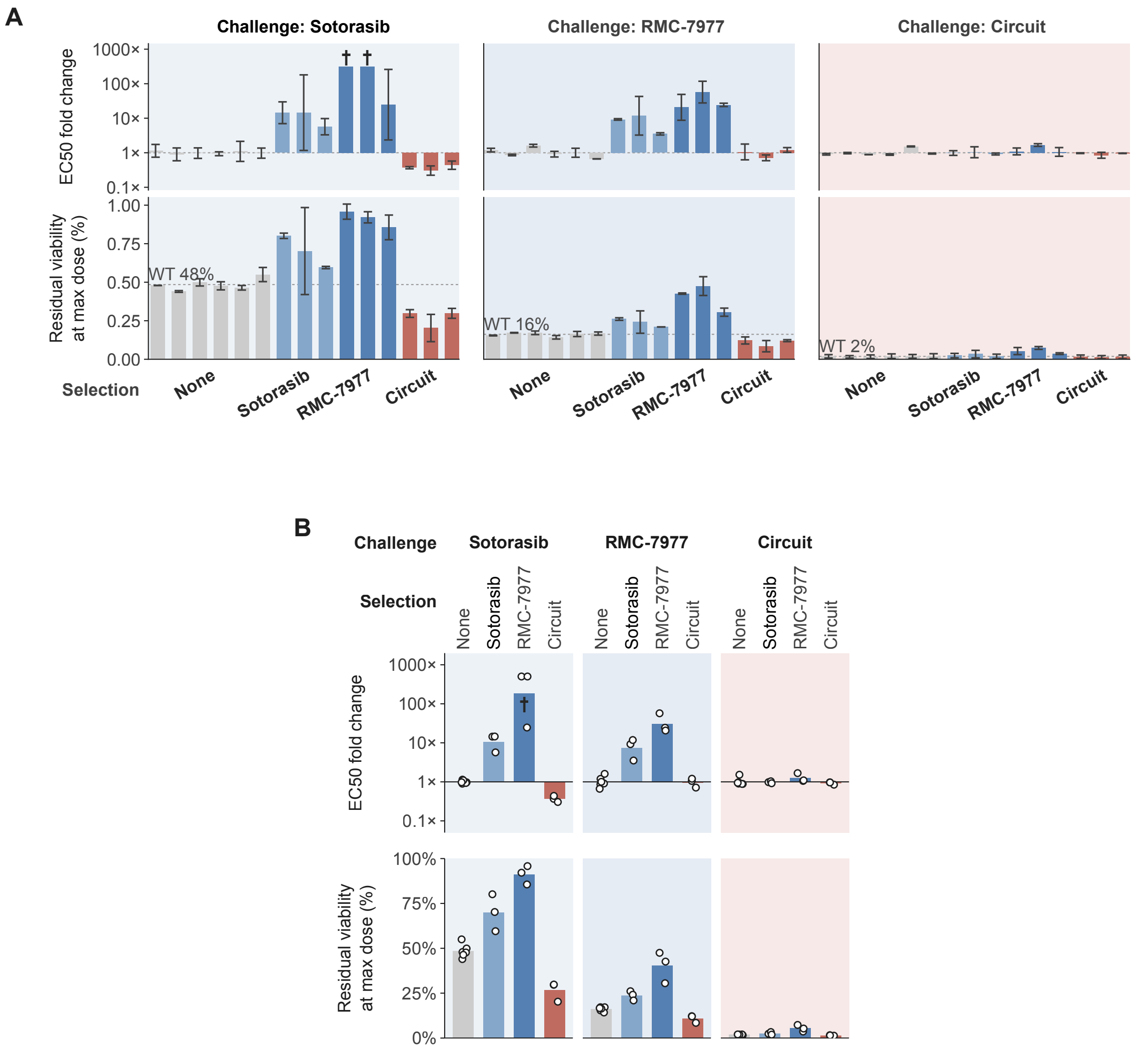


**Fig. S39. Full cross-resistance matrix for acquired resistance experiment I (fig. S38). (A)** $EC_{50}$ fold change (top row) and residual viability at maximum dose (bottom row) for each selection-challenge combination, organized by challenge treatment (columns). Dashed lines indicate baseline $EC_{50}$ (1x) and wild-type residual viability (WT 48% for Sotorasib, WT 16% for RMC-7977, WT 2% for circuit). ***Daggers denote conditions where $EC_{50}$ was not reached within the tested concentration range.*** **(B)** Same data as (A), reorganized by challenge and selection condition, with individual replicate values shown as open circles. Error bars represent mean ± s.d., n = 3 independent selection replicates.

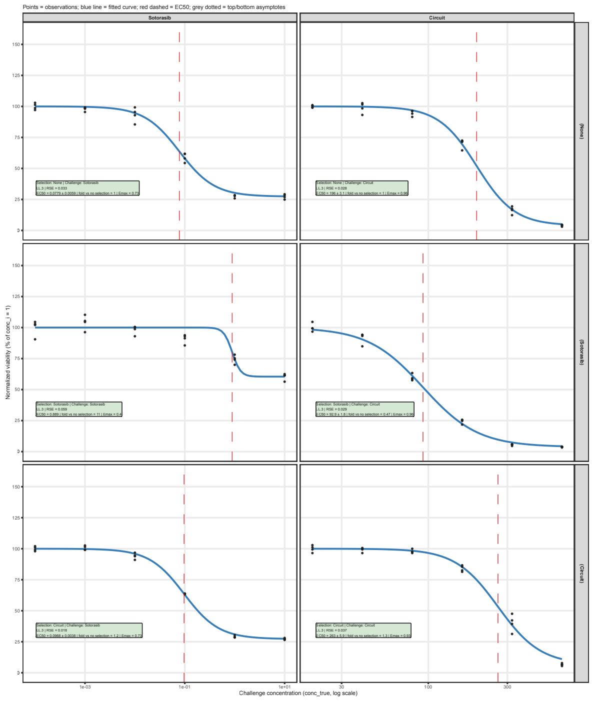


**Fig. S40. Individual dose-response curves for the acquired resistance experiment II**. MIA PaCa-2 cells were repeatedly exposed to Sotorasib or sensor$_{v4}$-casp circuit (new treatment every 5 days for 3 weeks), and then challenged with either Sotorasib or circuit. Each panel shows one selection replicate challenged with one treatment. Points represent replicate observations (n = 4 technical replicates per concentration). Blue lines are fitted four-parameter log-logistic curves; red dashed vertical lines indicate the fitted $EC_{50}$; grey dotted horizontal lines indicate the fitted top and bottom asymptotes. Fitted model parameters ($EC_{50}$, slope, and residual viability) are annotated within each panel.

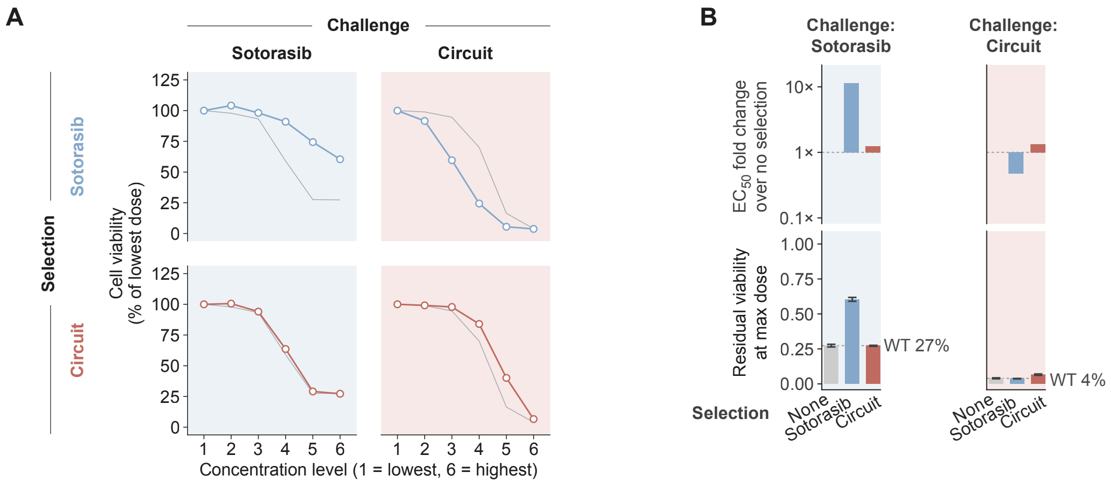


**Fig. S41. Full cross-resistance matrix for acquired resistance experiment II (fig. S40). (A)** Dose-response curves after continuous selection with Sotorasib (top row) or the sensor$_{v4}$-casp circuit (bottom row), challenged with Sotorasib (left column) or the circuit (right column). Gray curves represent non-selected negative controls. Cell viability was measured by CellTiter-Glo and normalized to the lowest concentration level (level 1 = lowest, level 6 = highest). **(B)** Quantification of resistance. Top: $EC_{50}$ fold change over no-selection controls (log scale). Bottom: residual viability at maximum dose. Bars are grouped by selection condition (None, Sotorasib, Circuit) and separated by challenge (Sotorasib, left; Circuit, right). Dashed lines indicate no change (1x) or the baseline residual viability of non-selected cells, respectively. Annotated percentages indicate wild-type (non-selected) residual viability at maximum dose (WT 27% for Sotorasib challenge; WT 4% for circuit challenge). Error bars represent mean ± s.d. across technical replicates.

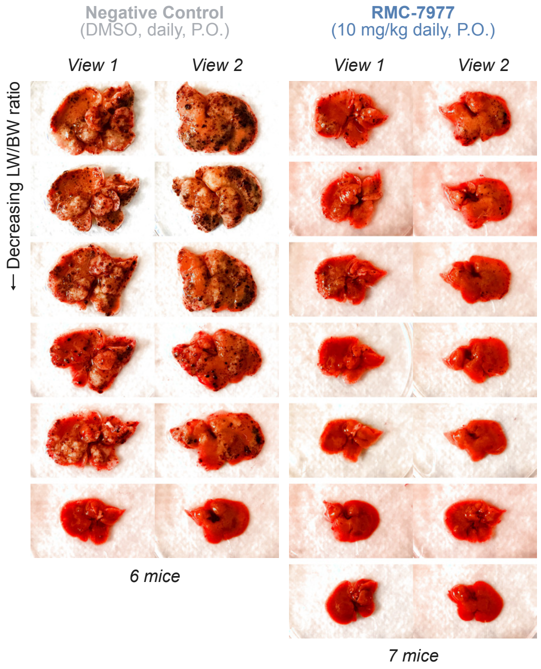


**Fig. S42. RMC-7977 suppresses tumor burden in the NRAS$^{G12V}$/shTP53 HDT liver tumor model.** Representative photographs of dissected livers from mice bearing NRAS$^{G12V}$/shTP53-driven liver tumors treated with daily oral vehicle (DMSO; n = 6 mice) or RMC-7977 (10 mg/kg daily, P.O.; n = 7 mice) over 28 days (treatment start 5 days post tumor initiation). Two views are shown for each liver. Livers are ordered by decreasing liver-to-body weight (LW/BW) ratio within each group. Quantification of LW/BW ratios is shown in Fig. S43.

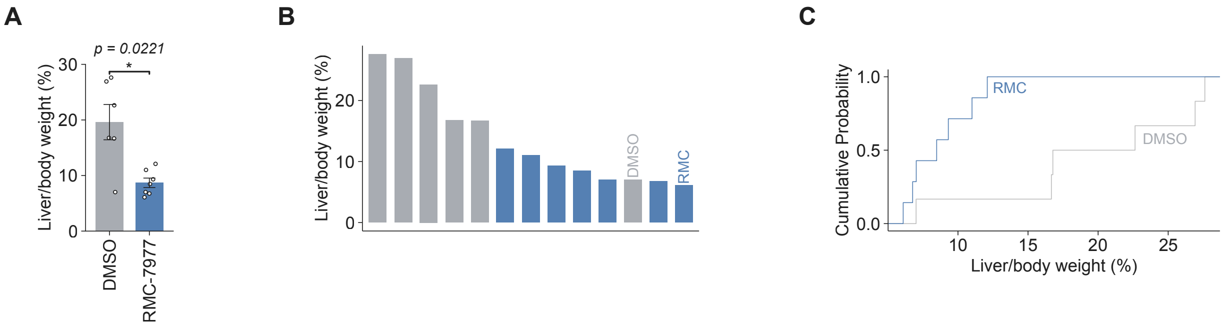


**Fig. S43. Quantification of liver tumor burden for fig. S42. (A)** Liver-to-body weight ratio (%) for mice treated with DMSO or RMC-7977. Bars depict mean ± s.e.; data points denote liver-to-body weight ratio of individual animals. **(B)** Liver-to-body weight ratio (%) for individual animals, ranked in descending order. **(C)** Empirical cumulative distribution of liver-to-body weight ratio (%) for DMSO-treated and RMC-7977-treated animals.

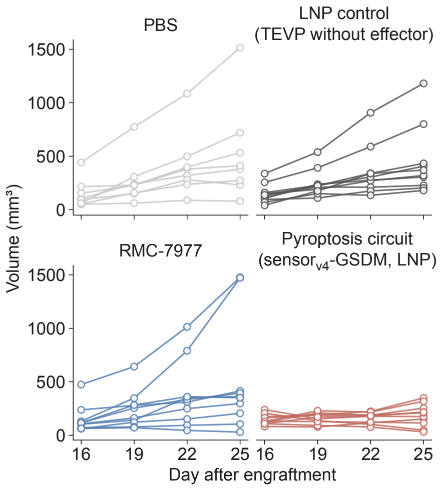


**Fig. S44. Individual tumor growth trajectories for the SW1573 cell-line-derived xenograft (CDX) model.** Individual tumor volumes ($mm^3$) over time (day after engraftment) for each mouse in the four treatment groups from the experiment quantified in Fig. 6E.

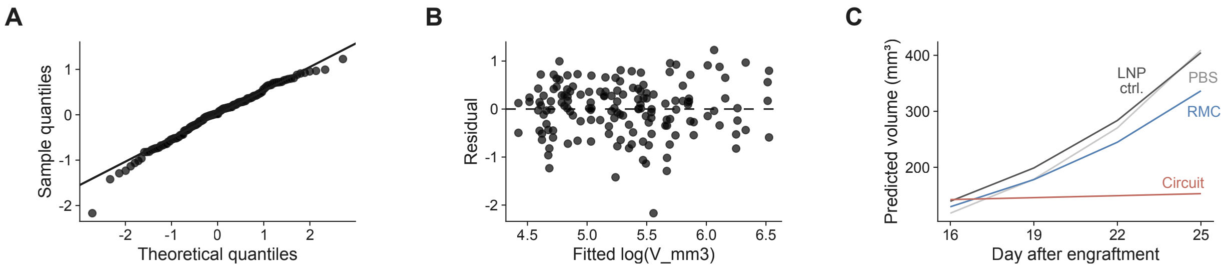


**Fig. S45. Diagnostic plots for the linear mixed-effects model used to analyze SW1573 CDX tumor growth.** Tumor volumes from the experiment shown in Fig. 6E were log-transformed and fit with a linear mixed-effects model (Methods). **(A)** Quantile-quantile plot of model residuals against theoretical normal quantiles, assessing the normality assumption. **(B)** Residuals plotted against fitted log-transformed tumor volume, assessing homoscedasticity. The dashed line indicates zero residual. **(C)** Model-predicted tumor volume ($mm^3$) over time (days after engraftment) for each treatment group.

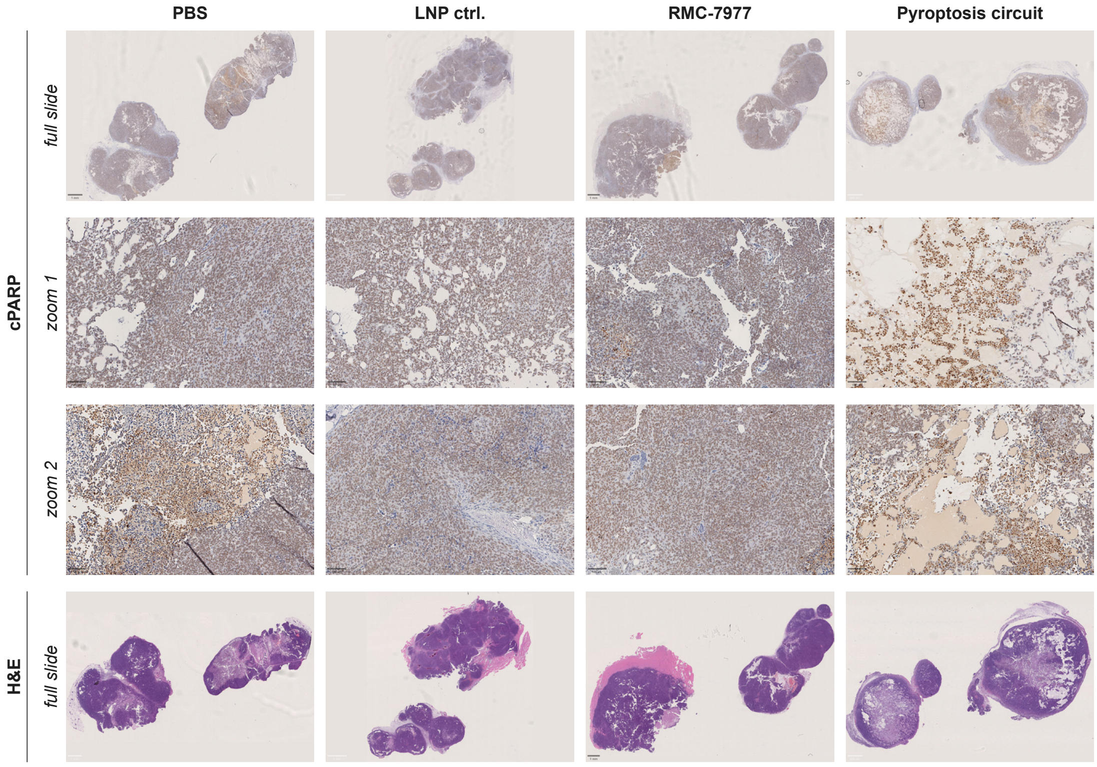


**Fig. S46. Cleaved PARP immunohistochemistry and histology of SW1573 xenograft tumors treated with the sensor$_{v4}$-GSDM pyroptosis circuit or controls.** Representative cleaved PARP (cPARP) immunohistochemistry (top three rows) and hematoxylin and eosin (H&E) staining (bottom row) of SW1573 tumors from Fig. 6E.

### Key Resource Table

| Reagent / Resource | Source | Identifier |
|---|---|---|
| Bacterial strains | | |
| Stable Competent E. coli (High Efficiency) | NEB | C3040H |
| Reagents | | |
| Dulbecco's Modified Eagle Medium | ThermoFisher Scientific | 11960-069 |
| Fetal bovine serum | Avantor | 97068-085 |
| Penicillin-Streptomycin-Glutamine | ThermoFisher Scientific | 10378016 |
| Sodium pyruvate | ThermoFisher Scientific | 11360070 |
| Minimal Essential Medium Non-Essential Amino Acids | ThermoFisher Scientific | 11140050 |
| Trypsin-EDTA (0.25%) | ThermoFisher Scientific | 25200056 |
| Dulbecco's Phosphate Buffered Saline | ThermoFisher Scientific | 14040117 |
| Opti-MEM I Reduced Serum Medium | ThermoFisher Scientific | 31985070 |
| N1-Methylpseudouridine-5'-Triphosphate | TriLink | N-1081 |
| Deoxynucleotide (dNTP) Solution Mix | NEB | N0447L |
| CleanCap AG | TriLink | N-7113 |
| DNase I (RNase-free) | NEB | M0303L |
| Lipofectamine 3000 | ThermoFisher Scientific | L3000008 |
| LB Broth with agar (Lennox) | Sigma Aldrich | L2897-250G |
| Human EGF, Animal-Free Recombinant Protein, PeproTech | Gibco | AF-100-15-500UG |
| Sotorasib | MedChemExpress | HY-114277 |
| RMC-7977 | MedChemExpress | HY-156498 |
| Paclitaxel | MedChemExpress | HY-B0015 |
| Deposited data (The following data will be deposited prior to publication) | | |
| Raw sequencing reads for RNA-seq of drugs and circuit treatment MCF10A cells. | This study | |
| Raw microscopy images | This study | |
| Cell lines | | |
| HEK293 | ATCC | CRL-1573 |
| HEK293 mRuby3 | This study | |
| MIA PaCa-2 | ATCC | CRL-1420 |
| MIA PaCa-2 GFP | Fenics Bio | CL-1199 |
| NCI-H358 | ATCC | CRL-5807 |
| NCI-H441 | ATCC | HTB-174 |
| HepG2 | ATCC | HB-8065 |
| PLC/PRF/5 | ATCC | CRL-8024 |
| SNU-387 | ATCC | CRL-2237 |
| SNU-423 | ATCC | CRL-2238 |
| SNU-449 | ATCC | CRL-2234 |
| SNU-398 | ATCC | CRL-2233 |
| SNU-475 | ATCC | CRL-2236 |
| SW1573 | ATCC | CRL-2170 |
| Panc1 | ATCC | CRL-1469 |
| Recombinant DNA | | |
| Plasmid for Ectopic Overexpression of HRAS G12A | This study | pLM0155 |
| Plasmid for Ectopic Overexpression of HRAS G12C | This study | pLM0156 |
| Plasmid for Ectopic Overexpression of HRAS G12D | This study | pLM0157 |
| Plasmid for Ectopic Overexpression of HRAS G12R | This study | pLM0158 |
| Plasmid for Ectopic Overexpression of HRAS G12S | This study | pLM0159 |
| Plasmid for Ectopic Overexpression of HRAS G12V | This study | pLM0160 |
| Plasmid for Ectopic Overexpression of HRAS G13C | This study | pLM0161 |
| Plasmid for Ectopic Overexpression of HRAS G13D | This study | pLM0162 |
| Plasmid for Ectopic Overexpression of HRAS G13dup | This study | pLM0163 |
| Plasmid for Ectopic Overexpression of HRAS G13R | This study | pLM0164 |
| Plasmid for Ectopic Overexpression of HRAS G13S | This study | pLM0165 |
| Plasmid for Ectopic Overexpression of HRAS G13V | This study | pLM0166 |
| Plasmid for Ectopic Overexpression of HRAS Q61H | This study | pLM0167 |
| Plasmid for Ectopic Overexpression of HRAS Q61K | This study | pLM0168 |
| Plasmid for Ectopic Overexpression of HRAS Q61L | This study | pLM0169 |
| Plasmid for Ectopic Overexpression of HRAS Q61R | This study | pLM0170 |
| Plasmid for Ectopic Overexpression of KRAS-4B A146P | This study | pLM0171 |

| | | |
|---|---|---|
| Plasmid for Ectopic Overexpression of KRAS-4B A146T | This study | pLM0172 |
| Plasmid for Ectopic Overexpression of KRAS-4B A146V | This study | pLM0173 |
| Plasmid for Ectopic Overexpression of KRAS-4B G12A | This study | pLM0174 |
| Plasmid for Ectopic Overexpression of KRAS-4B G12C | This study | pLM0175 |
| Plasmid for Ectopic Overexpression of KRAS-4B G12D | This study | pLM0176 |
| Plasmid for Ectopic Overexpression of KRAS-4B G12R | This study | pLM0177 |
| Plasmid for Ectopic Overexpression of KRAS-4B G12S | This study | pLM0178 |
| Plasmid for Ectopic Overexpression of KRAS-4B G12V | This study | pLM0179 |
| Plasmid for Ectopic Overexpression of KRAS-4B G13C | This study | pLM0180 |
| Plasmid for Ectopic Overexpression of KRAS-4B G13D | This study | pLM0181 |
| Plasmid for Ectopic Overexpression of KRAS-4B G13V | This study | pLM0182 |
| Plasmid for Ectopic Overexpression of KRAS-4B Q61E | This study | pLM0183 |
| Plasmid for Ectopic Overexpression of KRAS-4B Q61H | This study | pLM0184 |
| Plasmid for Ectopic Overexpression of KRAS-4B Q61K | This study | pLM0185 |
| Plasmid for Ectopic Overexpression of KRAS-4B Q61L | This study | pLM0186 |
| Plasmid for Ectopic Overexpression of KRAS-4B Q61P | This study | pLM0187 |
| Plasmid for Ectopic Overexpression of KRAS-4B Q61R | This study | pLM0188 |
| Plasmid for Ectopic Overexpression of NRAS G12A | This study | pLM0189 |
| Plasmid for Ectopic Overexpression of NRAS G12C | This study | pLM0190 |
| Plasmid for Ectopic Overexpression of NRAS G12D | This study | pLM0191 |
| Plasmid for Ectopic Overexpression of NRAS G12R | This study | pLM0192 |
| Plasmid for Ectopic Overexpression of NRAS G12S | This study | pLM0193 |
| Plasmid for Ectopic Overexpression of NRAS G12V | This study | pLM0194 |
| Plasmid for Ectopic Overexpression of NRAS G13D | This study | pLM0195 |
| Plasmid for Ectopic Overexpression of NRAS G13R | This study | pLM0196 |
| Plasmid for Ectopic Overexpression of NRAS Q61* | This study | pLM0197 |
| Plasmid for Ectopic Overexpression of NRAS Q61H | This study | pLM0198 |
| Plasmid for Ectopic Overexpression of NRAS Q61K | This study | pLM0199 |
| Plasmid for Ectopic Overexpression of NRAS Q61L | This study | pLM0200 |
| Plasmid for Ectopic Overexpression of NRAS Q61P | This study | pLM0201 |
| Plasmid for Ectopic Overexpression of NRAS Q61R | This study | pLM0202 |
| Plasmid for Ectopic Overexpression of HRAS WT | This study | pAL0109 |
| Plasmid for Ectopic Overexpression of KRAS-4B WT | This study | pAL0106 |
| Plasmid for Ectopic Overexpression of NRAS WT | This study | pAL0111 |
| Plasmid for iTEVP-CAAX reporter | This study | pAL0031 |
| Plasmid for mRuby3 | This study | pZAG0744 |
| Plasmid for neomycin resistance gene | This study | pZAG0530 |
| Plasmid to express 12VC1-nTEVP-tevs-dcTEVP-IRES-BFP | This study | P-0215 |
| Plasmid to express 12VC1-nTEVP-tevD-dcTEVP-IRES-BFP | This study | P-0216 |
| Plasmid to express 12VC1-nTEVP-tevs-DHFR-IRES-BFP | This study | P-0218 |
| Plasmid to express 12VC1-nTEVP-tevs-dcTEVP-3xNLS-IRES-BFP | This study | P-0219 |
| Plasmid to express 12VC1-nTEVP-tevD-dcTEVP-3xNLS-IRES-BFP | This study | P-0220 |
| Plasmid to express 12VC1-cTEVP-tevs-dnTEVP | This study | P-0222 |
| Plasmid to express 12VC1-cTEVP-tevD-dnTEVP | This study | P-0223 |
| Plasmid to express 12VC1-cTEVP-tevs-DHFR | This study | P-0225 |
| Plasmid to express 12VC1-nTEVP-tevs-3xNLS-IRES-BFP | This study | P-0393 |
| Plasmid to express P4-nTEVP-IRES-BFP | This study | pLM104 |
| Plasmid to express P3-cTEVP | This study | pLM107 |
| Plasmid to express GSGSG-nTEVP-IRES-BFP | This study | PTC-01 |
| Plasmid to express CRAF-nTEVP-IRES-BFP | This study | PTC-02 |
| Plasmid to express HRAS-peptide-CRAF-nTEVP-IRES-BFP | This study | PTC-03 |
| Plasmid to express NS1-nTEVP-IRES-BFP | This study | PTC-04 |
| Plasmid to express JAM20-nTEVP-IRES-BFP | This study | PTC-05 |
| Plasmid to express K13-nTEVP-IRES-BFP | This study | PTC-06 |
| Plasmid to express K19-nTEVP-IRES-BFP | This study | PTC-07 |
| Plasmid to express RA_RASSF1-nTEVP-IRES-BFP | This study | PTC-08 |
| Plasmid to express RA_RASSF6-nTEVP-IRES-BFP | This study | PTC-09 |
| Plasmid to express R15m10-nTEVP-IRES-BFP | This study | PTC-10 |
| Plasmid to express 12VC1-nTEVP-IRES-BFP | This study | PTC-11 |
| Plasmid to express 12VC3-nTEVP-IRES-BFP | This study | PTC-12 |
| Plasmid to express GSGSG-cTEVP-WT | This study | PTC-13 |
| Plasmid to express CRAF-cTEVP-WT | This study | PTC-14 |
| Plasmid to express HRAS-peptide-CRAF-cTEVP-WT | This study | PTC-15 |

| | | |
|---|---|---|
| Plasmid to express NS1-cTEVP-WT | This study | PTC-16 |
| Plasmid to express JAM20-cTEVP-WT | This study | PTC-17 |
| Plasmid to express K13-cTEVP-WT | This study | PTC-18 |
| Plasmid to express K19-cTEVP-WT | This study | PTC-19 |
| Plasmid to express RA_RASSF1-cTEVP-WT | This study | PTC-20 |
| Plasmid to express RA_RASSF6-cTEVP-WT | This study | PTC-21 |
| Plasmid to express R15m10-cTEVP-WT | This study | PTC-22 |
| Plasmid to express 12VC1-cTEVP-WT | This study | PTC-23 |
| Plasmid to express 12VC3-cTEVP-WT | This study | PTC-24 |
| Plasmid to express GSGSG-nTEVP-WT-IRES-BFP | This study | PTC-25 |
| Plasmid to express GSGSG-nTEVP-T30A-IRES-BFP | This study | PTC-26 |
| Plasmid to express GSGSG-nHyperTEV60-IRES-BFP | This study | PTC-27 |
| Plasmid to express GSGSG-cTEVP-WT | This study | PTC-28 |
| Plasmid to express GSGSG-cTEVP$\Delta$ | This study | PTC-29 |
| Plasmid to express GSGSG-cTEVP$\Delta$-I138T-S153N-T180A (uTEV3) | This study | PTC-30 |
| Plasmid to express GSGSG-cHyperTEV60 | This study | PTC-31 |
| Plasmid to express CRAF-RBD-nTEVP-WT-IRES-BFP | This study | PTC-32 |
| Plasmid to express CRAF-RBD-nTEVP-T30A-IRES-BFP | This study | PTC-33 |
| Plasmid to express CRAF-RBD-nHyperTEV60-IRES-BFP | This study | PTC-34 |
| Plasmid to express CRAF-RBD-cTEVP-WT | This study | PTC-35 |
| Plasmid to express CRAF-RBD-cTEVP$\Delta$ | This study | PTC-36 |
| Plasmid to express CRAF-RBD-cTEVP$\Delta$-I138T-S153N-T180A (uTEV3) | This study | PTC-37 |
| Plasmid to express CRAF-RBD-cHyperTEV60 | This study | PTC-38 |
| Plasmid to express GRB2-IRES-mRuby3 | This study | PTC-42 |
| Plasmid to express 12VC1-nTEVP-T30A-IRES-BFP | This study | PTC-44 |
| Plasmid to express 12VC1-cTEVP$\Delta$ | This study | PTC-45 |
| Plasmid to express 12VC1-cTEVP$\Delta$-I138T-S153N-T180A (uTEV3) | This study | PTC-46 |
| Plasmid to express CRAFx2-nTEVP-WT-IRES-BFP | This study | PTC-47 |
| Plasmid to express CRAFx3-nTEVP-WT-IRES-BFP | This study | PTC-48 |
| Plasmid to express 12VC1x2-nTEVP-WT-IRES-BFP | This study | PTC-49 |
| Plasmid to express 12VC1x3-nTEVP-WT-IRES-BFP | This study | PTC-50 |
| Plasmid to express pHRAS-WT-CRAF-nTEVP-WT-IRES-BFP | This study | PTC-51 |
| Plasmid to express pHRAS-G12V-CRAF-nTEVP-WT-IRES-BFP | This study | PTC-52 |
| Plasmid to express pHRAS-WT-12VC1-nTEVP-WT-IRES-BFP | This study | PTC-53 |
| Plasmid to express pHRAS-G12V-12VC1-nTEVP-WT-IRES-BFP | This study | PTC-54 |
| Plasmid to express CRAFx2-cTEVP-WT | This study | PTC-55 |
| Plasmid to express CRAFx3-cTEVP-WT | This study | PTC-56 |
| Plasmid to express 12VC1x2-cTEVP-WT | This study | PTC-57 |
| Plasmid to express 12VC1x3-cTEVP-WT | This study | PTC-58 |
| Plasmid to express pHRAS-WT-CRAF-cTEVP-WT | This study | PTC-59 |
| Plasmid to express pHRAS-G12V-CRAF-cTEVP-WT | This study | PTC-60 |
| Plasmid to express pHRAS-WT-12VC1-cTEVP-WT | This study | PTC-61 |
| Plasmid to express pHRAS-G12V-12VC1-cTEVP-WT | This study | PTC-62 |
| 12VC1-nTEVP-P2A-T2A-12VC1-cTEVP | This study | pAL0376 |
| 12VC1-nTEVP-tevs-GSGSG-tevs-12VC1-cTEVP | This study | pAL0377 |
| 12VC1-nTEVP-tevs-NZp-tevs-12VC1-cTEVP | This study | pAL0378 |
| IVT_deadTVMVP_T2A_halo | This study | pZAG0550 |
| IVT_TVMVP_T2A_halo | This study | pZAG0551 |
| IVT_SPOC_P3-AP4ms_T2A_AP3ms-P4_P2A_halo | This study | pZAG0553 |
| nTVMVP-37A-tevs-15B-tevs-cTVMVPm-T2A-nTVMVPm-tevs-15A-tevs-37B-cTVMVP | This study | pZAG0457 |
| IVT230_nTVMVP-P3-tevs-AP4ms-tevs-cTVMVPmut | This study | pZAG0468 |
| IVT230_nTVMVPmut-tevs-P3ms-tevs-AP4-cTVMVP | This study | pZAG0470 |
| IVT230_nTVMVP-P3-tevs-AP4ms-tevs-cTVMVPmut | This study | pZAG0468 |
| IVT230_nTVMVPmut-tevs-P3ms-tevs-AP4-cTVMVP-I40D | This study | pZAG0471 |
| IVT230_nTVMVP-P3-tevs-AP4ms-tevs-cTVMVPmut-I40D | This study | pZAG0469 |
| IVT230_nTVMVPmut-tevs-P3ms-tevs-AP4-cTVMVP | This study | pZAG0470 |
| IVT230_nTVMVP-P3-tevs-AP4ms-tevs-cTVMVPmut-I40D | This study | pZAG0469 |
| IVT230_nTVMVPmut-tevs-P3ms-tevs-AP4-cTVMVP-I40D | This study | pZAG0471 |
| P4_cTVMVP | This study | pZAG0504 |
| cTVMVPmut_AP4_tevs_P3_nTVMVP | This study | pZAG0505 |
| TVMVP_tevs_noninhibAI | This study | pZAG0148 |
| TVMVP --- tevs --- AI (v1.1) | This study | pZAG0149 |

| | | |
|---|---|---|
| TVMVP --- tevs --- AI with custom linker around tevs #1 (v1.2) | This study | pZAG0150 |
| TVMVP --- tevs --- AI with custom linker around tevs #2 (v1.3) | This study | pZAG0151 |
| circular_perm_C-TVMVP_tevs_AI_tevs_N-TVMVP_2-AAlinker | This study | pZAG0152 |
| circular_perm_C-TVMVP_tevs_AI_tevs_N-TVMVP_4-AAlinker | This study | pZAG0153 |
| circular_perm_C-TVMVP_tevs_AI_tevs_N-TVMVP_8-AAlinker | This study | pZAG0154 |
| cir-perm-AI-amp_CZp_20AAlinker_cTVMVP_tevs_AI_G_tevs_G_nTVMVP_20AAlinker_NZp | This study | pZAG0169 |
| cir-perm-AI-amp_CZp_20AAlinker_cTVMVP_tevs_AI_G_tevs_G_nTVMVP-delSKA_20AAlinker_NZp | This study | pZAG0170 |
| CZp_20linkr_cTVMVP_tevs_nTVMVP_20linkr_NZp | This study | pZAG0508 |
| CZp_20linkr_cTVMVP_tevs_nTVMVP_20linkr_NZp_dUP4 | This study | pZAG0510 |
| CZp_20linkr_cTVMVP_tevs_nTVMVP_20linkr_NZp_dDW4 | This study | pZAG0514 |
| CZp_20linkr_cTVMVP_tevs_37A_tevs_37B_tevs_nTVMVP_20linkr_NZp | This study | pZAG0526 |
| CZp_20linkr_cTVMVP_15A_tevs_15B___37B_tevs_37A_nTVMVP_20linkr_NZp | This study | pZAG0527 |
| CZp_20linkr_cTVMVP_37B_tevs_15B___15A_tevs_37A_nTVMVP_20linkr_NZp | This study | pZAG0528 |
| deadTEVP_mTagBFP2 | This study | pZAG0559 |
| TEVP_mTagBFP2 | This study | pZAG0560 |
